# Dihydroxy-Metabolites of Dihomo-gamma-linolenic Acid Drive Ferroptosis-Mediated Neurodegeneration

**DOI:** 10.1101/2023.01.05.522933

**Authors:** Morteza Sarparast, Elham Pourmand, Jennifer Hinman, Derek Vonarx, Tommy Reason, Fan Zhang, Shreya Paithankar, Bin Chen, Babak Borhan, Jennifer L. Watts, Jamie Alan, Kin Sing Stephen Lee

## Abstract

Even after decades of research, the mechanism of neurodegeneration remains understudied, hindering the discovery of effective treatments for neurodegenerative diseases. Recent reports suggest that ferroptosis could be a novel therapeutic target for neurodegenerative diseases. While polyunsaturated fatty acid (PUFA) plays an important role in neurodegeneration and ferroptosis, how PUFAs may trigger these processes remains largely unknown. PUFA metabolites from cytochrome P450 and epoxide hydrolase metabolic pathways may modulate neurodegeneration. Here, we test the hypothesis that specific PUFAs regulate neurodegeneration through the action of their downstream metabolites by affecting ferroptosis. We find that the PUFA, dihomo gamma linolenic acid (DGLA), specifically induces ferroptosis-mediated neurodegeneration in dopaminergic neurons. Using synthetic chemical probes, targeted metabolomics, and genetic mutants, we show that DGLA triggers neurodegeneration upon conversion to dihydroxyeicosadienoic acid through the action of CYP-EH, representing a new class of lipid metabolite that induces neurodegeneration via ferroptosis.

## Introduction

By 2050, the projected population older than age 65 is expected to be more than double, reaching over 1.5 billion, and the projected population older than 80 is predicted to triple to 426 million.^1^ As aging is a risk factor for neurodegeneration, it is expected that the population with dementia will significantly increase in the near future.^2^ However, the mechanisms of neurodegeneration remain unclear, and effective preventative measures and treatment are currently lacking.^3^ Therefore, identifying molecular mechanisms underlying neurodegeneration is an unmet medical need. While tauopathy, neuroinflammation, and excitotoxicity may play key roles in neurodegeneration, recent studies provide compelling evidence that ferroptosis could be a new mechanism underlying neurodegeneration.^3–5^ Ferroptosis is a non-apoptotic form of regulated cell death that is driven by an increase of iron-dependent lipid peroxidation in the cellular membrane.^4,6^ Epidemiological studies showed that patients with Parkinson’s disease (PD) or Alzheimer’s disease (AD) have elevated iron and lipid peroxide levels in the brain as compared to healthy controls, which is consistent with ferroptosis.^5,7–12^ The regulatory mechanism of ferroptosis in brain cells are understudied, although polyunsaturated fatty acids (PUFAs) play a critical role in this process.^13–17^

PUFAs are key structural components of plasma membranes and play a critical role in neuronal functions.^18^ Generally, ω-3 and ω-6 PUFAs are two of the major classes of PUFAs present in human diet.^19^ Human studies have demonstrated that an increase in plasma ω-3/ω-6 PUFA ratio decreases the risk of neurodegenerative diseases, including AD and PD.^20–23^ Nonetheless, even after decades of epidemiological studies in mammalian and cell-based models, how PUFAs affect neurodegeneration is poorly understood, with reported results that are contradictory.^21,24,25^ While most efforts in research have investigated the neuroprotective effects of ω-3 PUFA supplementation, few studies have examined the role of ω-6 PUFAs in neurodegeneration.^26–28^ This is surprising since the modern western diet has dramatically increased our consumption of ω-6 PUFAs.^29,30^ While the exact role of ω-6 PUFAs in neurodegenerative diseases is not understood, it is known that supplementing mammalian cells with ω-6 PUFAs sensitizes cells to ferroptosis.^13–15,31^ In addition, ω-6 dihomo-gamma-linolenic acid (20:3n-6, DGLA) induces ferroptosis in the *C. elegans* germline, while earlier studies have suggested that an epoxide metabolite of DGLA may mediate germ cell death.^17,32^

Although the mechanisms by which ω-6 PUFAs mediate biological effects remain undefined, recent studies have demonstrated that ω-6 PUFAs metabolites resulting from cytochrome P450 (CYP) enzymes and epoxide hydrolases (EHs) action are key signaling molecules for human physiology.^18,33,34^ As such, this study was initiated to test whether specific ω-6 PUFAs modulate neurodegeneration via their downstream CYP metabolites and to investigate if ferroptosis plays a role in the observed biology. Because the CYP enzymes and EHs are differentially expressed in tissues and cell types (Table S1),^35–39^ and the expression of both enzymes are significantly affected by cell passages,^40–42^ it is difficult to pinpoint a specific cell line suitable for our study. Therefore, a whole animal study is necessary to uncover this novel mechanism without worrying metabolites not being generated locally or overlooking critical cell-cell communications facilitated by these lipid metabolites.

To facilitate our study, we took an interdisciplinary approach by combining a simple genetic animal model, an inhibitor of a metabolic enzyme, synthesized lipid metabolites, and targeted metabolomics to systematically investigate the crosstalk between lipid metabolism, neurodegeneration, and ferroptosis. With this approach, we first demonstrate that among five tested PUFAs, only DGLA induces neurodegeneration in select neurons in *C. elegans*, with more pronounced effects in dopaminergic neurons, and to a lesser extent in glutaminergic neurons, with no observable effects in cholinergic and GABAergic neurons. Furthermore, we demonstrate that the DGLA-induced neurodegeneration is mediated through its downstream CYP-EH**s** metabolite, dihydroxyeicosadienoic acid (DHED), and ferroptosis is likely the mechanism involved in DHED-induced neurodegeneration.

## Results

### DGLA, but neither ω-3 nor other ω-6 PUFAs, induces degeneration specifically in dopaminergic neurons

Our prior lipidomic analysis shows that *C. elegans* absorbs exogenous PUFAs.^43,44^ To study the effect of dietary PUFAs on neuronal health span, we supplemented *Pdat-1::gfp* worms, in which the dopaminergic neurons are labeled by green fluorescent protein (GFP), with different ω-6 PUFAs and eicosapentaenoic acid (20:5n-5, EPA), the most abundant ω-3 PUFA in *C. elegans*,^45^ and tracked the dopaminergic neurons throughout the worm lifespan using fluorescent imaging (**Figure 1A-C**). Supplementation was done at the larvae stage 4 (L4) when *C. elegans* has a fully developed neuronal system thus enabling the investigation of neurodegeneration independent of neurodevelopment.^46^ Among the tested PUFAs, only DGLA induced significant degeneration in the dopaminergic neurons (**Figure 1B**). Furthermore, DGLA triggered degeneration in dopaminergic neurons in a dose-dependent manner with an EC_50_= 51.4 μM and 31.2 μM at day 1 and day 8 adulthood, respectively (**Figure 1D, and S1**). We also showed that the vehicle control, ethanol, did not change dopaminergic neurons’ health span as compared to the control (**Figure S2**). In addition, we found that different types of dopaminergic neurons in the hermaphrodite had varying sensitivities to treatment with DGLA, with the ADE neurons (ADE>>>CEP>PDE) being the most impacted (**Figure S3**). Moreover, loss of the GFP signal did not appear to result from transcriptional repression of the *Pdat-1::GFP* transgene induced by treatment with DGLA, since a similar trend was observed with the *Pcat-2::GFP* transgenic line upon treatment with DGLA (**Figure 1E and F**)). We then examined whether DGLA can induce degeneration in other major types of neurons that play key roles in neurodegenerative diseases including GABAergic, glutaminergic, and cholinergic neurons. Significant neurodegeneration was not observed in GABAergic (*Punc-25::gfp*) and cholinergic neurons (*Punc-17::gfp*) worms supplemented with DGLA (**Figure 1G-J**).

**Figure 1.**
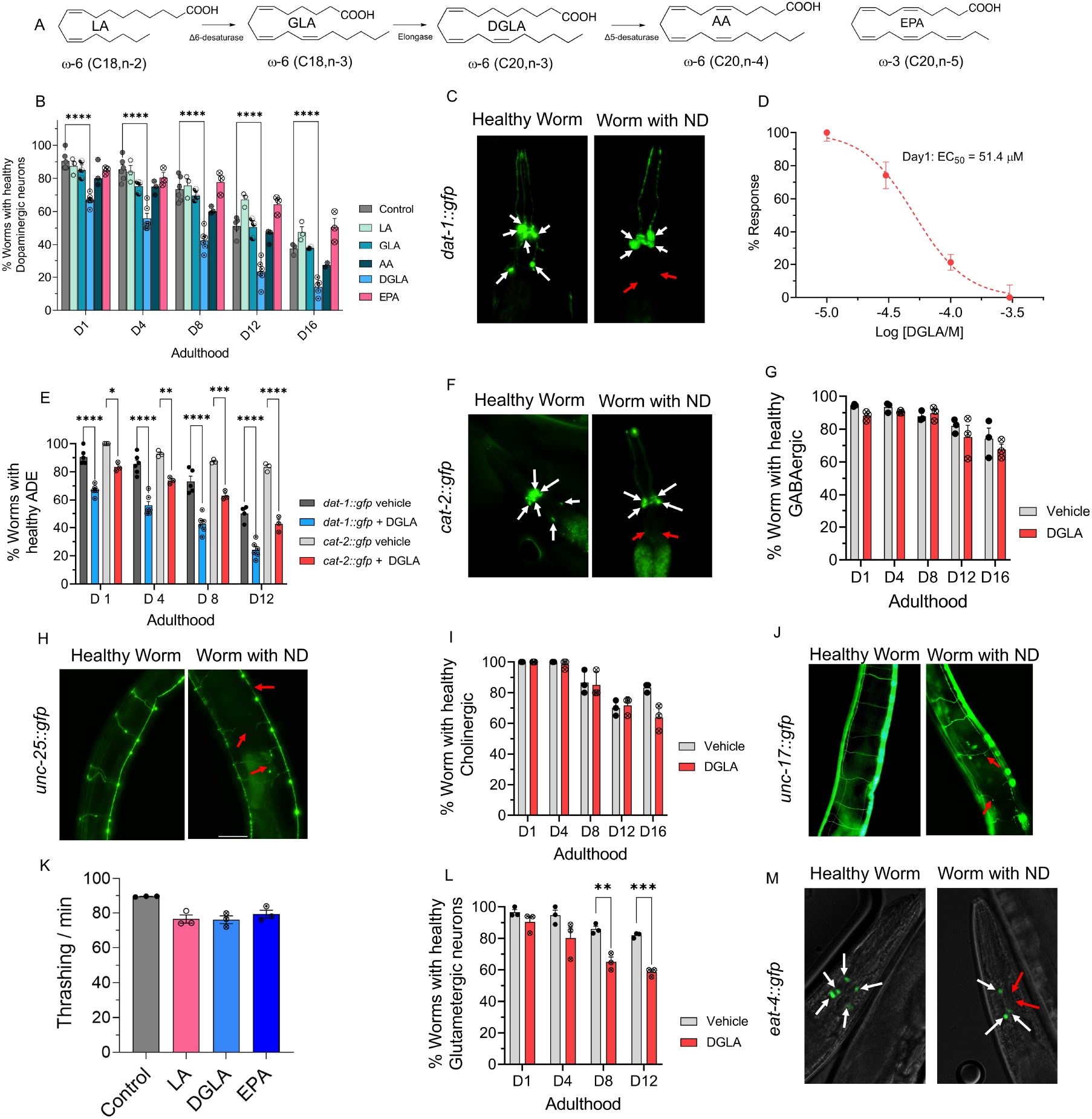
DGLA, but not other ω-3 and ω-6 PUFAs, induces degeneration, specifically in dopaminergic neurons. **(A**) Structure of different ω-6 and ω-3 PUFAs examined in this study; (**B**) Percentage (%) of worms with healthy dopaminergic neurons for *Pdat-1::gfp* with and without supplementation with 100 μM of different ω-6 and ω-3 PUFAs. (**C**) Fluorescent images of *Pdat-1::gfp* worms with healthy and degenerated dopaminergic neurons (White arrows represent healthy neurons, and red arrows show degenerated/disappeared neurons); (**D**) Dose response curve: the effect of different DGLA concentrations on degeneration of ADE neurons on Day 1 adulthood; **(E)** Comparing the ADE neurons degeneration in *Pdat-1::gfp* and *Pcat-2::gfp* supplemented with 100 μM DGLA; **(F)** Fluorescent images of *Pcat-2::gfp* worms with healthy and degenerated dopaminergic neurons (White arrows represents healthy neurons, and red arrow shows degeneration/disappearance neurons); (**G**) Percentage (%) of worms with healthy GABAergic neurons for *Punc-25::gfp* with and without supplementation with 100 μM DGLA; (**H**) Fluorescence images of *Punc-25::gfp* worm with healthy and degenerated GABAergic neurons (Red arrows show different signs of neurodegeneration including ventral cord break, commissure break and branches); (**I**) Percentage (%) of worms with healthy cholinergic neurons for *Punc-17::gfp* with and without supplementation with 100 μM DGLA; (**J**) Fluorescence Images of *Punc-17::gfp* worms with healthy and degenerated cholinergic neurons (Red arrows show different signs of neurodegeneration including ventral cord break, commissure break and branches); (**K**)Thrashing on Day 8 adulthood of wild-type raised on 100 μM of LA, DGLA, and EPA; (**L**) Percentage (%) of worms with healthy glutamatergic neurons with *Peat-4::gfp* with and without supplementation with 100 μM DGLA; (**M**) Fluorescent images of *Peat-4::gfp* worms with healthy and degenerated glutamatergic neurons (White arrows represent healthy neurons, and red arrows show degenerated/disappeared neurons). All supplementations were 6 done at the L4 stage. For all experiments N=3, and about 20 worms were tested on each trial. Two-way analysis of variance (ANOVA), Tukey’s multiple comparison test for (B) and (D); and t test for K: *P ≤ 0.05, **P ≤ 0.01, ***P ≤ 0.001, ****P < 0.0001, non-significant is not shown.

These findings were further confirmed with a lack of significant changes in thrashing assays in *C. elegans* treated with DGLA (**Figure 1K**), which requires cholinergic and GABAergic neurons activity.^47,48^ In the case of glutamatergic neurons (*Peat-4::gfp*), treatment with DGLA caused cell loss in glutamatergic neurons later in the *C. elegans* lifespan compared to the dopaminergic neurons (**Figure 1L and M**). Altogether, our results suggest that the effect of PUFAs on neurodegeneration is structurally specific. In addition, in contrast to conclusions reported previously,^13,14,49^ our data showed that treatment of the more peroxidizable arachidonic acid and EPA did not trigger neurodegeneration. Furthermore, our results indicate that the effect of DGLA on neurodegeneration is neuron-type selective, warranting future studies that may shine light on the molecular mechanism(s).

The remaining studies focused on the degeneration of dopaminergic neurons, as was found they are most sensitive to DGLA treatment. Because more robust data were obtained with transgenic *C. elegans Pdat1::gfp*, the rest of our studies were conducted using this strain. Most of the experiments were performed using day 1 and day 8 adults, enabling the determination of acute and chronic effects of DGLA treatment on neurodegeneration. Day 8 worms resemble a middle-aged population of *C. elegans*, thus the effect of DGLA treatment on age-associated neurodegeneration can also be investigated without a significant loss (death) of the tested population, facilitating the throughput of our studies.

### DGLA induces neurodegeneration in dopaminergic neurons through ferroptosis

Recent studies show that treatment with DGLA can induce ferroptosis in germ cells and cause sterility in *C. elegans*.^17^ To test whether treatment with DGLA induces degeneration in dopaminergic neurons through ferroptosis, *Pdat-1::gfp* expressing worms were co-treated with DGLA and liproxstatin-1 (Lip-1), a radical-trapping antioxidant and ferroptosis inhibitor.^50^ While C. *elegans* treated with Lip-1 alone showed no significant effect on age-associated degeneration of dopaminergic neurons as compared to the vehicle control, co-treatment of DGLA with Lip-1 fully rescued the neurodegeneration triggered by DGLA in day 1 adults and largely rescued DGLA-induced neurodegeneration in day 8 adults (**Figure 2A**). Encouraged by these results, we examined neurodegeneration caused by ferroptosis in DGLA-treated worms using pharmacological and genetic approaches. An increase in the labile iron (II) pool and membrane lipid peroxidation are molecular hallmarks of ferroptosis.^6,51^ Therefore, we tested whether treatment with Trolox, a water-soluble form of vitamin E and lipid peroxidation inhibitor, and 2,2’-bipyridine, an iron (II) chelator, alleviates DGLA-induced neurodegeneration. Co-treatment with either Trolox or 2,2’-bipyridine rescued DGLA-induced neurodegeneration, suggesting that both the labile iron (II) pool and membrane lipid peroxidation are involved in DGLA-induced neurodegeneration (**Figure 2B and C**). To specifically investigate whether ferroptosis is involved in DGLA-induced neurodegeneration, was also pursued a genetic approach.

**Figure 2.**
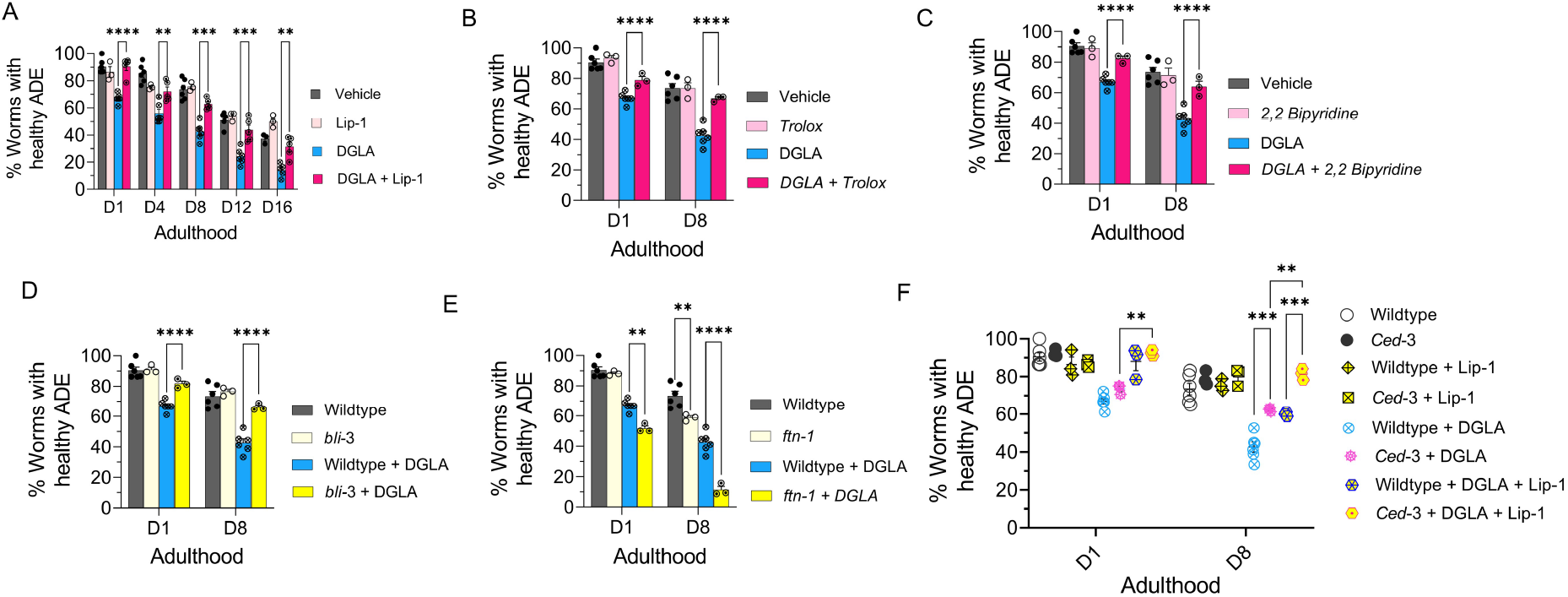
DGLA Induces neurodegeneration in dopaminergic neurons via ferroptosis. (**A**) Percentage (%) of worms with healthy ADE neurons of worms exposed to 100 μM DGLA ± 250 μM Liproxstatin-1; (**B**) Percentage (%) of worms with healthy ADE neurons in wild-type *C. elegans* treated with 100 μM DGLA ± 500 μM Trolox (vitamin E); (**C**) Percentage (%) of worms with healthy ADE neurons for *Pdat-1::*gfp worms treated with 100 μM DGLA± 100μM 2,2’-bipyridine; (**D**) Percentage (%) of worms with healthy ADE neurons in *Pdat-1::gfp* and *Pdat-1::gfp;bli-3* worms treated with 100 μM DGLA; (**E**) Percentage (%) of worms with healthy ADE neurons for *Pdat-1::gfp* and *Pdat-1::gfp; ftn-1* worms treated with 100 μM DGLA; (**F**)Percentage (%) of worms with healthy ADE neurons with *Pdat-1::gfp* and *Pdat-1::gfp; ced-3* worms treated with 100 μM DGLA ± 250 μM Liproxstatin-1. All supplementations were done at the L4 stage. A two-way analysis of variance (ANOVA), Tukey’s multiple comparison test. *P ≤ 0.05, **P ≤ 0.01, ***P ≤ 0.001, ****P < 0.0001, NS, not significant. DGLA: Dihomo-γ-linolenic acid, LA: Linoleic acid, EPA: eicosapentaenoic acid, Lip-1: Liproxstatin-1.

Previous studies have shown that nicotinamide adenine dinucleotide phosphate (NADPH) oxidase (NOX) family of superoxide-producing enzymes (NOX/DUOX) plays a critical role in ferroptosis in mammals and can exacerbate dopaminergic neurotoxicity triggered by ferroptosis inducers.^4,52^ In addition, ferritin (FTN) is also a key ferroptosis regulatory protein, and genetic knockout of FTN has been shown to sensitize *C. elegans* to ferroptosis.^17,53^ To further test whether ferroptosis is involved in DGLA-induced neurodegeneration, two new transgenic *C. elegans* strains were created by crossing the *Pdat-1::gfp* with transgenic strains that carry either a loss of function of *bli-3* (*C. elegans* homolog of NOX) mutant or genetic knockout of *ftn-1* (**Figure 2D, 2E, and S4**). Our results indicated that the loss of function *bli-3* mutant reduced the degeneration of dopaminergic neurons triggered by DGLA (**Figure 2D**). Worms with loss of function mutations of BLI-3 attenuated ability to generate reactive oxygen species, thus minimizing lipid peroxidation, and as a result, reduced ferroptosis.^17,54^ This result further confirms the pharmacological observation after supplementing worms with the lipophilic antioxidant vitamin E (Trolox), which led to the suppression of neurodegeneration in DGLA-treated worms (**Figure 2B**). Furthermore, genetic knockout of *ftn-1* enhanced DGLA-induced neurodegeneration, suggesting that DGLA requires the labile iron (II) pool to exert its effect on dopaminergic neurons (**Figure 2E**). Our data strongly suggest that DGLA causes degeneration of dopaminergic neurons at least partly through ferroptosis. As illustrated in **Figure 2A**, while Lip-1 fully rescued DGLA-induced neurodegeneration for day 1 adults, such rescuing effect diminished as *C. elegans* aged. Furthermore, the EC_50_ of DGLA in triggering neurodegeneration for day 1 and day 8 adults is different. Therefore, we hypothesize that chronic treatment of DGLA induces other programmed cell-death pathways, like apoptosis, leading to neurodegeneration.

To test whether DGLA also induces neurodegeneration through apoptosis,^55^ an additional transgenic strain was developed. The CED-3 protein, a key enzyme involved in apoptosis, was genetically knocked out in worms in which the dopaminergic neurons were labeled by GFP^17,56^ to create transgenic line, *Pdat-1::gfp;ced-3(n717)*. Interestingly, while no significant difference was observed between *Pdat-1::gfp;ced-3(n717)* and *Pdat-1::gfp* worms treated with DGLA at day 1 adulthood, worms that had the *ced-3* genetic knockout demonstrated partial rescue from dopaminergic neuron degeneration induced by DGLA (**Figure 2F**) at day 8 of adulthood. Furthermore, the *ced-3* knockout worms at day 8 adulthood that were co-treated with Lip-1 were fully rescued from dopaminergic neurodegeneration induced by DGLA (**Figure 2F**). These results could explain the differences observed for day 1 and day 8 worms treated with Lip-1 and DGLA (**Figure 2A**), as well as the differences in the EC_50_ of DGLA-induced neurodegeneration between day 1 adults (EC50 = 51.4 μM) and day 8 adults (EC_50_ = 31.2 μM) (**Figure 1D**). The lower EC_50_ for DGLA-induced neurodegeneration for day 8 adults suggests that other neurodegenerative mechanisms (i.e., apoptosis, autophagy) are involved and could either synergize or provide an additive effect with DGLA-induced neurodegeneration. Together, these results suggest that dietary DGLA induces neurodegeneration via ferroptosis in early adulthood and both ferroptosis and additional mechanism(s), such as apoptosis, are induced by DGLA in dopaminergic neurodegeneration in middle-aged *C. elegans*.

### Downstream metabolites of DGLA are key players in neurodegeneration induced by DGLA treatment

In mammals, DGLA and other PUFAs are mono-oxygenated by cytochrome P450 enzymes (CYPs) to hydroxy- and epoxy-PUFAs. Epoxy-PUFAs are further hydrolyzed by epoxide hydrolases (EHs) to the dihydroxy-PUFAs (**Figure 3A**).^18^ Numerous animal and human studies have demonstrated that endogenous levels of CYP and EH metabolites produced from various PUFAs are highly correlated to dietary intake of the corresponding PUFAs,^57–59^ in stark contrast to metabolites generated by cyclooxygenases and lipoxygenases, which are less correlated.^60–62^ Both epoxy- and dihydroxy-PUFAs are key signaling molecules for mammalian physiology including, but not limited to, neuroprotection.^18,63,64^ Therefore, we hypothesized that DGLA primarily induces ferroptosis-mediated neurodegeneration via its CYP and/or EH metabolites, a previously unexplored area. To test this hypothesis, we first investigated whether the CYP/EH metabolism is involved in DGLA-induced ferroptosis-mediated neurodegeneration by investigating how treatment with DGLA impacts CYP/EH metabolism. Our results indicated that treatment with 100 μM DGLA increased the whole animal endogenous levels of the corresponding epoxyeicosadienoic acid (EED) and dihydroxyeicosadienoic acid (DHED) to ∼200 pmol/g and ∼800 pmol/g, respectively (regioisomer-dependent **Figure S5**), using our oxylipin analysis (Pourmand *et. al*, unpublished, see experimental methods in SI). These results were similar to the endogenous levels of EPA CYP/EH metabolites, epoxyeicosatetraenoic acid (EpETEs) and dihydroxyeicosatetraenoic acids (DHETE) which are 50-919 pmol/g and 0-458 pmol/g, respectively (regioisomer-dependent) in intact *C. elegans*, suggesting that the increased level of EED and DHED is physiologically relevant (**Figures 3B and S5**). Therefore, we sought to determine whether these downstream metabolites (EED and DHED) are key mediators for neurodegeneration induced by treatment with DGLA. Transgenic *C. elegans* (*Pdat-1::gfp*) were co-treated with DGLA and 12-(1-adamantane-1-yl-ureido-) dodecanoic acid (AUDA, 100 μM), an EH inhibitor with selective action to inhibit the function of CEEH1 and CEEH2 (*C. elegans* EH1 and EH2 isoforms).^65^ AUDA treatment increased the level of EED and decreased the DHED *in vivo* concentration, and fully rescued dopaminergic neurodegeneration induced by DGLA, suggesting that the CYP/EH-derived downstream metabolites of DGLA play a critical role in DGLA-induced neurodegeneration (**Figures 3B, C**).

**Figure 3.**
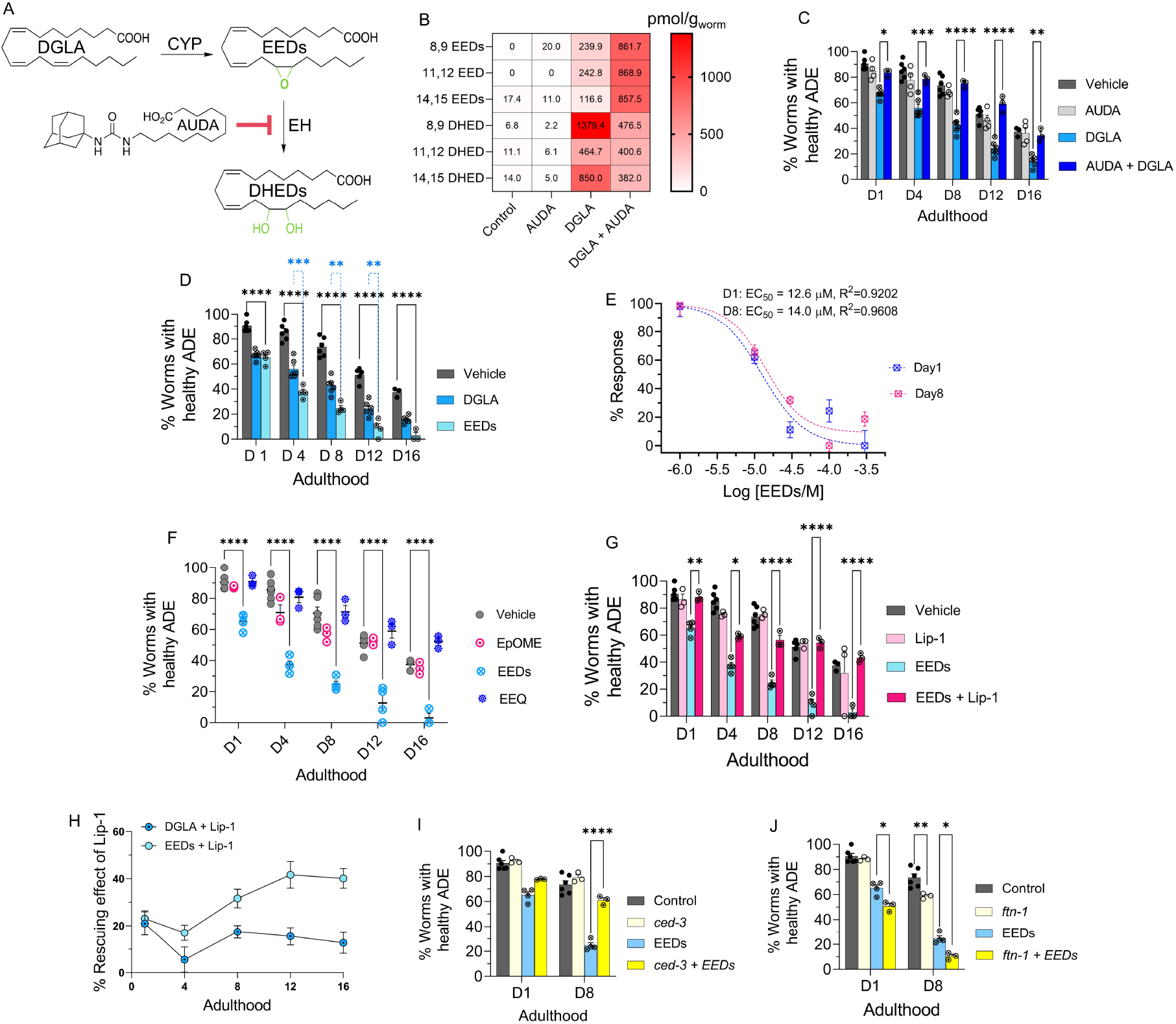
EED, epoxy metabolites downstream of DGLA, induce neurodegeneration by ferroptosis. (**A**) DGLA is metabolized to EED and DHED through the CYP and epoxide hydrolase enzymes, respectively; and AUDA inhibits epoxide hydrolase; (**B**) Oxylipin profile representing the pmol/g of EED and DHED regioisomers in worms treated with 100 μM of DGLA ± 100 μM AUDA compared to control (**C**) Percentage (%) of worms with healthy ADE neurons for *Pdat-1::gfp* treated with 100 μM of DGLA ± 100 μM AUDA; (**D**) Percentage (%) of worms with healthy ADE neurons for *Pdat-1::gfp* worms treated with 100 μM DGLA and 100 μM EED; (**E**) Dose response curve: the effect of different concentration of EED on degeneration of ADE neurons on Day 1 and Day 8 adulthood; (**F**) Percentage (%) of worms with healthy ADE neurons in *Pdat-1::gfp* worms treated with 100 μM of different Ep-PUFAs, EpOME, and EEQ; (**G**) Percentage (%) of worms with healthy ADE neurons of worms treated with 100 μM DGLA ± 100 μM Liproxstatin-1;(**H**) Comparison the effect of 250 μM Liproxstatin-1 on *Pdat-1::gfp* worms treated with 100 μM of DGLA compared to 100 μM EED;(**I**) Percentage (%) of worms with healthy ADE neurons with *Pdat-1::gfp* and *Pdat-1::gfp;ced-3* worms treated with 100 μM; (**J**) Percentage (%) of worms with healthy ADE neurons for *Pdat-1::gfp* and *Pdat-1::gfp; ftn-1* worms treated with 100 μM DGLA; All supplementations were done at the L4 stage. Two-way analysis of variance (ANOVA), Tukey’s multiple comparison test. *P ≤ 0.05, **P ≤ 0.01, ***P ≤ 0.001, ****P < 0.0001, without * not significant. DGLA: Dihomo-γ-linolenic acid, EED: epoxyeicosadienoic acids, DHED: dihydroxyeicosadienoic acids, CYP: Cytochrome P450, EH: epoxide hydrolase, AUDA: 12-(1-adamantane-1-yl-ureido-) dodecanoic acid, Lip-1: Liproxstatin-1, EpOME: epoxyoctadecenoic acids; EEQ: epoxyeicosatetraenoic acid.

Nonetheless, these results do not discriminate between AUDA’s ability to stabilize the level of EED *in vivo* for the observed rescue, or to block the production of DHED metabolites that result from inhibiting *C. elegans* EHs (**Figures 3B** and **S6**). To distinguish between the latter two possibilities, we synthesized both EED and DHED and tested their effects in *C. elegans* following the procedures in previous reports.^66–69^ Treatment with 100 μM EED at the L4 stage induced a more severe neurodegenerative phenotype than treatment with DGLA at the same concentration in the dopaminergic neurons in all tested ages (**Figure 3D)**, with a much lower EC_50_ (12.6 μM vs 51.4 μM) as compared to DGLA on day 1 adult **(Figures 3E and 1D)**. The same trend was observed in glutamatergic neurons when comparing treatments of EED and DGLA, and similar to DGLA treatment, no significant neurodegeneration was observed in GABAergic and cholinergic neurons after treatment with EED (**Figure S7**).

To test whether the effect of EED is structurally specific, C18:1 epoxyoctadecenoic acid (EpOME), an epoxy-metabolite of LA, and a more peroxidizable C20:4 epoxyeicosatetraenoic acid (EEQ), an epoxy-metabolite of EPA, were examined (**Figure 3F**) and had no effects on neurodegeneration. These results indicate that the effect of epoxy-PUFAs on neurodegeneration is specific to EED, but not other epoxy-PUFAs. Similar to the neurodegeneration induced by DGLA, co-treatment with Lip-1 rescued neurodegeneration caused by EED, and Lip-1 was more effective in alleviating EED induced neurodegeneration compared to DGLA-induced neurodegeneration **(Figures 3G, H)**. Furthermore, like DGLA, neuronal degeneration induced by EED was not rescued by a genetic knockout of *ced-3*. Genetic knockout of *ftn-1* escalates the effect of EED in both day 1 and day 8 adult, again suggesting that ferroptosis plays a critical role in EED-induced neurodegeneration **(Figures 3I and J)**.

Our results further suggested that DGLA metabolites are lipid mediators responsible for the effect of DGLA on neurodegeneration. Inhibition of EED hydrolysis using an EH inhibitor (AUDA) resulted in rescue of EED-induced neurodegeneration in *C. elegans* (**Figure 4A**). The oxylipin profile of worms co-treated with EED and AUDA at 100 μM shows that blocking the metabolism of epoxy-PUFAs to dihydroxy-PUFAs, specifically EED to DHED with AUDA, stabilizes endogenous levels of epoxy-PUFAs including EED, and decreases the *in vivo* levels of dihydroxy-PUFAs and DHED (**Figures 4B and S8**). Altogether, our data further suggest that specific DGLA downstream metabolites, either EED or DHED, are responsible for DGLA-mediated neurodegeneration.

**Figure 4.**
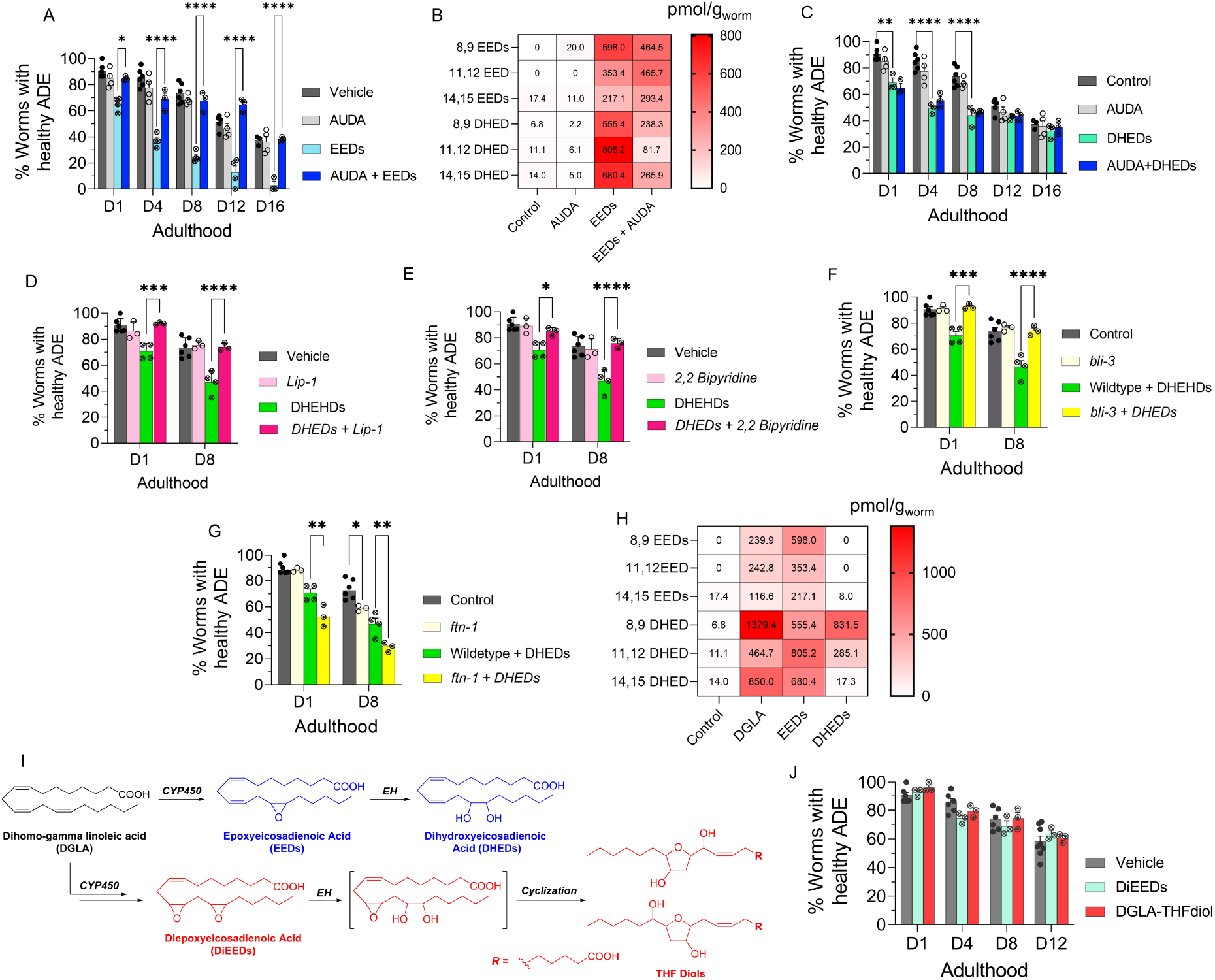
DHEDs, dihydroxy fatty acids downstream of DGLA/EED, are key candidates for neurodegeneration induced by DGLA in dopaminergic neurons. (**A**) Percentage (%) of worms with healthy ADE neurons in *Pdat-1::gfp* worms treated with 100 μM of EED ± 100 μM AUDA; (**B**) Oxylipin profile representing pmol/g of EED and DHED regioisomers in worms treated with 100 μM of EED ± 100 μM AUDA compared to control. (**C**) Percentage (%) of worms with healthy ADE neurons for *Pdat-1::gfp* treated with 100 μM of DHED ± 100 μM AUDA; (**D**) Percentage (%) of worms with healthy ADE neurons of worms exposed to 100 μM DHED ± 250 μM Liproxstatin-1; (**E**) Percentage (%) of worms with healthy ADE neurons for *Pdat-1::*gfp worms treated with 100 μM DHEDs ± 100 μM 2,2’-bipyridine; (**F**) Percentage (%) of worms with healthy ADE neurons for *Pdat-1::gfp* and *Pdat-1::gfp;ftn-1* worms treated with 100 μM DHED; (**G**) Percentage (%) of worms with healthy ADE neurons in *Pdat-1::gfp* and *Pdat-1::gfp;bli-3* worms treated with 100 μM DEHDs; (**H**) Oxylipin profile representing the pmol/g of EED and DHED regioisomers in worms treated with 100 μM of DGLA, EED, and DHED compared to control. (**I**) two possible metabolisms of DGLA through the CYP/EH pathways; the alternative metabolism is that CYP can do two consecutive oxidations (or under oxidative stress) to yield diepoxies EED, which then EH will open one epoxide which under physiological conditions can cyclize to THF diols. (**J**) Percentage (%) of worms with healthy ADE neurons for *Pdat-1::gfp* treated with 100 μM of DiEED and 100 μM DGLA-THFdiol. All supplementations were done at the L4 stage. Two-way analysis of variance (ANOVA), Tukey’s multiple comparison test. *P ≤ 0.05, **P ≤ 0.01, ***P ≤ 0.001, ****P < 0.0001, without * not significant. DGLA: Dihomo-γ-linolenic acid, EED: epoxyeicosadienoic acid, DHED: dihydroxyeicosadienoic acid, CYP: Cytochrome P450, EH: epoxide hydrolase, AUDA: 12-(1-adamantane-1-yl-ureido-) dodecanoic acid, DiEED: diepoxyeicosadienoic acid.

We further corroborated our hypothesis by supplementing *Pdat-1::gfp* worms with 100 μM of DHED, which showed significant neurodegeneration compared to the vehicle control (**Figure 4C**). Intriguingly, co-treatment with AUDA and DHED did not alleviate neurodegeneration induced by DHED, further confirming that DHED is likely the main driver of dopaminergic neurodegeneration in our model (**Figure 4C**). Co-treatment with AUDA alleviated DGLA-induced neurodegeneration likely by blocking the formation of DHED. In addition, co-treatment with Lip-1 and 2,2-bipyridine rescued the neurodegeneration caused by DHED **(Figures 4D, and E)**. Furthermore, the loss of function *bli-3* mutant also reduced the degeneration of dopaminergic neurons triggered by DHED, and genetic knockout of *ftn-1* augments DHED-induced neurodegeneration **(Figures 4F, and G)**. Together, these results suggest that the labile iron (II) pool and subsequently ferroptosis are involved in the effect of DHED on dopaminergic neurons. It is noteworthy that we did not observe more severe neurodegeneration induced by DHED as compared to EED supplementation. This effect may be due to the difference in lipid transport mechanism between dihydroxy-PUFAs, PUFAs and Ep-PUFAs, as suggested by a previous study.^70^ DHED is not absorbed as well as EED, and thus are not as potent at the same concentration. This hypothesis was confirmed by oxylipin profiling, which showed significantly lower DHED levels (especially for 11,12 and 14,15 DHED) in worms treated with 100 μM DHED as compared to those treated with 100 μM EED or 100 μM DGLA (**Figures 4D** and **S9**). While DGLA and EED exhibit a continuous increase in dopaminergic neurodegeneration over their lifespan, DHED exhibited a plateau after day 8, which suggests the presence of a possible mechanism for removal of the offending agent, either by inducing downstream metabolism or activating lipid transport of DHED after chronic treatment. Our data strongly suggest that DGLA/EED induces neurodegeneration through their downstream metabolites. Beyond DHED, there have been a few reports of other downstream CYP-mediated metabolites, namely epoxy-hydroxy-PUFA and diepoxy-PUFAs, which ultimately undergo spontaneous intramolecular cyclization in physiological conditions to form an understudied class of metabolites, the tetrahydrofuran-diols (THF-diols). These THF-diols could be a new class of lipid mediators in mammals.^71,72^ To examine these potential mediators of biological activity, isomeric vicinal diepoxyeicosenoic acid (DiEEe) and its corresponding THF-diol (DGLA-THF-diol) were synthesized and incubated with the worms (see the experimental section in SI and **Figure 4E**). However, no significant loss was observed in dopaminergic neurons in worms treated with 100 μM DiEEe and its corresponding THF-diol compared to the vehicle control (**Figure 4F)**. These results strongly suggest that DHED constitutes a novel class of lipid mediators that induces neurodegeneration largely mediated by ferroptosis. In addition, EpETE and EpOME produced from EPA and LA, respectively, showed no effect on neurodegeneration, which further corroborates the effect of DHED on ferroptosis-mediated neurodegeneration is structurally selective. Furthermore, the results obtained from the treatment with more peroxidizable EPA and EpETE indicate that the effect of DHED is not due to an increase in peroxidation of the cell membrane, a known mechanism that sensitizes ferroptosis.^15,73–75^ As a whole, the results summarized above are contrast with reports on the effect of PUFAs^76–78^ or their metabolites, such as lipoxygenase’s metabolites,^50,73,77^ ether lipids,^79^ etc. on ferroptosis.

## Discussion

Our studies revealed that DHED, CYP-EH metabolite of DGLA, is a novel class of lipid molecules that triggers ferroptosis-mediated degeneration in select neuron types in *C. elegans*. Our study addresses critical gaps in knowledge in the field of lipid pharmacology, neurodegeneration, and ferroptosis, including how ω-6 PUFAs may trigger neurodegeneration and the identity of endogenous signaling molecules that induces ferroptosis-mediated neuronal cell death. Most research investigates the beneficial effects of ω-3 PUFA supplementation in neurodegenerative diseases, with contradictory findings.^26,27,30^ Few studies have tested the effect of ω-6 PUFAs on neurodegeneration.^28^ This is of high interest since ω-6 PUFA levels are typically high in western diets. Our findings in *C. elegans* demonstrate that, unlike other PUFAs, the ω-6 DGLA induces ferroptosis-mediated degeneration specifically in dopaminergic and to a lesser extent in glutaminergic neurons. Recent reports suggest PUFAs play a critical role in ferroptosis and treating cells with PUFAs, their ether-lipid metabolites, and lipoxygenase metabolites, hydroperoxyeicotetraenoic acids, sensitize cells to ferroptosis, but not induce ferroptosis themselves.^50,79^ However, specific endogenous mediators that regulate the upstream pathway of ferroptosis remain unknown, although synthetic compounds such as erastin, RSL-3, and natural products like α-eleostearic acid have been identified as agents that can induce ferroptosis.^13–15,31^ Our results indicate that DGLA induces ferroptosis-mediated neuronal death likely through its downstream endogenous CYP-EH metabolites, DHED, and EH plays a critical role in modulating DGLA-mediated ferroptosis. Our study complements the elegant work showing that DGLA induces ferroptosis in germline and cancer cells.^17^ The identification of potential lipid signaling molecules represents a critical first step to investigating the molecular mechanism behind the effects of PUFAs on ferroptosis-mediated neurodegeneration.

Recent reports demonstrated that the expression of soluble EH, an human ortholog of CEEH 1/2, is upregulated in patients with neurodegenerative diseases including Parkinson’s disease and Alzheimer’s disease and, inhibition of soluble EH is beneficial for neurodegeneration in multiple neurodegenerative diseases animal models.^18,80–83^ While the specific role of soluble EH in neurodegeneration is largely unknown, these studies suggested that the epoxy-fatty acids, the substrates of soluble EH, are neuroprotective, and the corresponding downstream EH metabolites dihydroxy-fatty acids have no effect, although a few studies in cell and animal models have showed that these dihydroxy-fatty acids can have detrimental or toxic effects on cells.^84,85^ Our results provide an alternate perspective of how neurodegeneration could be regulated endogenously by modulating EH activity to increase ferroptotic metabolites, namely DHED, which has seldom been studied.

In this study, we employed an approach, which comprised of a simple animal model, an inhibitor of a metabolic enzyme, synthesized lipid metabolites, and targeted metabolomics to systematically investigate the crosstalk between lipid metabolism, neurodegeneration, and ferroptosis in a highly efficient way. We have not only identified the key mediator for ferroptosis-mediated neurodegeneration but have also revealed that DGLA and its metabolites have more pronounced effects on dopaminergic neurons, mild effects on glutaminergic neurons, and no effects on cholinergic and GABAergic neurons in *C. elegans*. Our results complement previous studies by Zille. *et al*., which showed that different cell types could have distinct regulatory pathways for ferroptosis.^86^ While the specific mechanism behind why DGLA and its metabolites, DHED, are more detrimental to dopaminergic neurons remains unknown, Solano Fonseca *et al*. reported a similar vulnerability of different neuron-types in response to biomechanical injury and suggested that such observation could be due to different physiological regulatory mechanisms between different neuron types.^87^ Such neuron type-specific effects triggered by DGLA and DHED warrant future investigation to uncover potential new neurodegeneration mechanisms. Investigating ferroptosis at the systems level to understand differential ferroptosis mechanisms between tissues is challenging, owing to the lack of appropriate genetic and imaging tools. The genetic malleability of *C. elegans* provides a suitable platform for the study of ferroptosis in a tissue-specific manner. Furthermore, as illustrated in Table S1, most cell lines do not express soluble EH, a human ortholog of CEEHs and studies have demonstrated that different tissues express CYP enzymes and soluble epoxide hydrolase differently.^35–38^ Therefore, a whole animal approach is more appropriate for us to explore this novel mechanism and *C. elegans* provides a simple animal model. In addition, the adaptability to high-throughput studies of *C. elegans* allows us quickly to dissect complicated pathways. As such, it is possible to explore how ferroptosis can be regulated differentially by endogenous signaling molecules, such as DHED, between cell-types in an intact organism. Furthermore, the chemical tools developed and utilized for this study lead to the exploration of novel hypotheses that aim to unravel PUFAs effects on organismal physiology, an area that is not only understudied, but also is challenging to execute in mammalian models and humans.

## Conclusion

Oxidized lipid metabolites are key mediators for organismal physiology. Ferroptosis, characterized by an increase of iron-dependent lipid peroxidation, could be a novel mechanism for neurodegeneration. In this study, we reported that exogenous DGLA triggers neurodegeneration predominantly in dopaminergic neurons via its downstream cytochrome P450-epoxide hydrolase (CYP-EH) metabolite, dihydroxyeicosadienoic acid (DHED). The observed neurodegeneration induced by DGLA/DHED is likely mediated by ferroptosis at the early stages and a combination of ferroptosis and apoptosis after chronic treatment with DGLA/DHED. This study revealed that CYP-EH polyunsaturated fatty acid (PUFA) metabolism is one of the key intrinsic regulatory mechanisms of ferroptosis-mediated neurodegeneration, and EH could be a novel target for ferroptosis-mediated diseases.

## Acknowledgments

Some nematode strains were provided by the Caenorhabditis Genetics Center, funded by the NIH Office of Research Infrastructure Programs (P40 OD010440). The EM641 *Pcat-2::gfp* strain was a gift from Dr. Scott Emmons. The GA912 *ftn-1(ok3625)* strain was a gift from Dr. David Gems (University College London, London, UK). Funding to K.S.S.L. was provided by the NIGMS R35 GM146983, and Pearl Aldrich Endowment for aging research. Funding to J.A. was provided by the NIH (R03 AG075465). J.L.W. was supported by NIH R01 GM133883. K.S.S.L. and J. A. were partially supported by startup funding from Michigan State University. M.S. was partially supported by Pearl Aldrich Endowment for aging research. Funding to T.R was graciously provided by Integrative Pharmacological Sciences Training Grant NIH T32GM142521. We acknowledge the MSU RTSF Mass Spectrometry and Metabolomics Core Facilities for support with oxylipin and lipidomic analysis. We would like to thank Dr. Brian Ackley (University of Kansas) for his advice on the neuronal studies; Dr. Scott Dixon for his comments for this study, as well as Mr. Devon Dattmore, Ms. Leslie Ramirez, and Ms. Heather deFeijter-Rupp (Michigan State University) for their help and assistance.

## Supporting Information

The Supporting Information including experimental methods and materials, supplemental figures and table, synthesis and characterization of DGLA metabolites.

