## supplemental information for "Dihydroxy-Metabolites of Dihomo-gamma-linolenic Acid Drive Ferroptosis-Mediated Neurodegeneration"

1. **Supporting Figures and Tables (Figure S1-S8, and Table S1):**

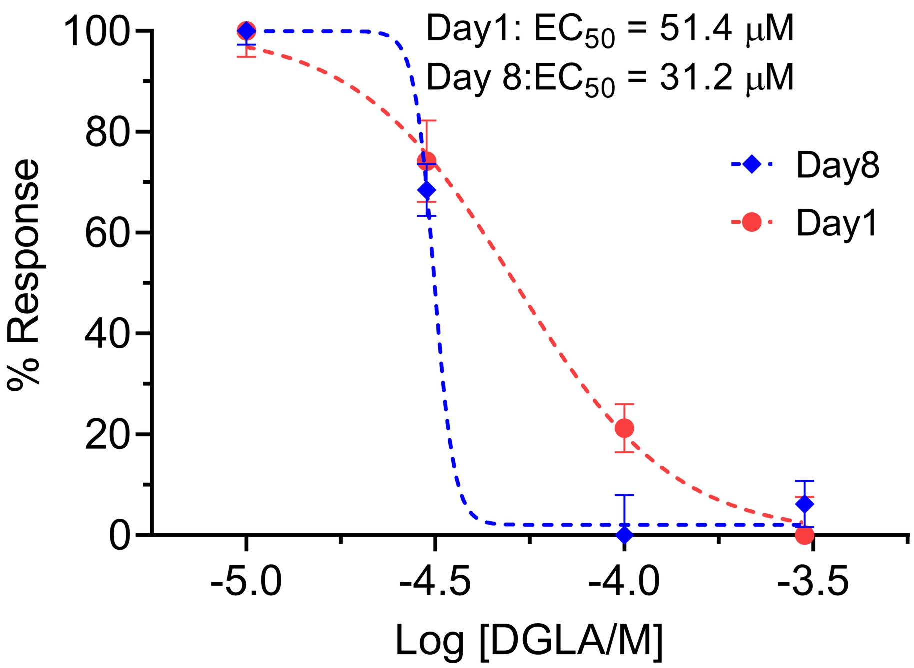

**Figure S1:** Dose response curve: the effect of different DGLA concentrations on degeneration of ADE neurons at Day 1 and Day 8 of adulthood. The slope for dose response curve on day 8 adulthood is significantly different compared to day1 adulthood, suggesting there may be different mechanism for neurodegeneration at these 2 timepoints.

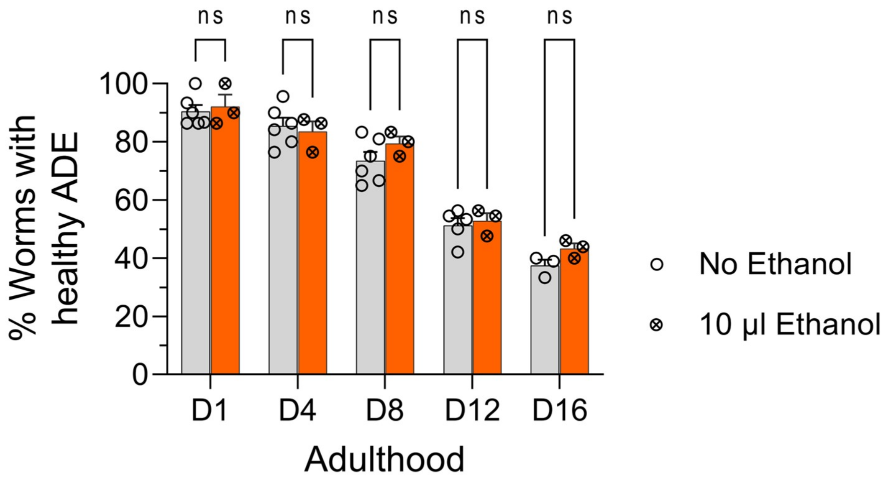

**Figure S2: Ethanol does not alter ADE neuron phenotypes.** Percentage (%) of worms with healthy ADE dopaminergic neurons in *dat-1::gfp* transgenic worms +/- supplementation with 10 µl absolute ethanol. This test was done to determine whether ethanol in PUFA supplementation (which is 10 µl) affects the overall healthspan of dopaminergic neurons. N=3, and about 20 worms were tested in each replicate.. Two-way analysis of variance (ANOVA), Tukey's multiple comparison test, *P ≤ 0.05, **P ≤ 0.01, ***P ≤ 0.001, ****P < 0.0001, ns: not significant.

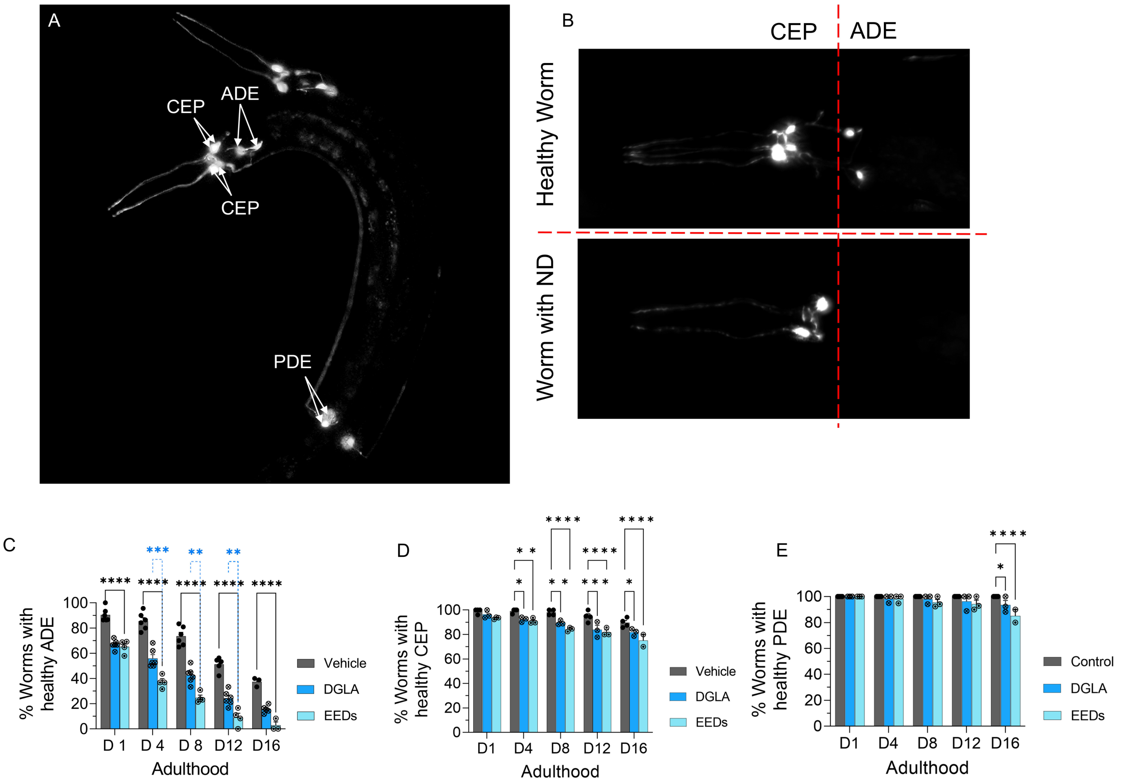

**Figure S3: Different types of dopaminergic neurons in the hermaphrodite have varying sensitivity to the treatment with DGLA or EEDs.** (A). Normal dopaminergic neurons in *C. elegans* labeled with *dat-1::gfp*, (B) Healthy worms (top) and degenerated dopaminergic neurons (bottom). (C) Percentage (%) of worms with healthy ADE dopaminergic neurons in *dat-1::gfp* transgenic +/- supplementation with 100 µM of DGLA or EEDs. (D) Percentage (%) of worms with healthy CEP dopaminergic neurons in *dat-1::gfp* transgenic worms +/1 supplementation with 100 µM of DGLA or EEDs. (E) Percentage (%) of worms with healthy PDE dopaminergic neurons in *dat-1::gfp* transgenic +/- supplementation with 100 µM of DGLA or EEDs. For all experiments N=3, and about 20 worms were tested for each replicate. A two-way analysis of variance (ANOVA), Tukey's multiple comparison test, *P ≤ 0.05, **P ≤ 0.01, ***P ≤ 0.001, ****P < 0.0001, ns: not significant.

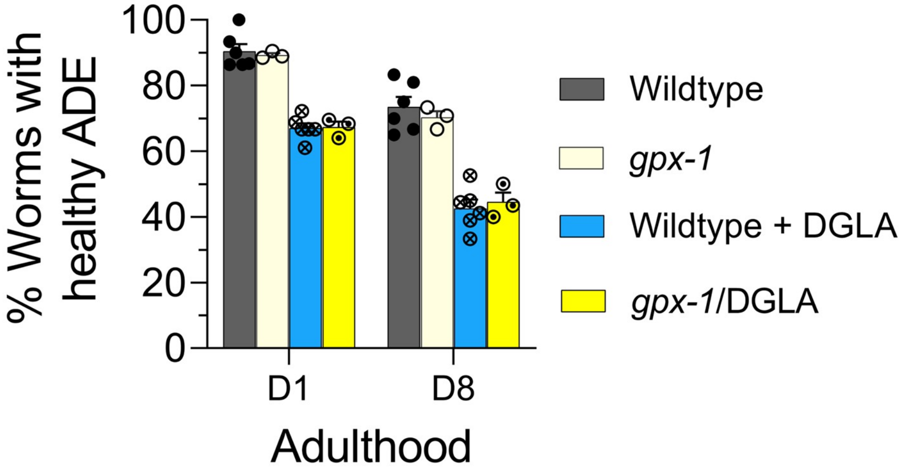

**Figure S4: The genetic knockout of *gpx-1* does not result any observable changes of DGLA-induced neurodegeneration in dopaminergic neurons.** Percentage (%) of worms with healthy ADE neurons for *dat-1::gfp* and *gpx-1;dat-1::gfp* worms treated with 100 µM DGLA were show. This result suggests that DGLA could trigger ferroptosis-mediated neurodegeneration independent of the GPX pathway, which is not without precedent. It has been reported that cytochrome P450 oxidoreductase mediates ferroptosis distinct from the GPX pathway(Zou et al., 2020). Another likely possibility is that there is redundancy, and we have only tested one of the seven isoforms of GPX. More testing on other GPX isoforms is underway.

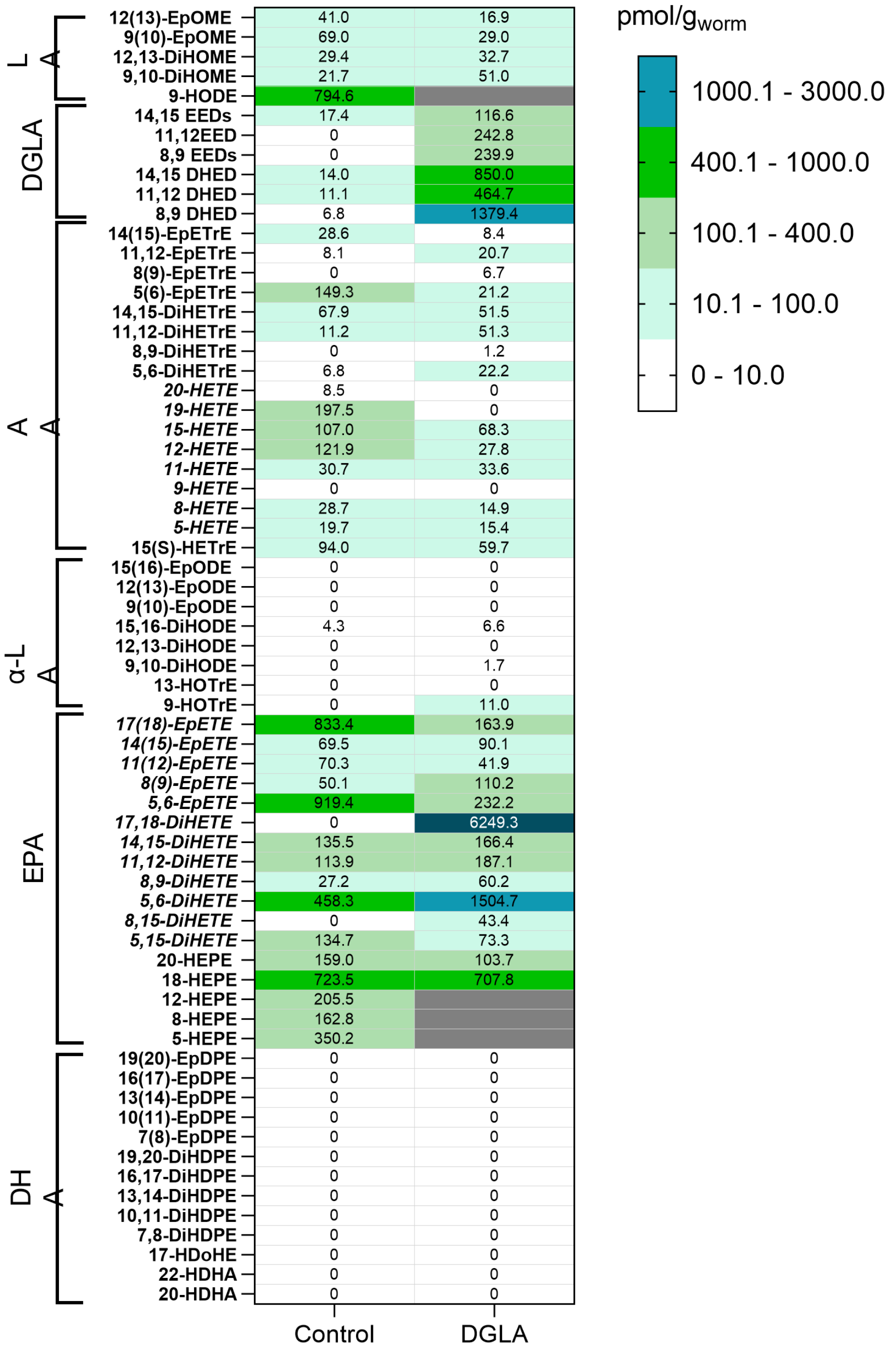

**Figure S5: DGLA Supplementation significantly changes the EEDs and DHEDs levels in worm.** Oxylipin profile representing the pmol/g of Epoxy- and dihydroxy- PUFA levels in worms treated with 100 μM of DGLA compared to control. The worms were supplemented at the L4 stage, and were tested at day 1 of adulthood. Black boxes represent the values are those that were inconsistent in different trials or were out of standard curve range.

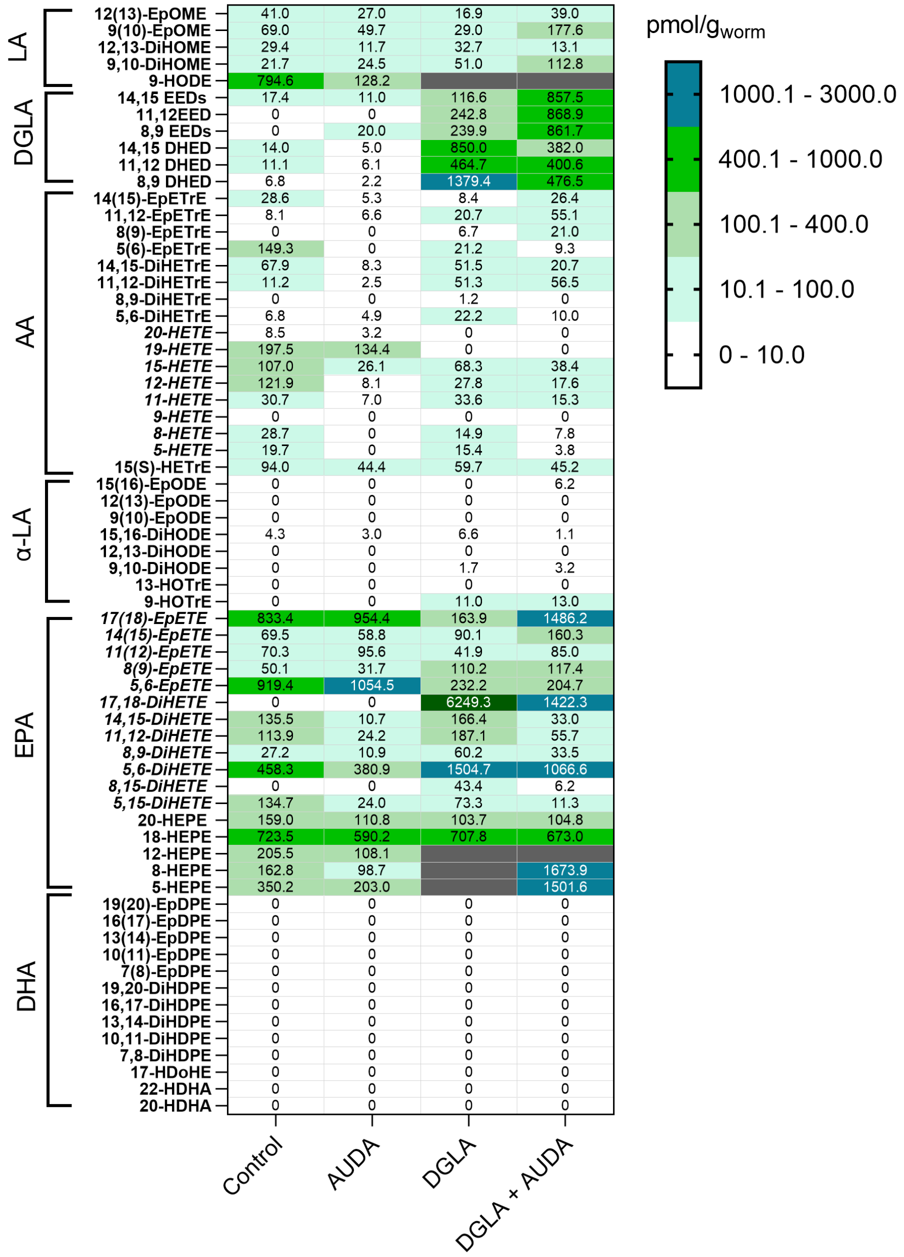

**Figure S6: AUDA changes the EEDs and DHEDs levels in worm supplemented by DGLA, through inhibition of epoxide hydrolase enzyme.** Oxylipin profile representing the pmol/g of Epoxy- and dihydroxy- PUFA level in worms treated with 100 µM of DGLA ±100 µM AUDA compared to control. The worms were supplemented at the L4 stage and were tested at day 1 of adulthood. Black boxes represent the values are those that were inconsistent in different trials or were out of standard curve range.

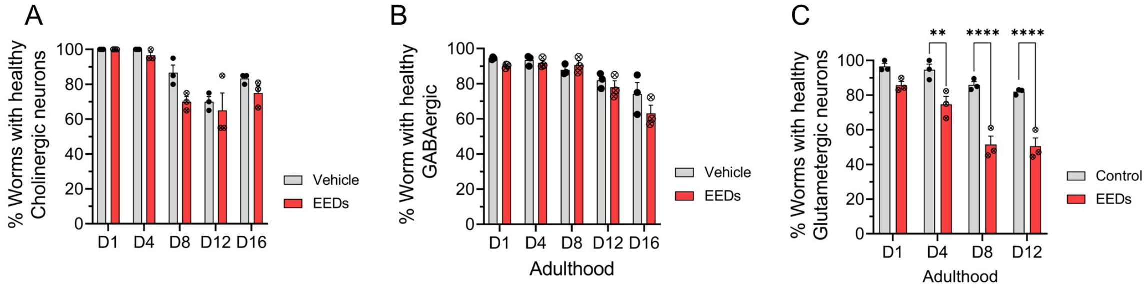

**Figure S7: EEDs does not effect on neuronal healthspan of GABAergic and Cholinergi neurons, and its effect of glutamatergic neurons starts after day 4.** The effect of 100 µM of EEDs on various neurons types compared to vehicle. Percentage (%) of worms healthy (A) cholinergic neurons, (B) GABAergic neurons, and C) glutamatergic neurons. Worms are supplemented with 100 µM of EEDs at the L4 stage. For all experiments N=3, and 20-30 worms were tested in each replicate. A two-way analysis of variance (ANOVA), Tukey's multiple comparison test, $*P \leq 0.05$, $**P \leq 0.01$, $***P \leq 0.001$, $****P < 0.0001$, not significant is not shown.

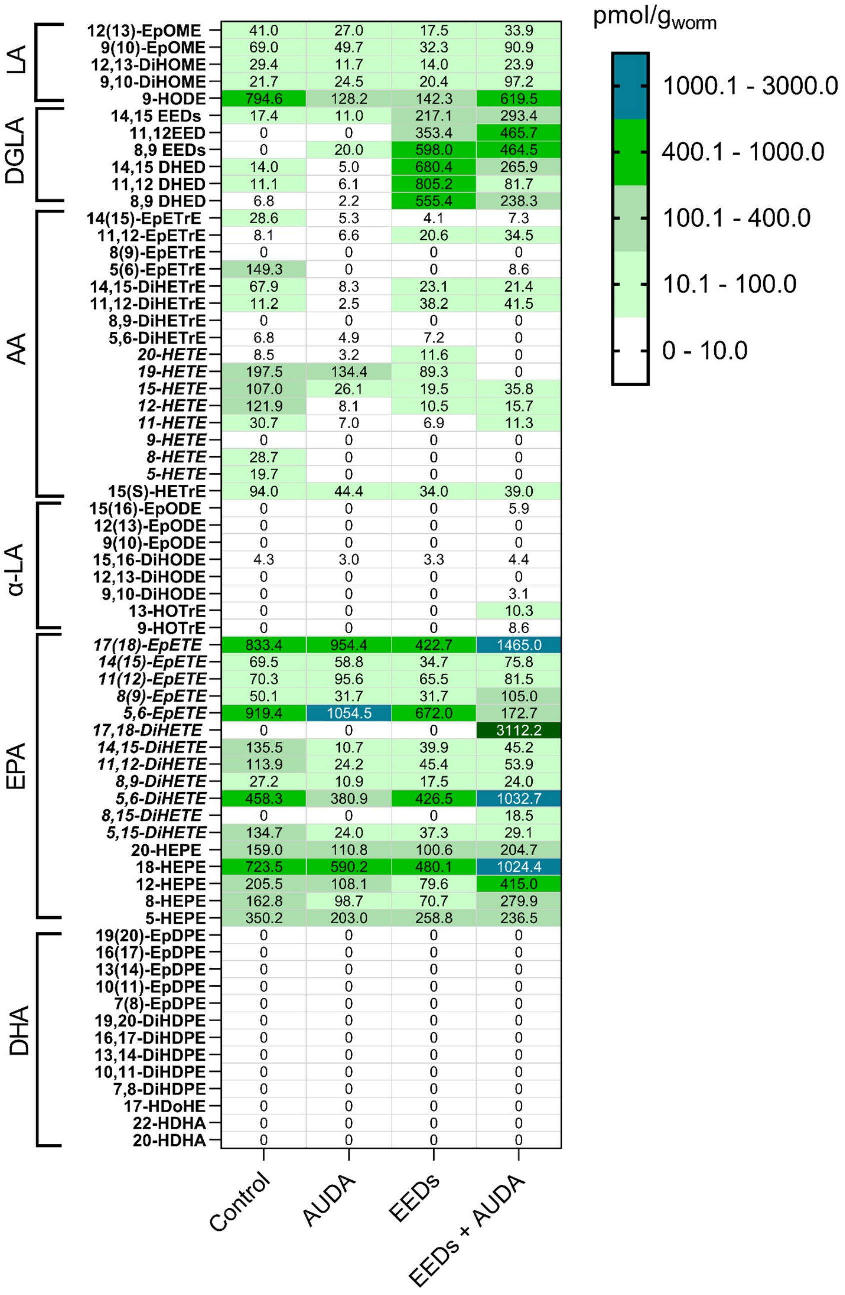

**Figure S8: EEDs Supplementation significantly changes the EEDs and DHEDs levels in worm. AUDA changes the EEDs and DHEDs levels in worm supplemented by DGLA, through inhibition of epoxide hydrolase enzyme.** Oxylipin profile representing the pmol/g of Epoxy- and dihydroxy- PUFA levels in worms treated with 100 μM of EEDs ±100 μM AUDA compared to control. The worms were supplemented at the L4 stage, and were tested at day 1 of adulthood. Black boxes represent the values are those that were inconsistent in different trials or were out of standard curve range.

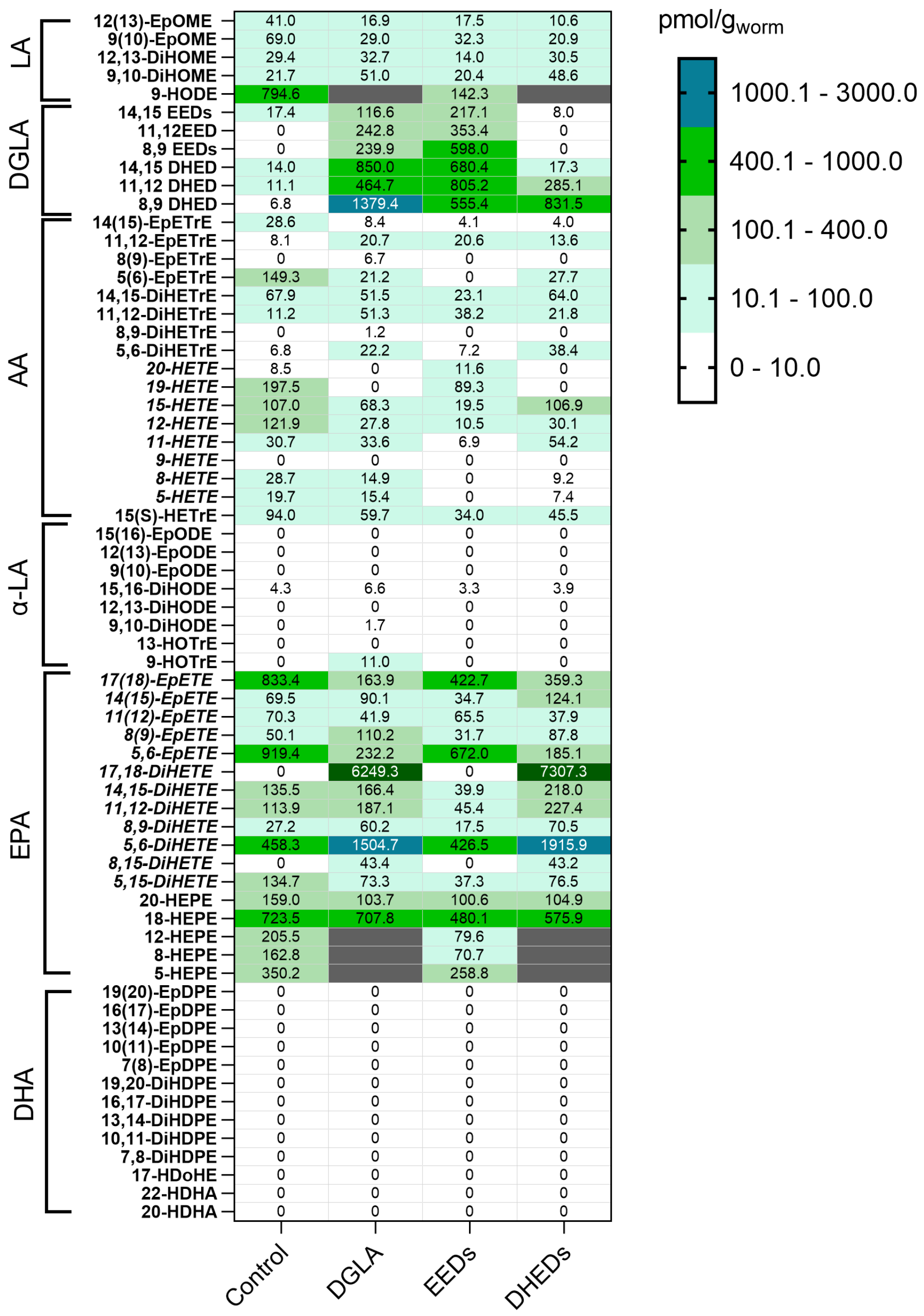

**Figure S9: DGLA, EEDs, and DHEDs Supplementation alters the EEDs and DHEDs levels in worms**. Oxylipin profile representing the pmol/g of Epoxy, hydroxy, and dihydroxy-PUFA regioisomers, CYP/EH metabolites in worms treated with 100 μM of either DGLA or EEDs, or DHEDs compared to control. Worms were supplemented at the L4 stage and were tested at day 1 of adulthood. Black boxes represent the values those that were inconsistent in different trials or were out of standard curve range.

#### 2. Experimental Sections:

**Table 1:** Reagent and resource:

| REAGENT or RESOURCE | SOURCE | IDENTIFIER |
|---|---|---|
| Trolox | Cayman Chemicals | Cat# 10011659; CAS: 53188-07-1 |
| liproxstatin-1 | BioVision | Cat#B2312-5,25; |
| 2,2'-Bipyridine | Oakwood Chemical | Lot# 003434M03F; CAS: 366-18-7 |
| Cholesterol | Alfa Aesar | Cat#A11470; CAS: 57-88-5 |
| Agar | Fisher Bioreagents | Cat#BP9744-500; CAS: 9002-18-0 |
| Bacto Agar | Becton, Dickinson, and Company | Cat# DIFCO 214010 |
| Tryptone | Fisher Bioreagents | Cat#BP1421-500; CAS: 91079-40-2 |
| Bacto Tryptone | Life Technologies Corporation | Cat# DIFCO 211705 |
| Yeast Extract | Becton, Dickinson, and Company | Cat# DIFCO 212750 |
| Sodium Chloride | VWR | Cat#BDH9286 |
| Magnesium Sulfate heptahydrate | Fisher Chemical | Cat#M63-500; CAS: 10034-99-8 |
| Potassium Phosphate, monobasic, crystal | Fisher Bioreagents | Cat#BP362-500; CAS: 7778-77-0 |
| Potassium Phosphate, dibasic, powder | Fisher Chemical | Cat#P288-500; CAS: 7758-11-4 |
| Calcium Chloride (anhydrous) | Sigma-Aldrich | Cat#C1016-500; CAS: 10043-52-4 |
| Sodium Azide | Fisher Scientific | Cat#BP9221-500; CAS: 26628-22-8 |
| Ethanol | Fisher Chemical | Cat#A409-4; CAS: 64-17-4 |
| Arachidonic acid | NU-CHEK | Lot# U-71A-M31-B |
| DGLA | TCI Chemicals | Cat#E0640; CAS:1783-84-2 |
| GLA | NU-CHEK | Lot#U-69A-D16-E |
| LA | NU-CHEK | Lot#U-62A-O21-E |
| EPA | NU-CHEK | Lot#U-99A-MA10-B |
| Hexane | Fisher Chemical | Lot#176581; CAS: 110-54-3 |
| Acetic Acid | Fisher Scientific | Lot#193296; CAS: 64-19-7 |
| Acetonitrile | Fisher Chemical | Lot#195771; CAS: 75-05-8 |
| Chloroform | Acros Organic | Lot# B0541409A; CAS: 67-66-3 |
| Methanol | Fisher Chemical | Lot#195771; CAS: 67-56-1 |
| Acetone | Fisher Chemical | CAS: 67-64-1 |

**Table 2.** Organisms/Strains

| Strain | Source | Strain name |
|---|---|---|
| N2 Bristol | Caenorhabditis Genetics Center | N2 |
| CB767 *bli-3(e767)* I | Caenorhabditis Genetics Center | CB767 |
| GA912 *ftn-1(ok3625)* V | David Gems (Jennifer watts) | GA912 |
| MT1522 *ced-3(n717)* IV | Caenorhabditis Genetics Center | MT1522 |
| FX2100 *gpx-1(tm2100)* | National Bioresource Project | FX2100 |
| BZ555 *[egIs1* [P*dat-1:gfp*)] | Caenorhabditis Genetics Center | BZ555 |
| EM641 *(Pcat-2::gfp)* | A gift from Scott W. Emmons | |
| JKA76 [*ced-3*(n717)*; Pdat::gfp]* | Generated in the lab of Jamie Alan | |
| JKA77 *[bli-3(e767);Pdat-1::gfp]* | Generated in the lab of Jamie Alan | |
| JKA78 *[ftn-1*(ok3625)*;Pdat-1::gfp]* | Generated in the lab of Jamie Alan | |
| JKA79 *[gpx-1(tm2100); Pdat::gfp]* | Generated in the lab of Jamie Alan | |

**Table 3.** Software and Algorithms

| | | |
|---|---|---|
| Microsoft Excel | Microsoft Corporation | N/A |
| ImageJ | Rasband, W.S. | https://imagej.nih.gov/ij/ |
| BioRender | BioRender | https://biorender.com/ |
| GraphPad Prism 6 | GraphPad Software, Inc. | https://www.graphpad.com/ |

#### 2.1. *C. elegans* Strains and Maintenance

All nematode stocks were maintained on nematode growth media (NGM) plates seeded with bacteria (*E. coli* OP50) and maintained at 20°C unless otherwise noted.. N2 Bristol (wild-type), MT1522 *ced-3(n717)*, and CB767 *bli-3(e767)*. The BY250 (P*dat-1::gfp*) was a gift form

Dr. Randy Blakely (Florida Atlantic University). The EM641 (*cat-2::gfp*) strain was a gift from Dr. Scott W. Emmons (Albert Einstein College of Medicine, New York, United States). The GA912 *ftn-1(ok3625)* strain was a gift from Dr. David Gems (University College London, London, UK). Table S2 shows all the strains used in this study.

The JKA 76 [*ced-3*(n717)*; Pdat::gfp], JKA 77 [bli-3(e767);Pdat-1::gfp], and JKA 78 [ftn-1(*ok3625)*;Pdat-1::gfp], and JKA 79 [gpx-1(tm2100); Pdat::gfp]* . strains were constructed using standard methods [1].

#### 2.2. Age synchronized worms:

The age-synchronized population was prepared by transferring specific numbers (depending on the experiments and required number of progeny) of healthy and well-fed Day 1 adult worms to a fresh nematode growth media (NGM) with OP50, as described in previously published protocol [2]. The adult worms were allowed to lay eggs for 6-10 hours. The laid eggs were isolated and allowed to hatch. About 36-48 hours later, plates were washed off with s-basal solution and transferred to a 40 µm cell strainer placed on top of a 50 mL centrifuge tube. The large sized L4 larvae stick to the filter, whereas eggs, larva, bacteria carryover or possible contamination were passed through the filter. L4 larvae were then washed with 75-100 µl of s-basal, transferred to a 1.7 ml centrifuge tube using a glass pipet, and spun at 325 x g on a table-top centrifuge for 30 s. The s-basal solution was removed by aspiration leaving behind a pellet of L4. Finally, L4 worms were resuspend in s-basal solution and transferred to the supplemented or control plates seeded with OP50.

During lifespan, every day the age synchronized population was filtered through a 40 µm cell strainer placed on top of a 50 mL centrifuge tube. The progeny was collected in the filtrate and removed. The age-synchronized adult worms were removed from the surface of the cell strainer and placed on a freshly seeded NGM/supplemented plate. The filtration process was repeated every day during early adulthood of the age synchronized population to avoid any contamination from the progeny, and to provide fresh supplementation for worm during their lifespan.

#### 2.3. Fatty acid supplementation:

In order to supplement worms with fatty acids and/or their downstream metabolites, 10 ul of each compound at desired concentration was spread on the NGM plate, and then immediately seeded with 250-400 µl E. *coli* OP50 (2.8 $\times 10^8$ cell/ml). The seeded plates were sealed with parafilm? and kept for 2 days at room temperature (20-23°C) and then transferred to the refrigerator to be used later. In all experiments in this study, the NGM solution plates were made using standard methods[3].

#### 2.4. Epoxide Hydrolase inhibitor supplementation:

For 12-(3-((3*s*,5*s*,7*s*)-adamantan-1-yl) ureido)dodecanoic acid (AUDA ) supplementation, a 20 mM AUDA stock solution was prepared in ethanol, and then was added to NGM agar solution at 55-65 °C to reach the final concentration of 100 µM before plating. Plates kept at room temperature for 1 day and then seeded by 250-400 ul *E. coli* OP50 (2.8 $\times 10^8$ cell/ml).

#### 2.5. Supplementations for ferroptosis studies:

To study the possible role of DGLA and EEDs supplementation in ferroptosis, 10 µl of 100 µM of DGLA or EEDs was spread on NGM plates, followed by spreading 10 µl of 100 µM of liproxstatin-1 (Lip-1) solution in ethanol. Immediately after that, 250-400 µl E. *coli* OP50 was plated and allowed to dry for two days. The plates were then either used immediately or kept in the fridge (4°C) for later use. The same procedure was used for the 2,2-bipyridine (BP) (100 µM) and Trolox (500 µM), For the control experiments, 10 µl of ethanol solution was used.

#### 2.6. Fluorescence microscopy imaging for tracking dopaminergic neurons.

In order to track neurodegeneration, age-synchronized worms with *dat-1::gfp* transcriptional fusions were used. The age-synchronized worms were analyzed based on a previously published protocol with some modifications[4]. First agarose gel pad were prepared as previously described. (For quantitative analyses of changes in DAergic neuron cell morphology, 20-25 worms were mounted on the layer of the agar pad and paralyzed with 5 mM $NaN_3$ for 5 minutes (Fig.15). Finally, a coverslip was placed onto an agar pad containing worms. A fluorescent microscope (Eclipse Ti2-E–Nikon) was used to image the worms and NIS-Elements software was used to analyze the data. All 8 DAergic neurons were analyzed in each worm. The ADE neurons were the ones with significant neurodegeneration with the treatment of DGLA and its downstream metabolites. Therefore, in all microscopic tests in this study, neurodegeneration refers to the absence of fluorescent signal in the ADE neurons. Worms with healthy ADE are those with both ADE cell bodies or processes that could be seen under fluorescent microscope. The same procedure was followed for dopaminergic neuron analyses using the *cat-2*::gfp (EM641).

#### 2.7. Oxylipin Analysis:

##### 2.7.1. Step 1: Collecting and freezing worm samples for oxylipin analysis

Oxylipins are a class of bioactive oxidized lipid metabolites derived from PUFAs via cyclooxygenase (COX), lipoxygenase (LOX), and cytochrome P450 (CYP) enzymatic pathways. To investigate the oxylipin profile in *C. elegans,* we collected about 5 mg of worms per trial to ensure that the whole worm lysates contain a sufficient concentration of oxylipins for detection. A sufficiently sized population of worms was generated using a minimum of 7 P100 plates with a diameter of 100 mm?, per trial. To generate 5 mg of whole worm lysates, we prepared approximately 2000-3000 worms (300-400 worms per plate). The age-synchronized population of worms was generated and maintained using the filtration method illustrated and described above. When a population of worms was ready for isolation and collection, the entire population of seven plates per trial was transferred and filtered using s-basal solution and a cell strainer with a pore size of 40 µm. The worms that collected on the surface of the cell strainer were transferred using a Pasteur pipet to an Eppendorf vial to avoid *C. elegans* from sticking on the wall of the pipet. The worms were rinsed with s-basal medium and centrifuged. The supernatant was collected and discarded. The worms were then washed four more times with s-basal medium to ensure that all bacteria and PUFA supplements were removed. After the bacteria and supplements were removed, the Eppendorf vials containing each worm sample were transferred to a benchtop centrifuge. The vials were centrifuged for 10 minutes at 10,000 rpm at 4°C. The supernatant was removed using 100 µL and 10 µL pipets. A 20 µL pipet with a long tip was pushed to the bottom of the vial to remove the liquid between the worms. Lastly, the standard filter paper was cut and inserted into the Eppendorf vials to remove any remaining liquid within the worm sample. After all liquid was removed, the worm samples flash frozen using liquid nitrogen and stored in the -80°C freezer. Collection of worms for oxylipin and lipidomic analysis is illustrated in the Figure S10.

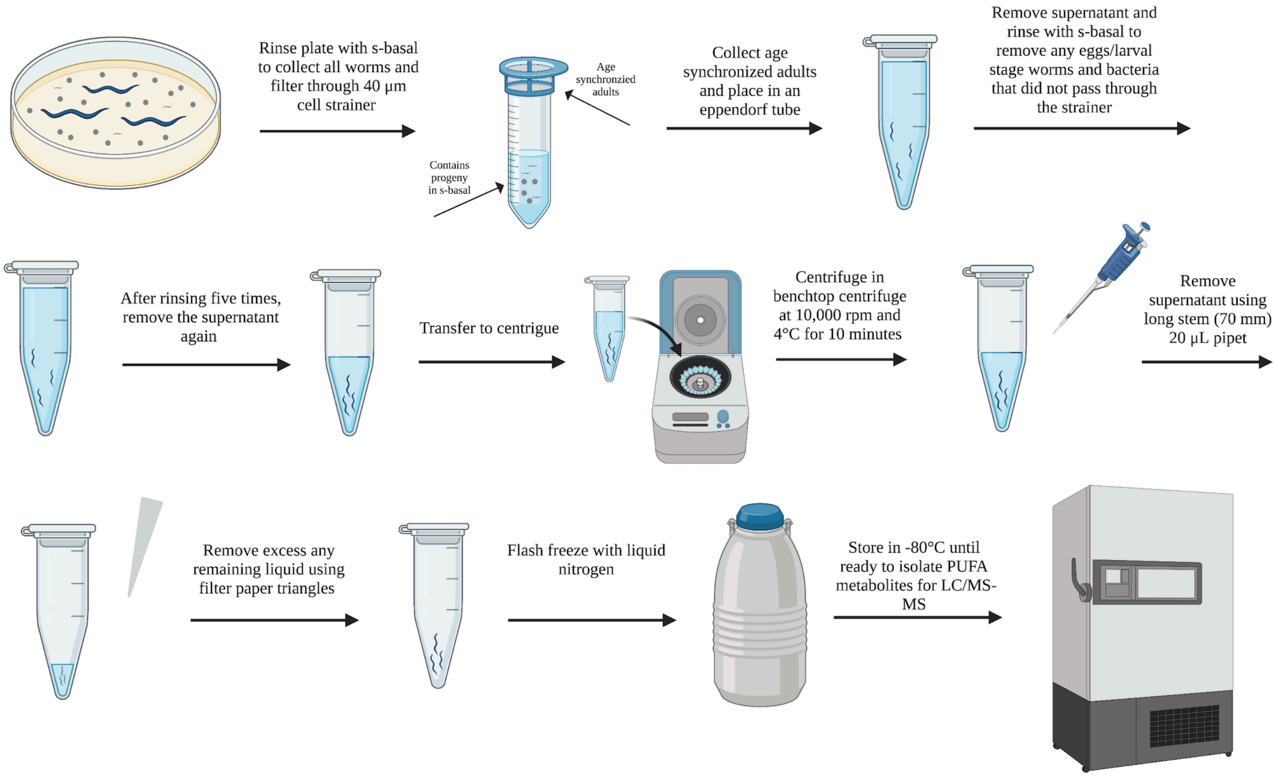

**Figure S10.** Worm sample preparation for oxylipin and analysis

2.7.2. Step 2: Worm homogenization for oxylipin analysis

Eppendorf vials containing worms were removed from -80°C storage. The weight of one 2 mL cryogenic homogenizer vial per trial was recorded. The worm samples were flash frozen using liquid nitrogen, and the frozen worm samples was broken loose using a 0.7 mm needle. The worm samples were transferred to the homogenizer vial and the weight was recorded. The weight of each vial with the worms was measured to determine the weight of worms used for each trial. Three homogenization beads were added to each homogenizer vial. Additionally, 100 µL phosphate-buffered saline (PBS), 10 µL of internal standard, consisting of 10 µL deuterated oxylipins, and 10µL of antioxidants, consisting of ethylenediamine tetraacetic acid (EDTA) (1 mg/ml in water), butylated hydroxytoluene (BHT) (0.2 mg/ml in methanol), and triphenylphosphine (TPP) (0.2 mg/ml in Ethanol). The details of the deuterated oxylipin standards are shown in Table S4**.** Each homogenizer vial containing the worm samples was flash frozen using liquid nitrogen and then homogenized for five 30-second cycles at 5 M/s using an Omni bead ruptor 24 homogenizer. After homogenization, an additional 900 μL of PBS was added to the homogenized sample. The sample was centrifuged using a benchtop centrifuge at 10,000 rpm for 5 minutes. The supernatant was collected and transferred to a new Eppendorf for solid phase extraction. The process of homogenization is illustrated in the Figure S11.

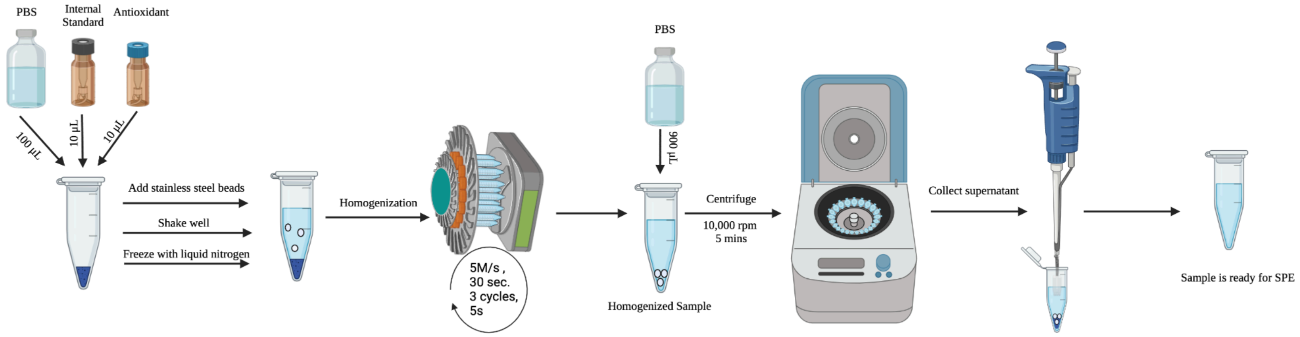

**Figure S11.** Worm homogenization for oxylipin analysis

**Table 4**: Deuterated standards used for oxylipin analysis.

| Oxylipin standard name | Oxylipin standard abbreviation |
|---|---|
| 6-keto prostaglandin $F_{1\alpha}$-d4 | 6-keto-$PGF_{1\alpha}$-d4 |
| 5(S)-hydroxyeicosatetrenoic-d8 acid | 5(S)-HETE-d8 |
| 8,9-epoxyeicosatrienoic-d11 acid | 8,9-EET-d11 |
| Arachidonic-d8 acid | AA-d8 |
| 15(S)-hydroxyeicosatetraenoic-d8 acid | 15(S)-HETE-d8 |
| Prostaglandin $B_2$-d4 | PGB2-d4 |
| 8,9-dihydroxyeicosatrienoic-d11 acid | 8,9-DiHETrE-d11 |
| 9(S)-hydroxyoctadecadienoic-d4 acid | 9(S)-HODE-d4 |
| Leukotriene $B_4$-d4 | LTB4-d4 |
| Prostaglandin $E_2$-d9 | PGE2-d9 |

##### 2.7.3. Step 3: Solid phase extraction to isolate the oxylipins from the whole worm lysate

To isolate the oxylipins from the whole worm lysates solid phase extraction (SPE) (Waters Oasis-HLB cartridges, (Part No. WAT094226, Lot No. 176A30323A) was used. We used a polar stationary phase to trap the extremely polar biological material such as sugars. The oxylipins that we are isolating are significantly less polar in comparison. The SPE column was prepared by sequential washing with 2 mL ethyl acetate, 2 mL methanol twice, and 2 mL of 95:5 (v/v) mixture of water and methanol containing 0.1% acetic acid. The column was kept moist during preparation. The process of SPE column preparation is illustrated in the Figure S12.

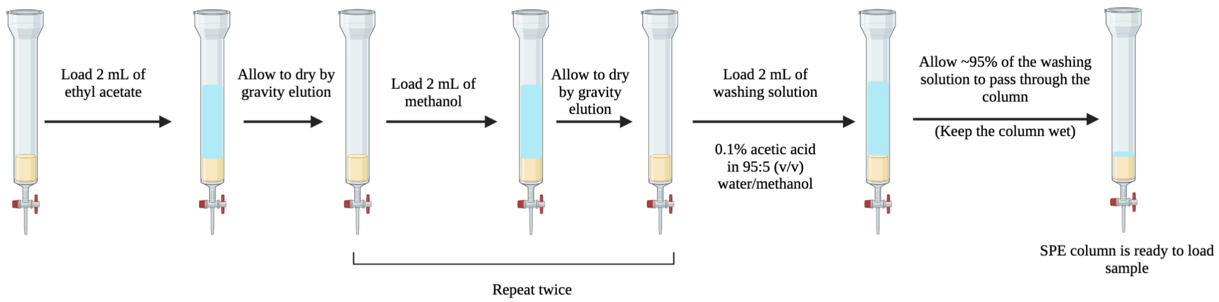

**Figure S12.** Solid phase extraction column preparation

After the SPE column was prepared, the Eppendorf vials containing the homogenized samples were loaded onto the SPE column. After the column was loaded with the sample by gravity, 1.5 mL of the washing solution, 95:5 (v/v) mixture of water and ethanol with 0.1% acetic acid, was added to the column. The column was dried by gravity. Next, the column was thoroughly dried with a vacuum pump for 20 minutes. After thorough drying, the column was ready for elution. The process of loading the sample to the SPE column is illustrated in the Figure S13**.**

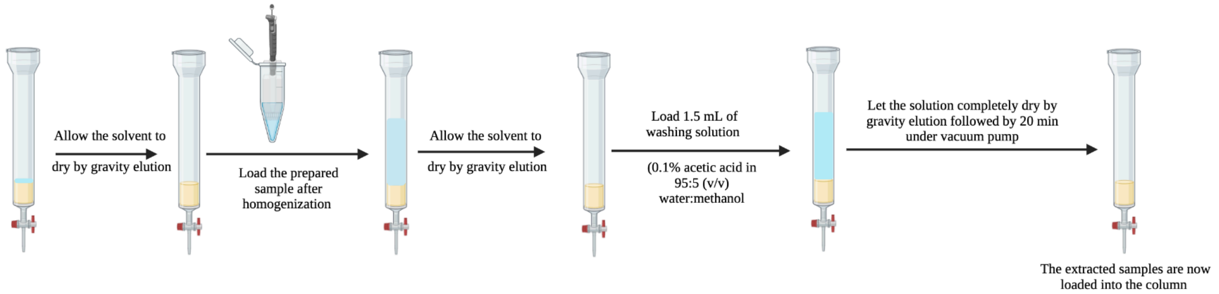

**Figure S13.** Loading the sample to the SPE column

After the column was loaded with the sample and completely dried, 0.5 mL of methanol was added to begin the elution step. Eluted compounds were collected to an Eppendorf vial containing 6 µL of 30% glycerol in methanol, which serves as a trap solution. The column was allowed to gravity elute until the column appeared dry. A 5 mL syringe was filled with air and placed on the top of the SPE column to gently push the remaining solvent out of the column with air. Once the column was completely dry, 1 mL of ethyl acetate was added to the column. The solvent was allowed to gravity elute until the column appeared dry to the eye. The remaining solvent was again removed using a 5 mL syringe and gently pushing air through the column. The process of eluting is illustrated in the Figure S14.

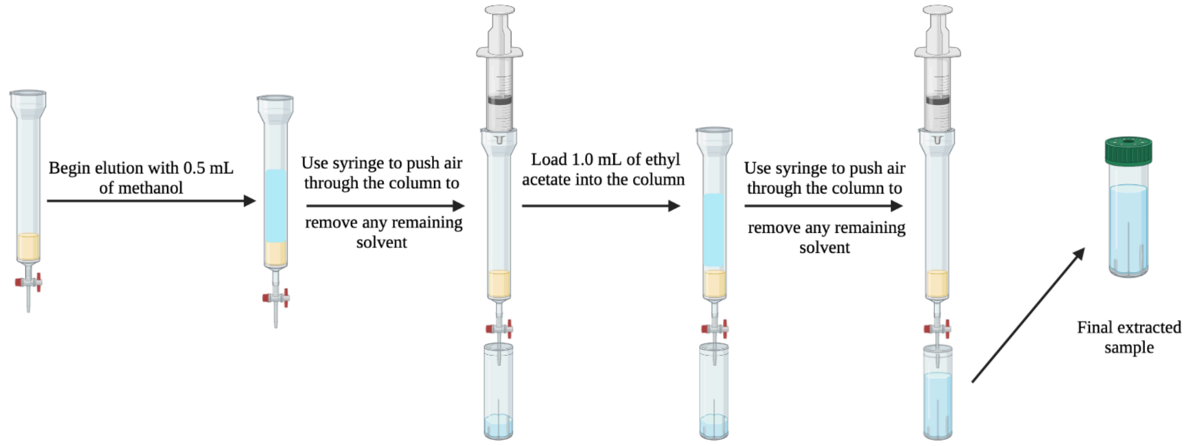

**Figure S14.** Elution of oxylipins from SPE column.

Upon completion of SPE, the final extracted sample was dried using a speed-vac until the trap solution was all that remained. The residues were reconstituted with 100 µL of 75% methanol/water containing 10nM of internal standard, 12-[(cyclohexylcarbamoyl)amino]dodecanoic acid (CUDA). The samples were then mixed on a vortex for five minutes and filtered with a 0.45 µm filter. Lastly, the samples were transferred to auto-sampler vials with salinized inserts, purged with argon gas, and stored at -80°C until injection.

2.7.4. <u>Step 4: Oxylipin analysis using LC/MS-MS</u>

The LC conditions were optimized to separate all eicosanoids of interest with the desired peak shape and signal intensity using an XBridge BEH C18 2.1x150mm HPLC column. The mobile phase A comprised of 0.1% acetic acid in water. Mobile phase B consisted of acetonitrile: methanol (84:16) with 0.1% glacial acetic acid. Gradient elution was performed at a flow rate of 250 µL /min. Chromatography was optimized to separate all analytes in 20 min. The autosampler, Waters ACQUITY FTN, was kept at 10°C. The column was connected to a TQXS tandem mass spectrometer (Waters) equipped with Waters Acquity SDS pump and Waters Acquity CM detector. Electrospray was operated as ionization source for negative multiple reaction monitoring (MRM) mode. To generate the best selectivity and sensitivity, each analyte standards were infused into the mass spectrometer and multiple reaction monitoring was used to analyze the desired compound.

#### 2.8. Synthesis of Diepoxyeicosaenoic acid (DiEEMe) and DLGA THF-diols

##### 2.9.1. General synthesis Methods

Reactions using air-sensitive reagents were conducted under an inert atmosphere of argon. All purchased chemicals were used as received (without further purification). Dihomo-gamma linoleic acid (DGLA) was purchased from Nu-Chek-Prep**.** SiliCycle irregular silica gel p60 (60, 230-400 mesh) was used for column chromatography and AnalTech 250 micron silica gel plates were used for analytical thin layer chromatography and visualized using either vanillin or $KMnO_4$ stain. $^{1}H$ and $^{13}C$ NMR spectra were collected at 25°C on Carian Inova 500 MHz instrument and reported in parts per million (ppm) relative to the solvent resonances (δ), with coupling constants (J) in Hertz (Hz). High-resolution mass spectroscopy (HRMS) data was collected on a Waters Xevo G2-XS UPLC/MS/MS Quadrupole/Time-of-Flight instrument. High-performance liquid chromatography (HPLC) was performed on a Rainin HPXL instrument with a Dynamax Absorbance Detector Model UV-D detection system with a Zorbax Sil 9.4 mm x 25 cm column (5 micron particle size, 100 Angstrom pore size). Gas-chromotography/mass spectroscopy (GC/MS) was performed on an Agilent Technologies 7890A GC system with an Agilent Technologies 5975C inert XL EI/CI MSD Triple-Axis Detector, and Agilent HP-5ms Fused Silica Capillary Column with 0.25 micron film thickness, 30 m long, and 0.25 mm inner diameter.

##### 2.9.2. Synthesis of Diepoxyeicosaenoic acid (DiEEMe) (1a – 1c)**:**

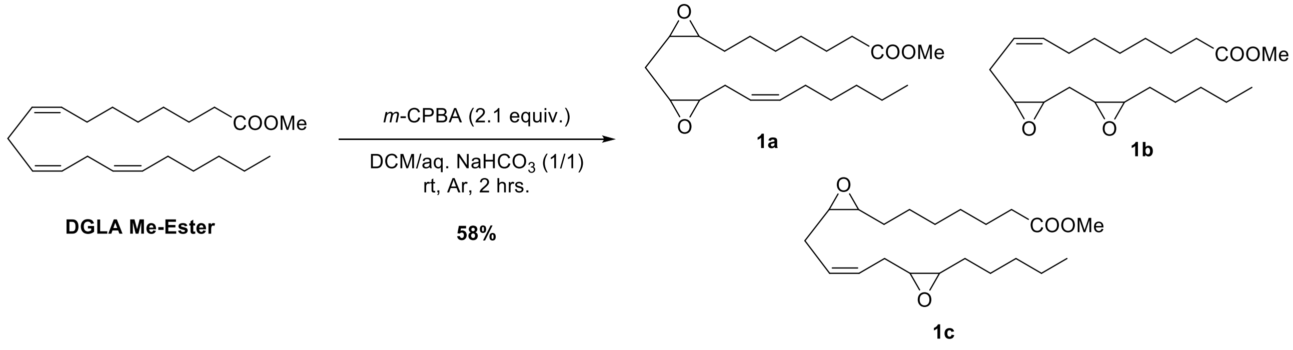

Methyl dihomo-gamma linoleate (1.569 mmol, 0.500 grams, 1 equiv.) was dissolved in 40 mL of dichloromethane (DCM). Meta-chloroperbenzoic acid (*m*-CPBA) (3.296 mmol, 0.813 grams, 2.1 equiv.) was added followed by saturated aqueous $NaHCO_3$ (40 mL) and the reaction was stirred vigorously under argon atmosphere at rt for 2 hours. Then, the organic layer was separated and collected. The aqueous phase was extracted with DCM (20 mL) for three times. The combined organic layers were washed with saturated aqueous $NaHCO_3$ (3 x 15 mL), brine (1 x 15 mL), dried over $Na_2SO_4$, and concentrated under reduced pressure to afford a clear oil. The crude product was purified with column chromatography (5–10% EtOAc/Hexanes) to afford **1a – 1c** as a clear oil (mixture of regioisomers) (0.318 grams, 58% yield). HRMS (ES+): calculated $C_{21}H_{36}O_4Na+$ [M+Na]+ 375.2506 observed 375.2519. For $^1H$ and $^{13}C$ NMR of mixture and separated fractions, see spectra below.† See characterization in Appendix.

2.9.3. Synthesis of DLGA THF-diols (2 – 3):

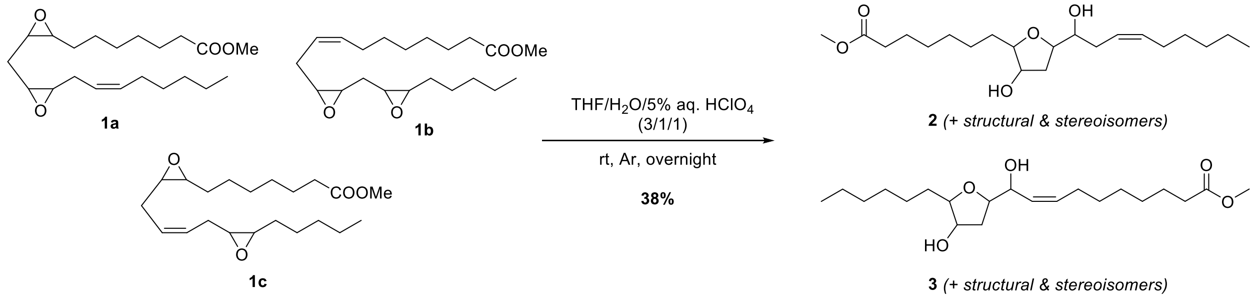

DGLA diepoxides (**1a – 1c**) (0.902 mmol, 0.318 grams, 1 equiv.) was dissolved in 7.5 mL of aq. 5% $HClO_4$, THF, and $H_2O$ (1:3:1) and stirred at rt under argon atmosphere overnight. Then, the reaction was cooled to 0°C and quenched with 15 mL of saturated aqueous $NaHCO_3$. The reaction was diluted with 25 mL of EtOAc, the layers separated and the organic layer was collected. The aqueous phase was extracted with 3 x 10 mL of EtOAc. The combined organic layers were washed with saturated aqueous $NaHCO_3$ (3 x 15 mL), brine (1 x 15 mL), dried over $Na_2SO_4$, and concentrated under reduced pressure to afford the crude product as a clear oil. Purification via column chromatography (50% EtOAc/Hexanes) gave **2 – 3** as a clear oil (mixture of structural and stereoisomers) (0.109 grams, 33% yield). HRMS (ES+): calculated $C_{21}H_{38}O_5Na+$ [M+Na]+

393.2611 observed 393.2625. For $^{1}H$ and $^{13}C$ NMR of mixture, see spectra below.† See characterization in Appendix.

##### 2.9.4. Determination of Isomeric Ratios of DGLA THF-Diols:

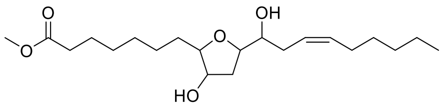

**2**

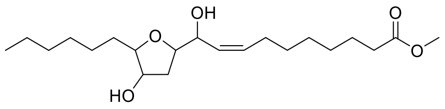

**3**

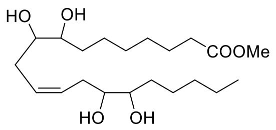

**4**

Isomers of a mixture of DGLA diepoxides (**1a–1c**) were separated via HPLC (1% isopropanol/hexanes, 2 mL/min), giving 5 fractions((retention times, in minutes, were 18.6, 22.1, 23.8, 25.5, and 27.6 for fractions 1 – 5, respectively). Each fraction was subjected to $^{1}H$ and gCOSY NMR analysis (see pages 48 – 57 for $^{1}H$ NMR and gCOSY spectra), leading to identification of isomer **(4)** and showing that the HPLC separation yielded two diastereomeric pairs of isomers **2** & **3**. Following acid hydrolysis of each isolated fraction, isomer **4** was confirmed with low resolution mass spectroscopy (LRMS).

To prepare samples for GC/MS analysis, each of the other 4 fractions were individually dissolved in 1 mL of THF and incubated with 15 equivalents of N,O-Bis(trimethylsilyl)trifluoroacetamide (BSTFA) and pyridine at 60°C for 30 minutes. The samples were dried under a stream of nitrogen, resuspended in DCM, and analyzed via GC/MS. Unique fragmentation patterns then successfully identified isomers **2** and **3** (see HPLC trace and GC/MS data, pages 62 – 70). Once the peaks (F1 – F5, see HPLC trace) were identified, relative ratios were determined by taking the total area of each diastereomeric isomer pair divided by the total area of all peaks (F1 – F5) obtained on HPLC trace.

2.9.5. Synthesis of DGLA monoepoxide esters (5a – 5c):

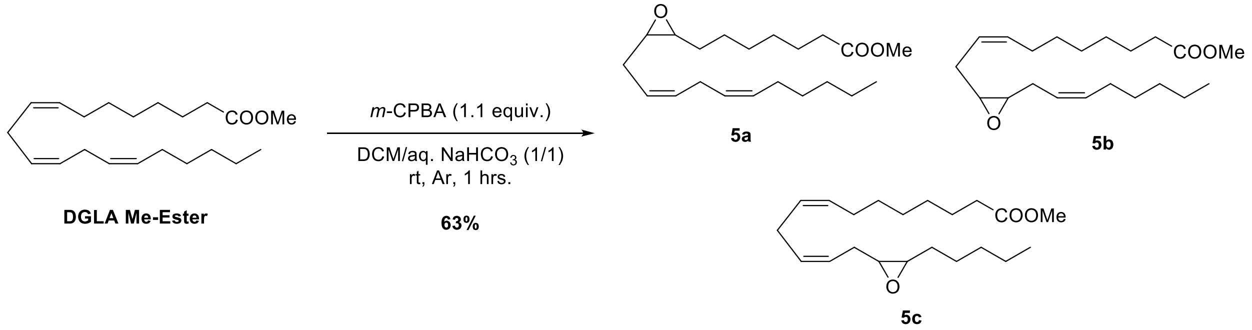

Methyl dihomo-gamma linoleate (3.112 mmol, 1.000 grams, 1 equiv.) was dissolved in DCM (80 mL). Meta-chloroperbenzoic acid (*m*-CPBA) (3.423 mmol, 0.767 grams, 1.1 equiv.) was added followed by saturated aqueous $NaHCO_3$ (80 mL) and the reaction stirred vigorously under argon atmosphere at room temperature for 1 hour. Then, the layers were separated, and the aqueous phase extracted with DCM (3 x 20 mL). The combined organic layers were washed with saturated aqueous $NaHCO_3$ (3 x 15 mL), brine (1 x 15 mL), dried over $Na_2SO_4$, and concentrated under reduced pressure to afford a clear oil. The crude product was purified with column chromatography (5–10% EtOAc/Hexanes) to afford **5a – 5c** as a clear oil (mixture of regioisomers,0.513 grams, 49% yield). Product **5a – 5 c** was carried forward without further characterization.

2.9.6. Synthesis of dihydroxyeicosadienoic acid (DHEDs) (6a – 6c):

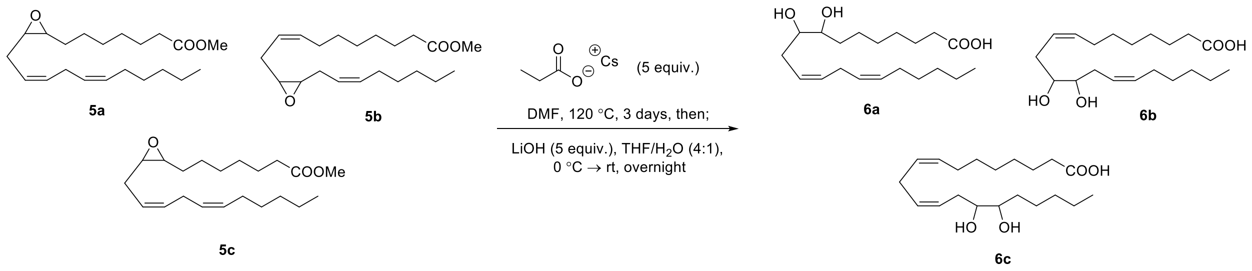

A mixture of **5a–5c** (0.200 g, 0.598 mmol) was diluted in anhydrous DMF (2 mL) and added to dry cesium propionate (0.621 g, 2.98 mmol) in DMF (5 mL) in a sealable tube. The tube was flushed with argon, sealed, and heated to 120 ˚C for 68 h. The mixture was cooled, poured into $H_2O$ (20 mL) and extracted with EtOAc (3 x 30 mL). The organic phase was washed with 5% NaCl (2 x 50 mL) and saturated aqueous NaCl (50 mL), dried over $Na_2SO_4$, and concentrated. The residue was purified by flash column chromatography (CombiFlash® Rf+ Lumen) (25 g $SiO_2$ cartridge, 0 – 50% EtOAc/Hexanes) to yield the EED propionate intermediate as a pale-yellow syrup (0.220 g, 90%) after drying in vacuo (See Table 5 for solvent gradient used for EEDs Methyl ester).

**Table 5**. Solvent gradient for EEDs methyl ester separation.

| Duration (minutes) | %B | Solvent A | Solvent B |
|---|---|---|---|
| 0.0 | 0.0 | Hexane | Ethyl Acetate |
| 2.0 | 0.0 | Hexane | Ethyl Acetate |
| 7.0 | 5.0 | Hexane | Ethyl Acetate |
| 10.0 | 10.0 | Hexane | Ethyl Acetate |
| 5.0 | 12.0 | Hexane | Ethyl Acetate |
| 8.0 | 50.0 | Hexane | Ethyl Acetate |
| 3.0 | 0.0 | Hexane | Ethyl Acetate |

The mixture of EED propionates was then diluted in THF/$H_2O$ (5.5/1.4 mL) and cooled to 0˚C under argon. LiOH (1.21 mL of a 2 M solution in $H_2O$, 2.42 mmol) was added, and the mixture was warmed to rt and stirred overnight. The mixture was then quenched by the dropwise addition of formic acid, until the pH of the mixture was approximately 4. $H_2O$ and EtOAc (10 mL each) were then added, and the layers were separated. The aqueous layer was extracted with EtOAc (4 x 10 mL) and the organic phase was washed with saturated aqueous NaCl (40 mL), dried over $Na_2SO_4$, concentrated, and azeotroped with hexanes (3 x 20 mL) to remove residual formic acid. The residue was then purified by flash column chromatography (25 g $SiO_2$ cartridge, 40 – 75% EtOAc/Hexanes) to yield the DGLA diol mixture **6a – 6c** (0.119 grams, 65% yield). Regioisomers were separated via HPLC with [CONDITIONS]. **6a**; $^1$H NMR ($CDCl_3$, 500 MHz): δ 5.57 – 5.28 (m, 4H), δ 3.50 – 3.44 (m, 2H), δ 2.80 (t, *J* = 7.3 Hz, 2H), δ 2.34 – 2.29 (m, 2H), δ 2.04 (q, *J* = 7.2 Hz, 2H), δ 1.66 – 1.60 (m, 2H), δ 1.49 – 1.23 (m, 12H), δ 0.88 (t, *J* = 6.9 Hz, 3H). $^{13}$C ($CDCl_3$, 125 MHz): δ 179.4, 131.8, 130.9, 127.4, 125.1, 74.0, 74.0, 34.1, 33.6, 31.7, 31.6, 29.4, 29.3, 29.1, 27.4, 25.9, 25.5, 24.7, 22.7, 14.2. **6b**; $^1$H NMR ($CDCl_3$, 500 MHz): δ 5.59 – 5.39 (m, 4H), δ 3.54 – 3.51 (m, 2H), δ 2.35 – 2.26 (m, 6H), δ 2.07 – 2.03 (m, 4H), δ 1.66 – 1.60 (m, 2H), δ 1.39 – 1.24 (m, 12H), 0.88 (t, *J* = 6.8 Hz, 3H). $^{13}$C ($CDCl_3$, 125 MHz): δ 179.2, 133.8, 133.4, 125.0, 124.7, 73.5, 73.4, 34.0, 31.9, 31.9, 31.7, 29.4, 29.3, 28.9, 28.8, 27.5, 27.3, 24.7, 22.7, 14.2. **6c**: $^1$H NMR ($CDCl_3$, 500 MHz): δ 5.57 – 5.31 (m, 4H), δ 3.52 – 3.47 (m, 2H), δ 2.82 (t, *J* = 6.8 Hz, 2H), δ 2.36 – 2.31 (m, 4H), δ 2.07 – 2.03 (m, 2H), δ 1.65 – 1.60 (m, 2H), δ 1.53 – 1.46 (m, 2H), δ 1.40 – 1.25 (m, 12H), δ 0.90 (t, *J* = 7.0 Hz, 3H).

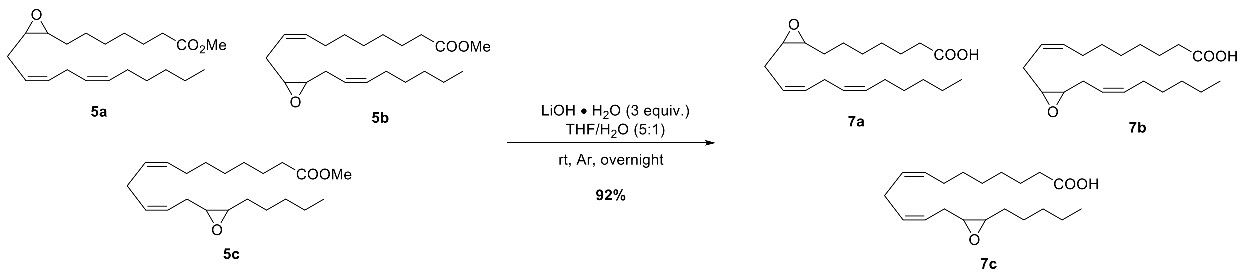

2.9.7. Synthesis of epoxyeicosadienoic acid (EEDs)(7a–7c):

Each separated isomer **5a–5c** was individually subjected to the following conditions: The methyl ester (0.0891 mmol, 0.0300 g, 1.00 equiv.) was added to a 5 mL round bottom flask with a stir bar and diluted with 0.750 mL of THF/$H_2O$ (5:1). To this solution was added LiOH • $H_2O$ (0.267 mmol, 0.00640 g, 3 equiv.) and allowed to stir under argon atmosphere overnight. Then, the pH was adjusted to 4 with formic acid, diluted with water and EtOAc, and added to a separatory funnel. The aqueous layer was extracted with 4 x 5 mL of EtOAc, the combined organic layer washed with brine, dried over $Na_2SO_4$, and concentrated. The residue was azeotroped with 4 x 10 mL of hexanes to remove any remaining formic acid. The crude product was purified via column chromatography (1/1 hexanes/EtOAc, 1% formic acid) to give the products as clear oils. **7a**; $^1$H NMR ($CDCl_3$, 500 MHz): δ 5.53 – 5.29 (m, 4H), δ 2.97 – 2.91 (m, 2H), δ 2.80 (t, *J* = 6.6 Hz, 2H), δ 2.43 – 2.34 (m, 3H), δ 2.24 – 2.19 (m, 1H), 2.05 (q, *J* = 7.1 Hz, 2H), δ 1.68 – 1.62 (m, 2H), δ 1.57 – 1.24 (m, 14H), δ 0.89 (t, *J* = 6.9 Hz, 3H). $^{13}$C ($CDCl_3$, 125 MHz): δ 179.4, 131.0, 130.9, 127.3, 124.2, 57.3, 56.6, 34.0, 31.7, 29.4, 29.3, 29.1, 27.8, 27.4, 26.6, 26.4, 26.0, 24.7, 22.7, 14.2. **7b**; $^1$H NMR ($CDCl_3$, 500 MHz): δ 5.56 – 5.39 (m, 4H), δ 2.97 – 2.93 (m, 2H), δ 2.44 – 2.38 (m, 2H), δ 2.34 (t, *J* = 7.5 Hz, 2H), δ 2.24 – 2.18 (m, 2H), δ 2.07 – 2.03 (m, 4H), δ 1.66 – 1.60 (m, 2H), 1.40 – 1.24 (m, 14H), 0.88 (t, *J* = 6.8 Hz, 3H). $^{13}$C ($CDCl_3$, 125 MHz): δ 179.5, 133.0, 132.7, 124.1, 123.8, 56.7, 56.7, 34.0, 31.6, 29.4, 29.4, 29.1, 29.0, 27.6, 27.5, 26.3, 24.7, 22.7, 14.2. **7c**; $^1$H NMR ($CDCl_3$, 500 MHz): δ 5.53 – 5.31 (m, 4H), δ 2.98 – 2.93 (m, 2H), δ 2.8 (t, *J* = 6.9 Hz, 2H), δ 2.44 – 2.39 (m, 1H), δ 2.35 (t, *J* = 7.6 Hz, 2H), δ 2.24 – 2.19 (m, 1H), δ 2.07 – 2.03 (m, 2H), δ 1.66 – 1.61 (m, 2H), δ 1.57 – 1.25 (m, 14H), δ 0.89 (t, *J* = 7.0 Hz, 3H). $^{13}$C ($CDCl_3$, 125 MHz): δ 179.0, 130.9, 130.6, 127.5, 124.3, 57.5, 56.7, 34.0, 31.9, 29.5, 29.0, 29.0, 27.8, 27.3, 26.4, 26.4, 26.0, 24.8, 22.7, 14.2.

† Compounds were used as mixtures for biological testing. The LC/UV-vis and LC/MS/MS oxylipin analysis was used to confirm the mixture of regioisomer and the purity of the compound

Figure S16 shows the different regioisomers ratio in EEDs and DHEDs supplementation used in this study. Confirmation of desired structure was done by examining ratio of integrations of alkene protons (~5 – 6 ppm) to the methyl ester (~3.6 ppm) on $^{1}H$ NMR. HRMS was also done to ensure desired mass. Separations and characterization were performed to determine relative percentages of each isomer in mixture used for biological testing.

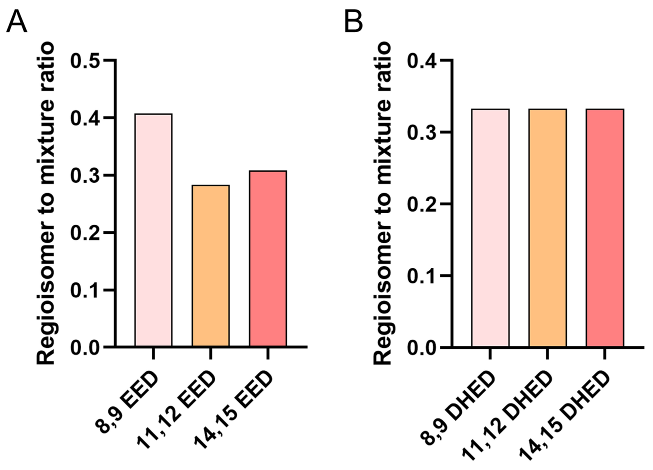

**Figure S16.** The different regioisomers ratio in EEDs and DHEDs supplementation used in this study.

2.9.8. Synthesis of 12-(3-((3*s*,5*s*,7*s*)-adamantan-1-yl)ureido)dodecanoic acid (AUDA):

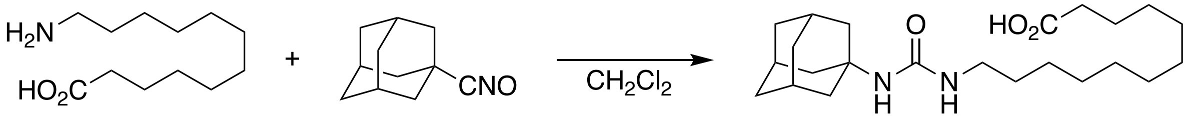

The synthesis followed published procedure [5]. To a suspension of 12-aminododecanoic acid (1g, FW = 215.33, 95% purity, 4.4 mmol) in 1,2-dichloroethane (100 mL), 1-adamantyl isocyanate (0.782 g, FW = 177.24, 97%, 4.28 mmol) was added. The reaction mixture was heated at 80 ˚C. The reaction was monitored by TLC until all the isocyanate is consumed. The reaction mixture was then cooled down to room temperature and the solid product was filtered. The solid paste was further triturated and washed with hexane (100 mL). The isolated solid product was dried in vacuum overnight to afford final compound as a white solid in 69 % yield (1.2 g, FW = 392.58, 3.1 mmol)

$^{1}$H NMR (500 MHz, dmso-$d_6$): δ 11.97 (br, 1H), 5.59 (t, *J = 5 Hz*, 1H), 5.43 (s, 1H), 2.89 (q, *J = 5 Hz*, 2H), 2.18 (t, *J = 5 Hz*, 2H), 1.95-2.00 (m, 3H), 1.80-1.86 (m, 6H), 1.54-1.63 (m, 6H), 1.43-1.52 (m, 2H), 1.25-1.35 (m, 2H), 1.17-1.27 (m, 14H)

$^{13}$C NMR (125 MHz, dmso-$d_6$): δ 174.53, 157.06, 49.32, 42.07, 38.76, 36.17, 33.69, 30.06, 29.08, 28.99, 28.97, 28.84, 28.81, 28.60, 26.46, 24.54

### 3. Appendix: Characterization of Synthesized molecules

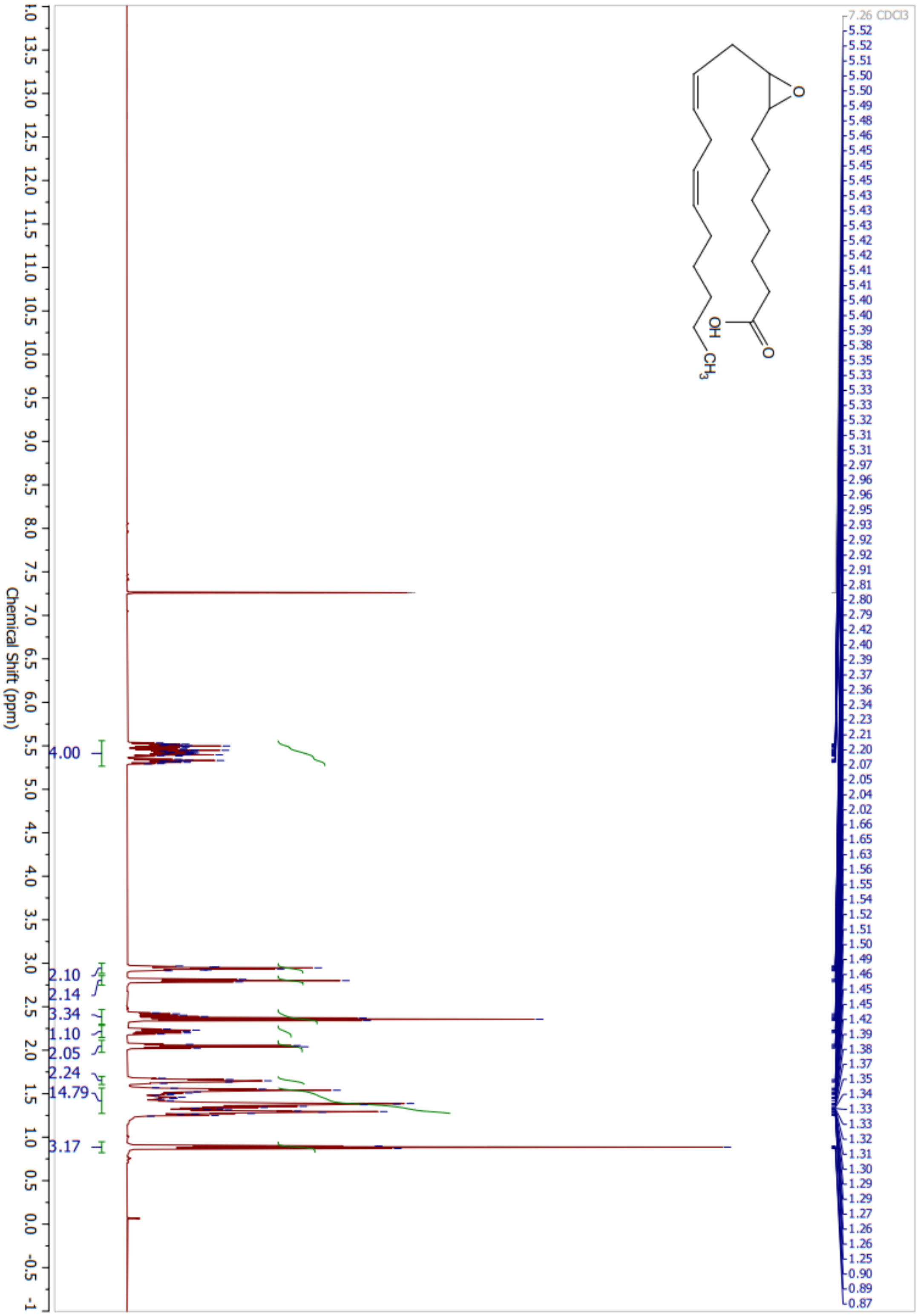

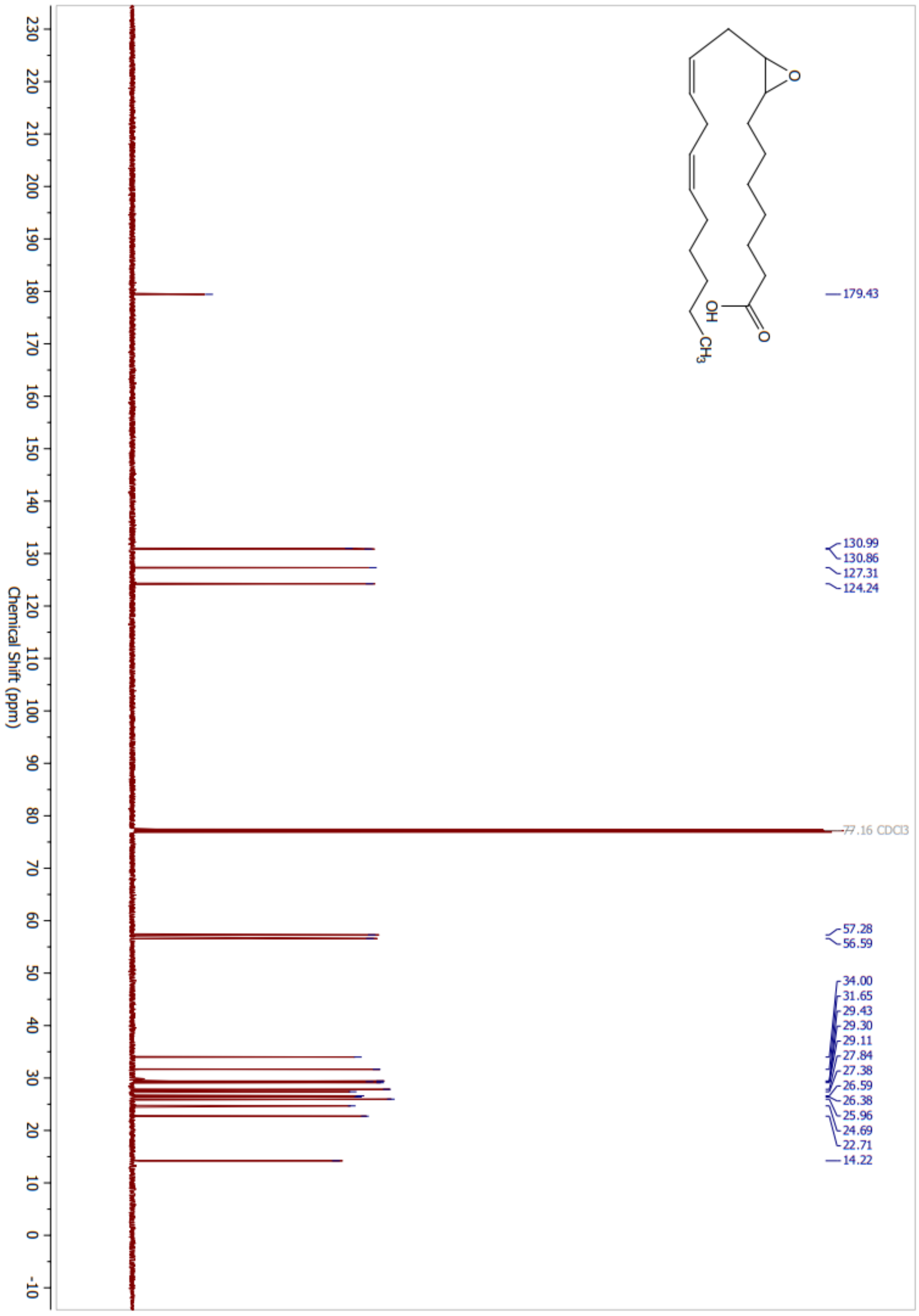

O
OH
O
CH3
179.43
130.99
130.86
127.31
124.24
77.16 CDCl3
57.28
56.59
34.00
31.65
29.43
29.30
29.11
27.84
27.38
26.59
26.38
25.96
24.69
22.71
14.22
Chemical Shift (ppm)

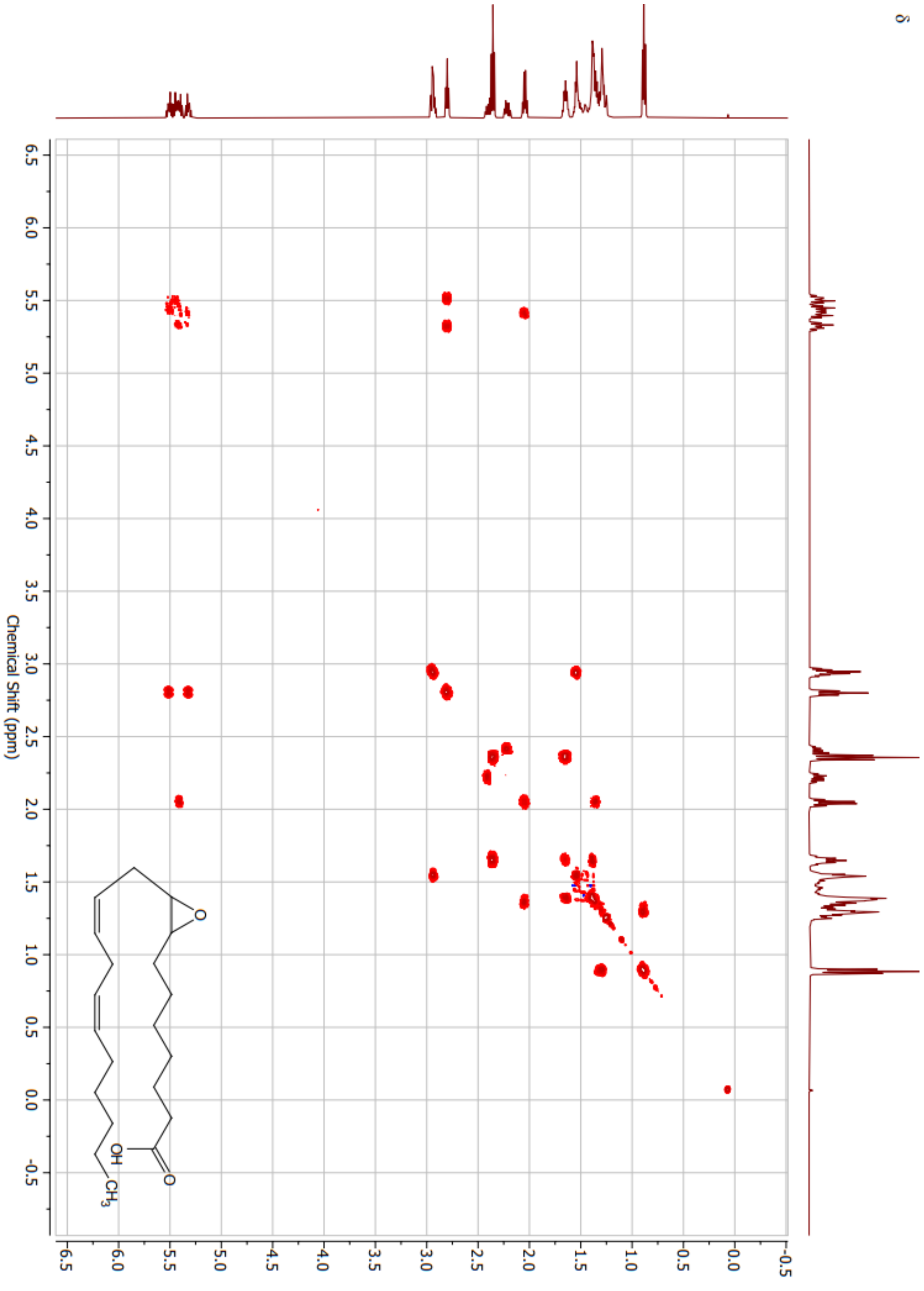
Chemical Shift (ppm)
δ
O
OH
CH3

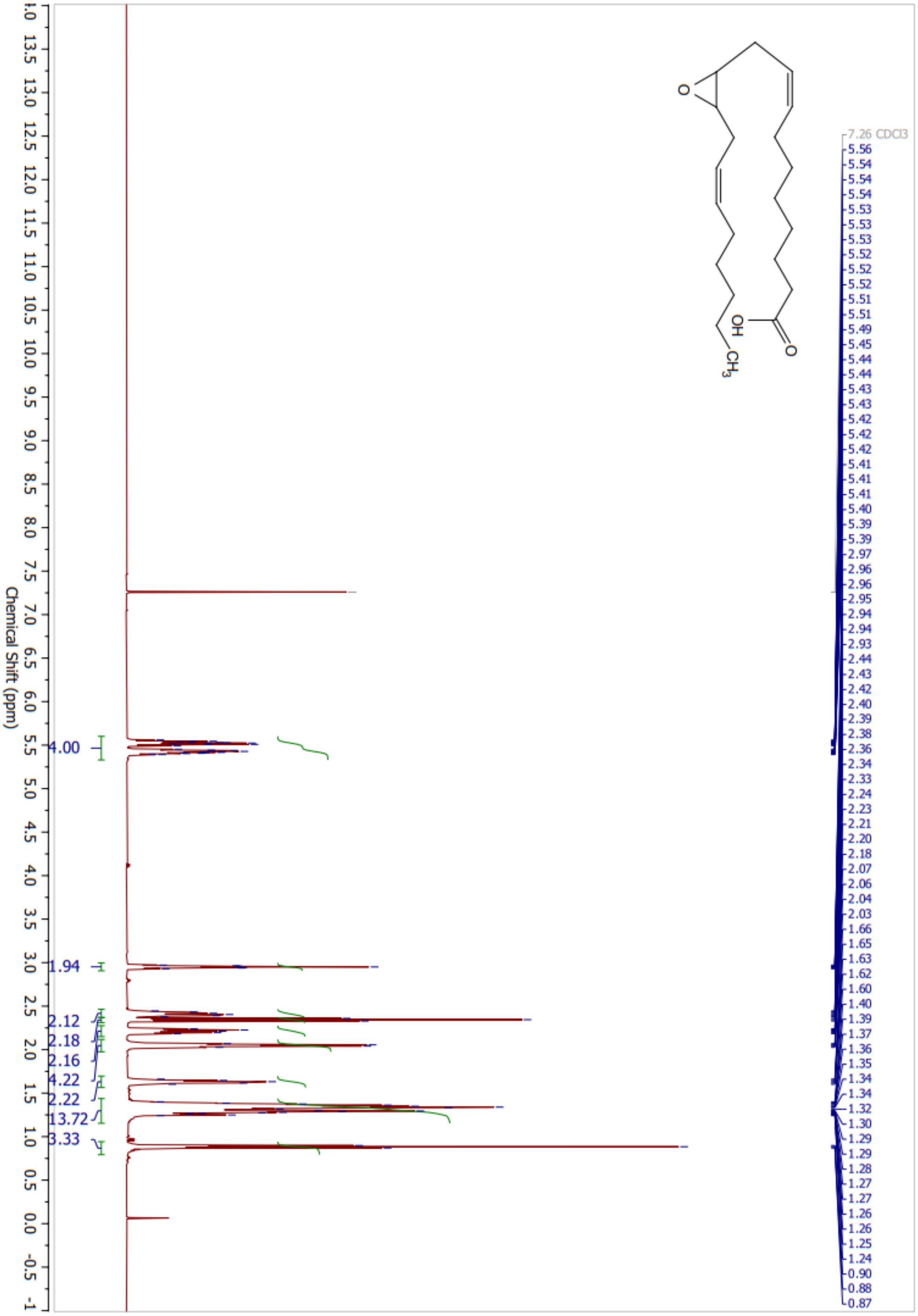
O
OH
O
CH3
7.26 CDCl3
5.56
5.54
5.54
5.54
5.53
5.53
5.53
5.52
5.52
5.52
5.51
5.51
5.49
5.45
5.44
5.44
5.43
5.43
5.42
5.42
5.42
5.41
5.41
5.41
5.40
5.39
5.39
2.97
2.96
2.96
2.95
2.94
2.94
2.93
2.44
2.43
2.42
2.40
2.39
2.38
2.36
2.34
2.33
2.24
2.23
2.21
2.20
2.18
2.07
2.06
2.04
2.03
1.66
1.65
1.63
1.62
1.60
1.40
1.39
1.37
1.36
1.35
1.34
1.34
1.32
1.30
1.29
1.29
1.28
1.27
1.27
1.26
1.26
1.25
1.24
0.90
0.88
0.87
4.00
1.94
2.12
2.18
2.16
4.22
2.22
13.72
3.33
13.5 13.0 12.5 12.0 11.5 11.0 10.5 10.0 9.5 9.0 8.5 8.0 7.5 7.0 6.5 6.0 5.5 5.0 4.5 4.0 3.5 3.0 2.5 2.0 1.5 1.0 0.5 0.0 -0.5 -1
Chemical Shift (ppm)

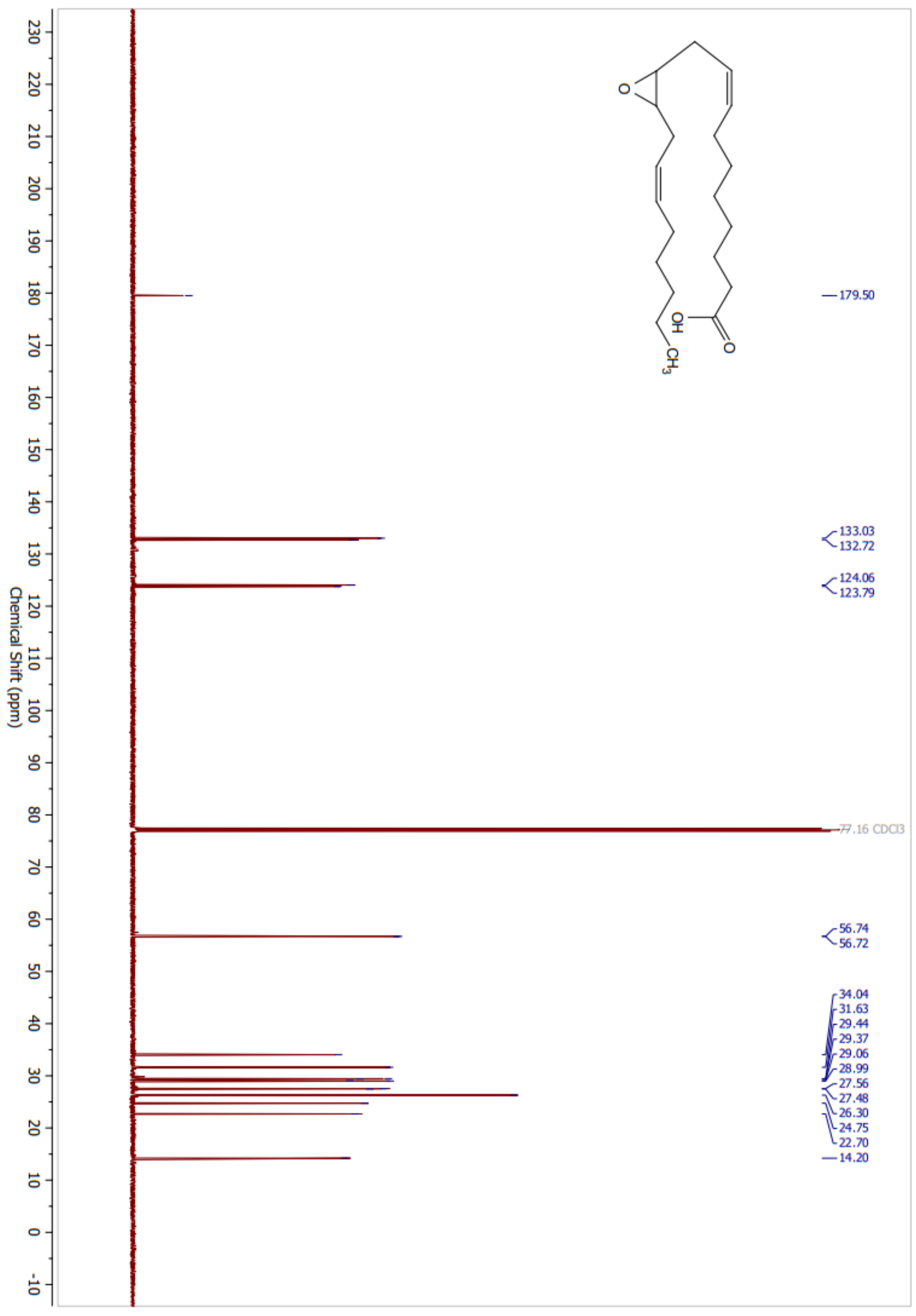

179.50
133.03
132.72
124.06
123.79
77.16 CDCl3
56.74
56.72
34.04
31.63
29.44
29.37
29.06
28.99
27.56
27.48
26.30
24.75
22.70
14.20
OH
O
CH3
Chemical Shift (ppm)

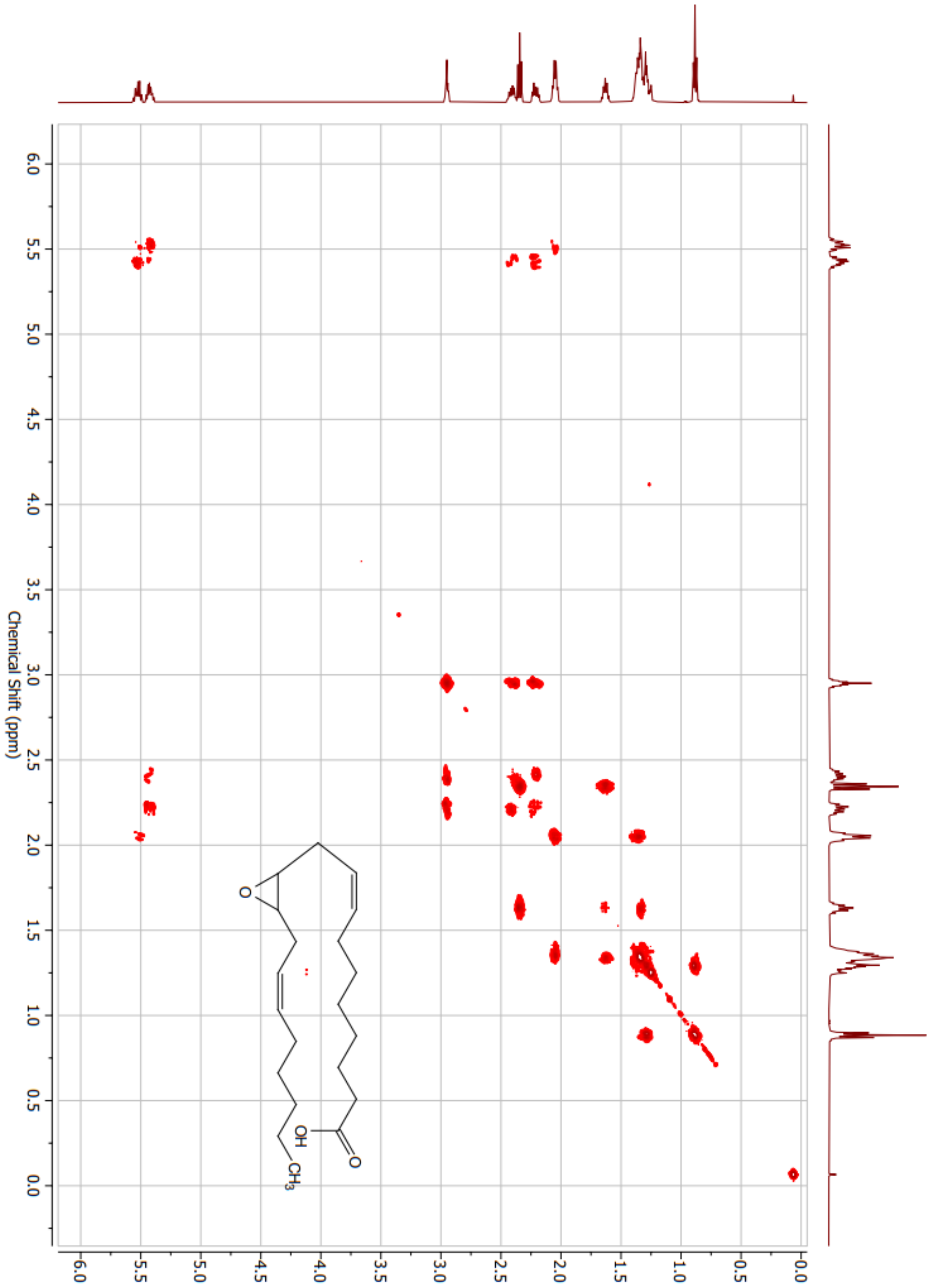
Chemical Shift (ppm)
OH
O
O
CH3

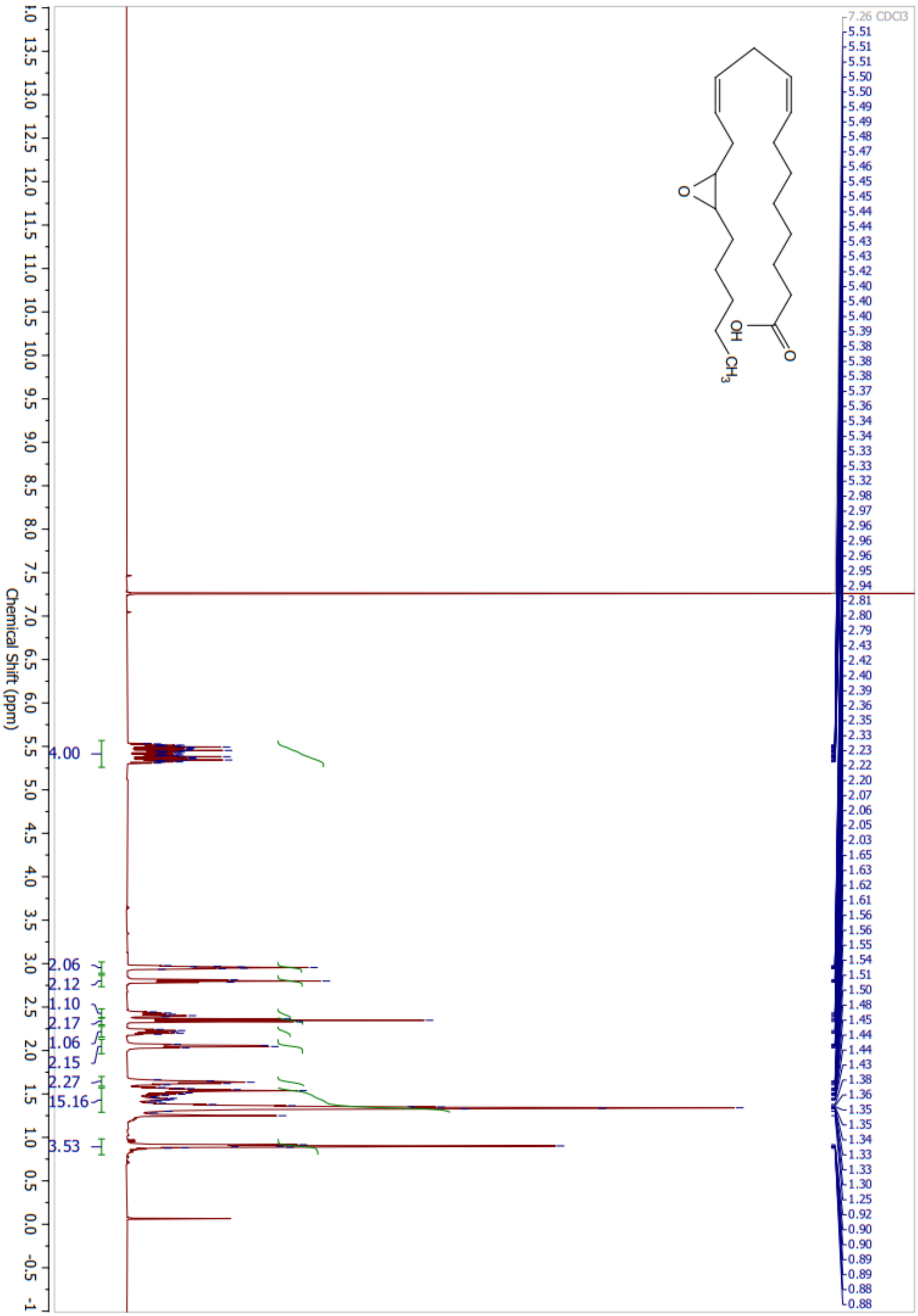
7.26 CDCl3
Chemical Shift (ppm)
4.00
2.06
2.12
1.10
2.17
1.06
2.15
2.27
15.16
3.53
OH
O
CH3

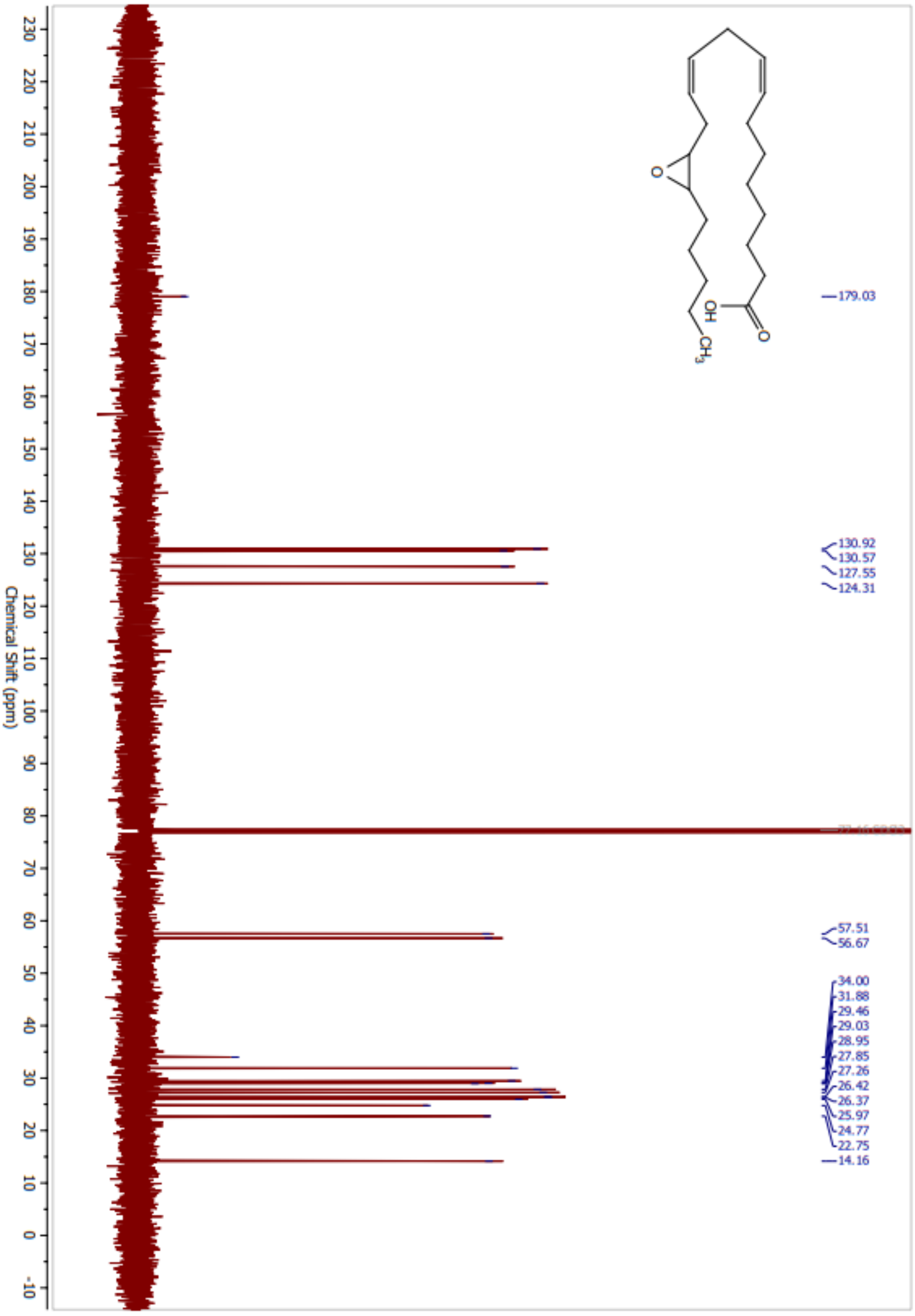
179.03
130.92
130.57
127.55
124.31
57.51
56.67
34.00
31.88
29.46
29.03
28.95
27.85
27.26
26.42
26.37
25.97
24.77
22.75
14.16
Chemical Shift (ppm)
OH
O
CH3

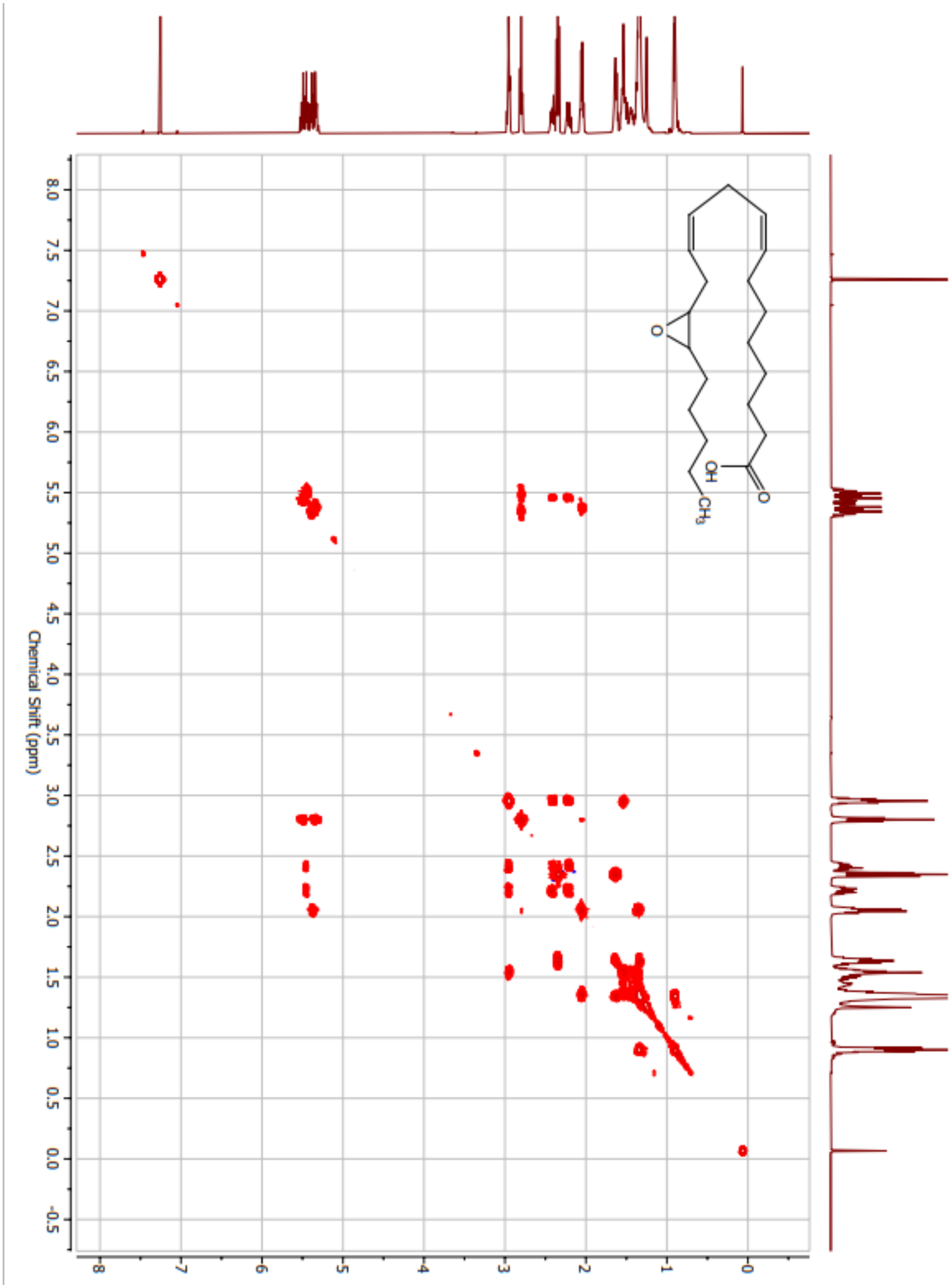
Chemical Shift (ppm)
OH
O
CH3

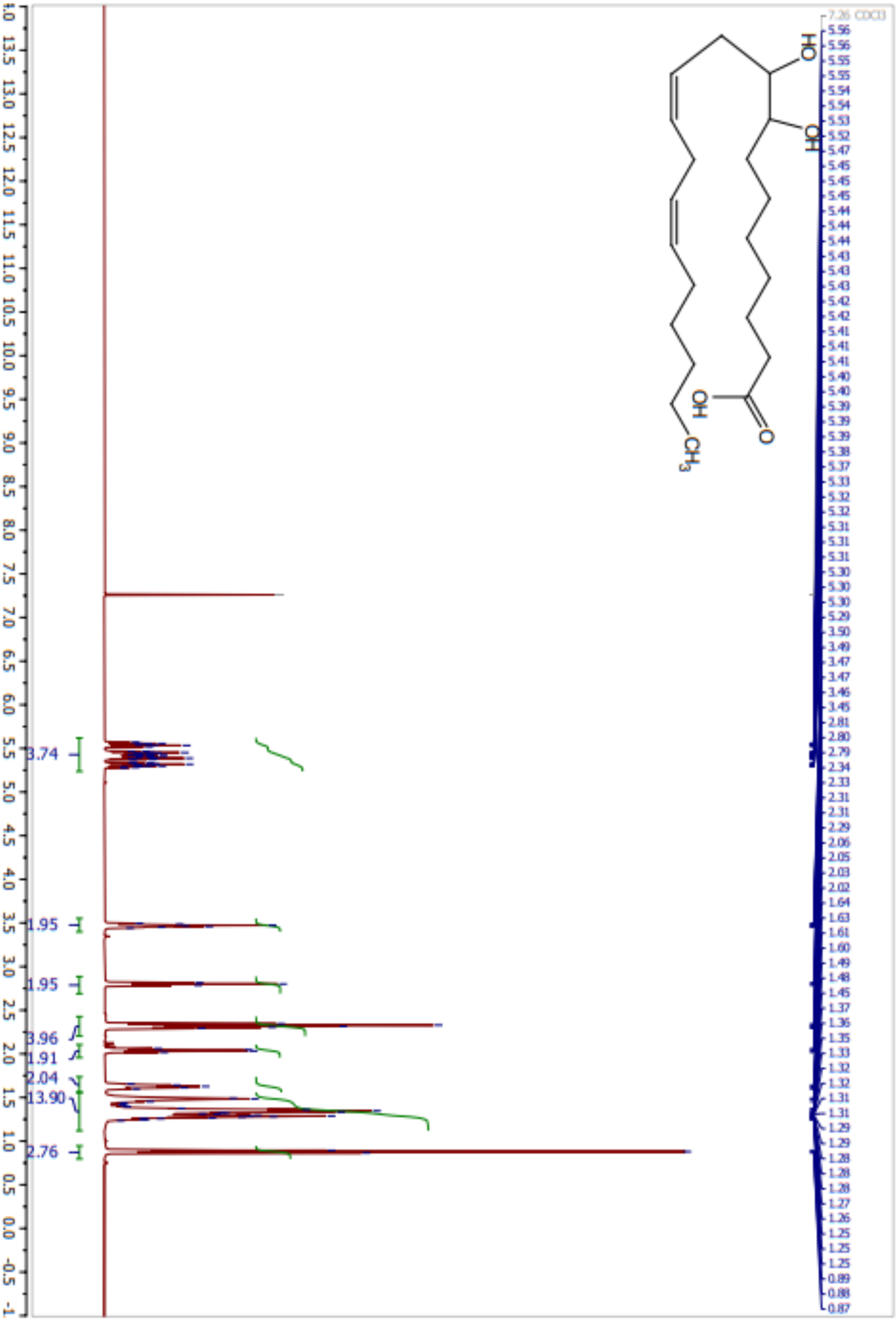
Chemical Shift (ppm)
7.26 CDCl3
3.74
1.95
1.95
3.96
1.91
2.04
13.90
2.76

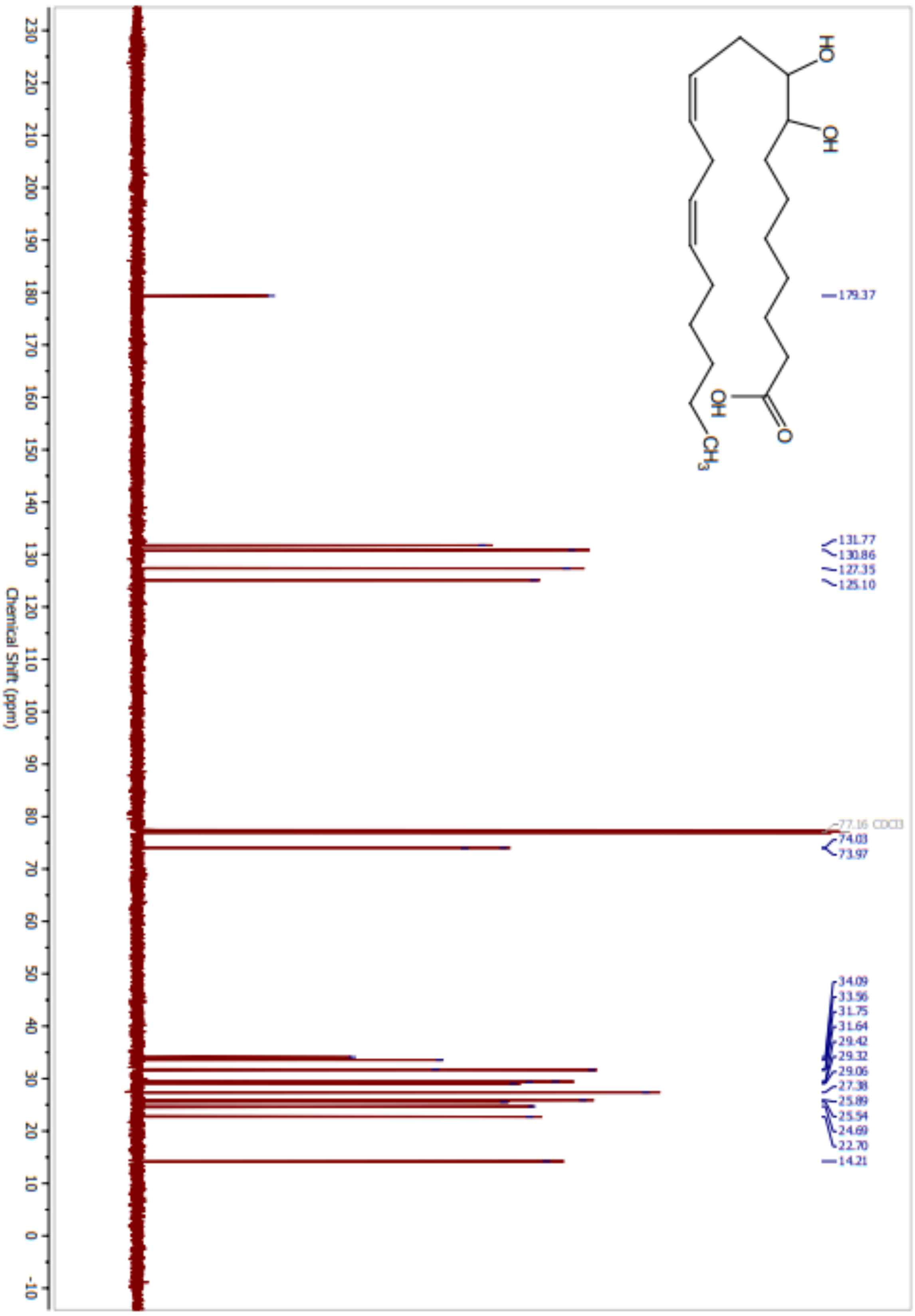
HO
OH
OH
O
CH3
179.37
131.77
130.86
127.35
125.10
77.16 CDCl3
74.03
73.97
34.09
33.56
31.75
31.64
29.42
29.32
29.06
27.38
25.89
25.54
24.69
22.70
14.21
230 220 210 200 190 180 170 160 150 140 130 120 110 100 90 80 70 60 50 40 30 20 10 0 -10
Chemical Shift (ppm)

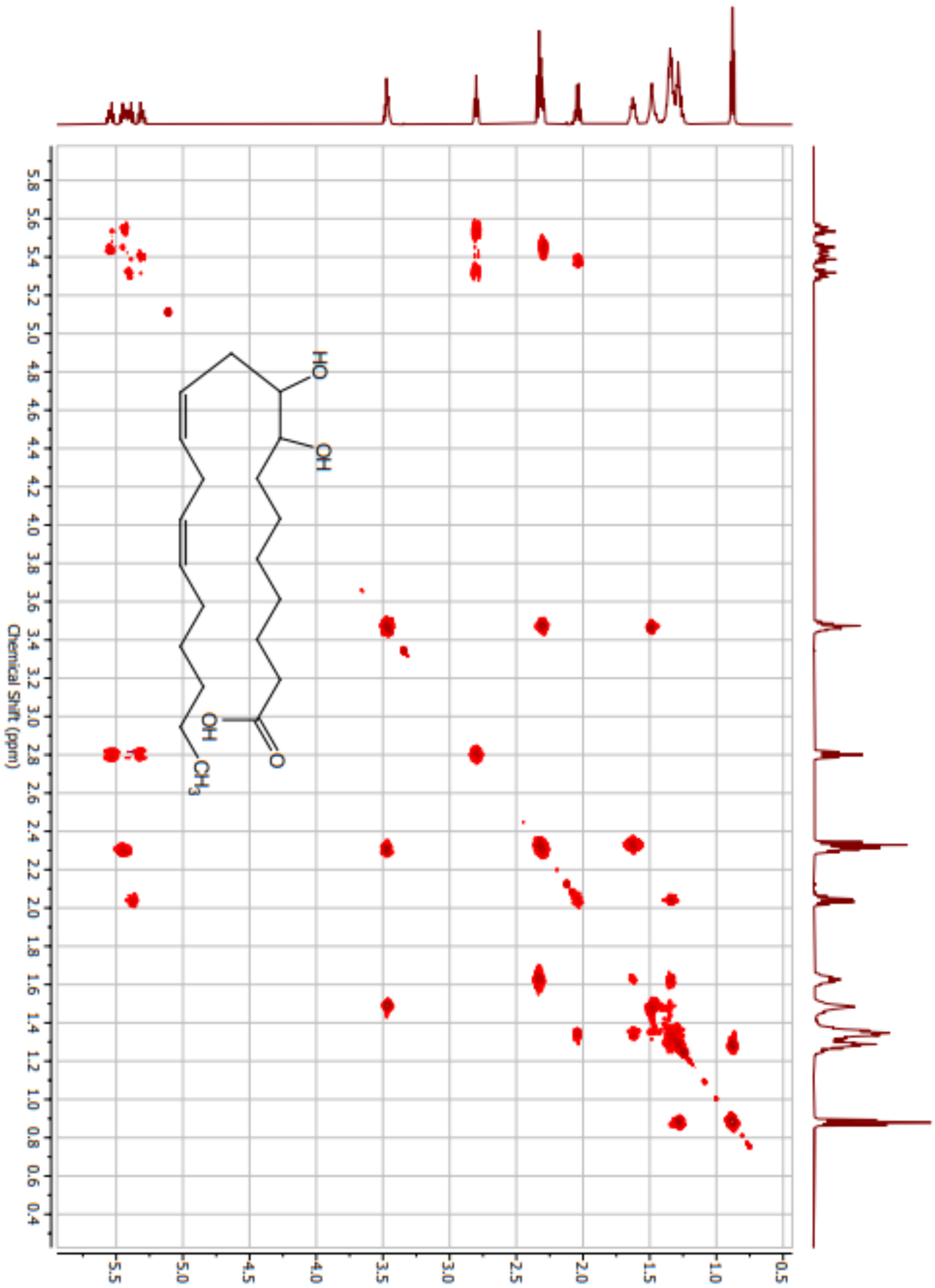
Chemical Shift (ppm)
HO
OH
OH
O
CH3

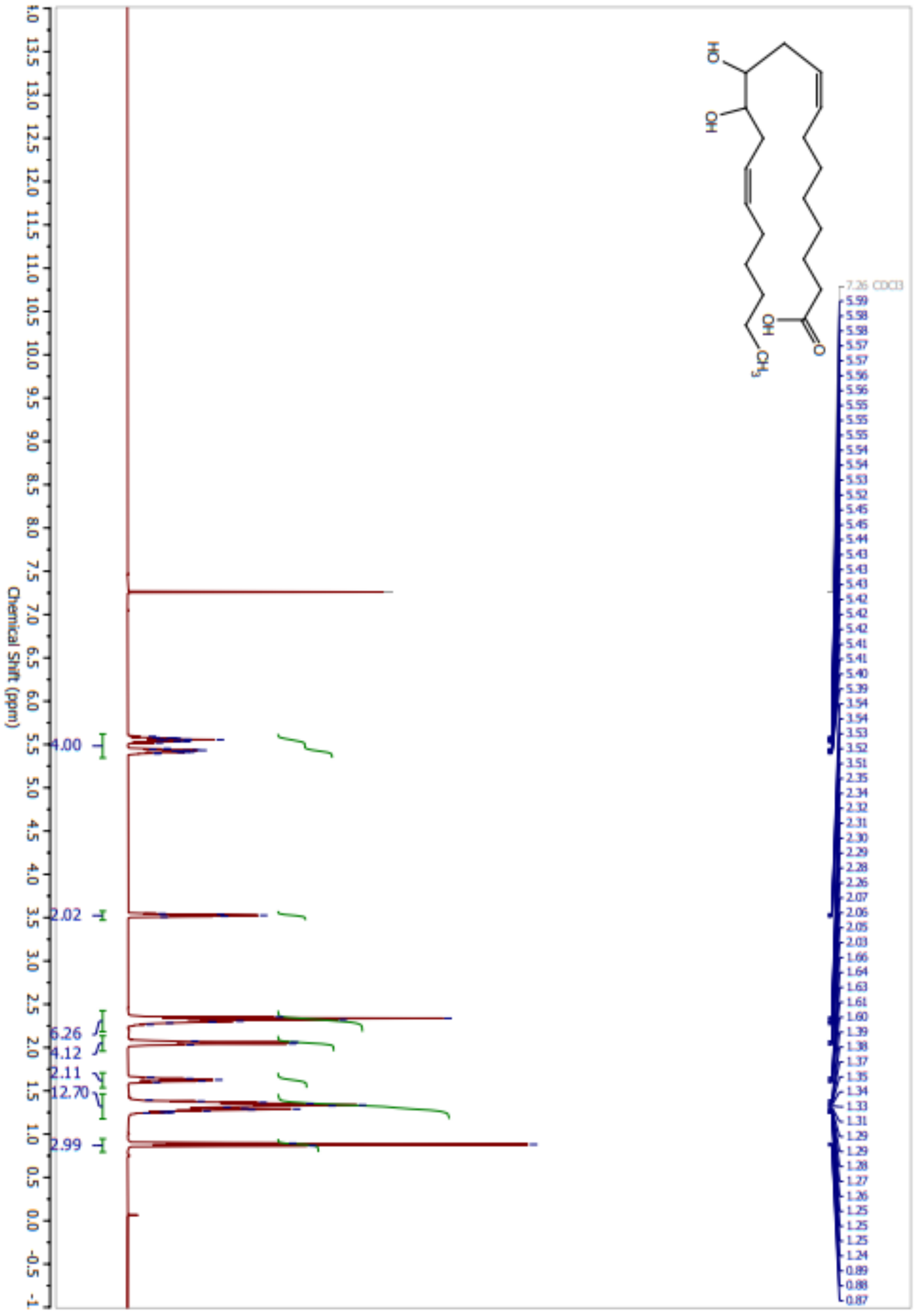
Chemical Shift (ppm)
7.26 CDCl3
4.00
2.02
6.26
4.12
2.11
12.70
2.99

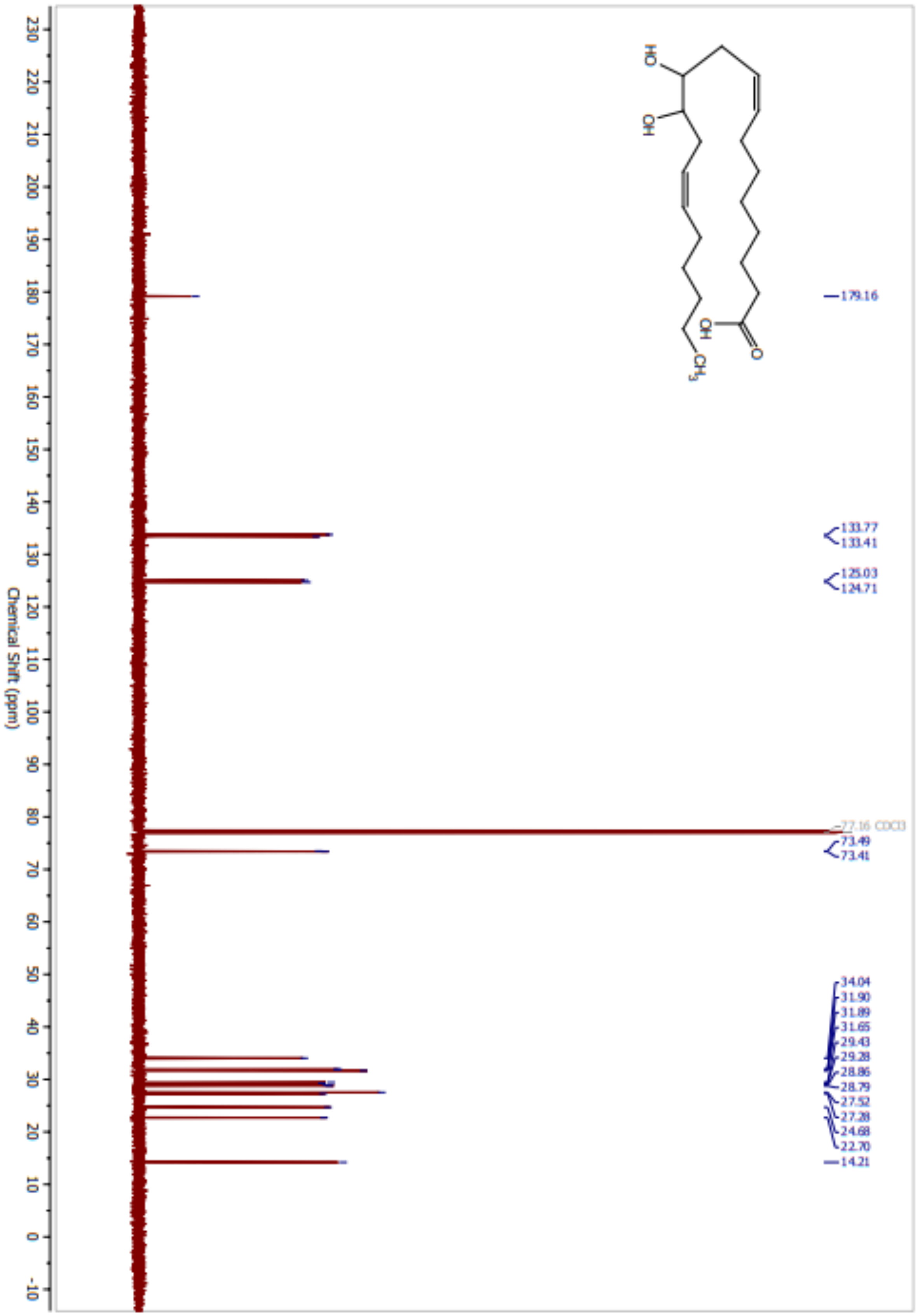
Chemical Shift (ppm)
179.16
133.77
133.41
125.03
124.71
77.16 CDCl3
73.49
73.41
34.04
31.90
31.89
31.65
29.43
29.28
28.86
28.79
27.52
27.28
24.68
22.70
14.21
OH
HO
OH
O
CH3

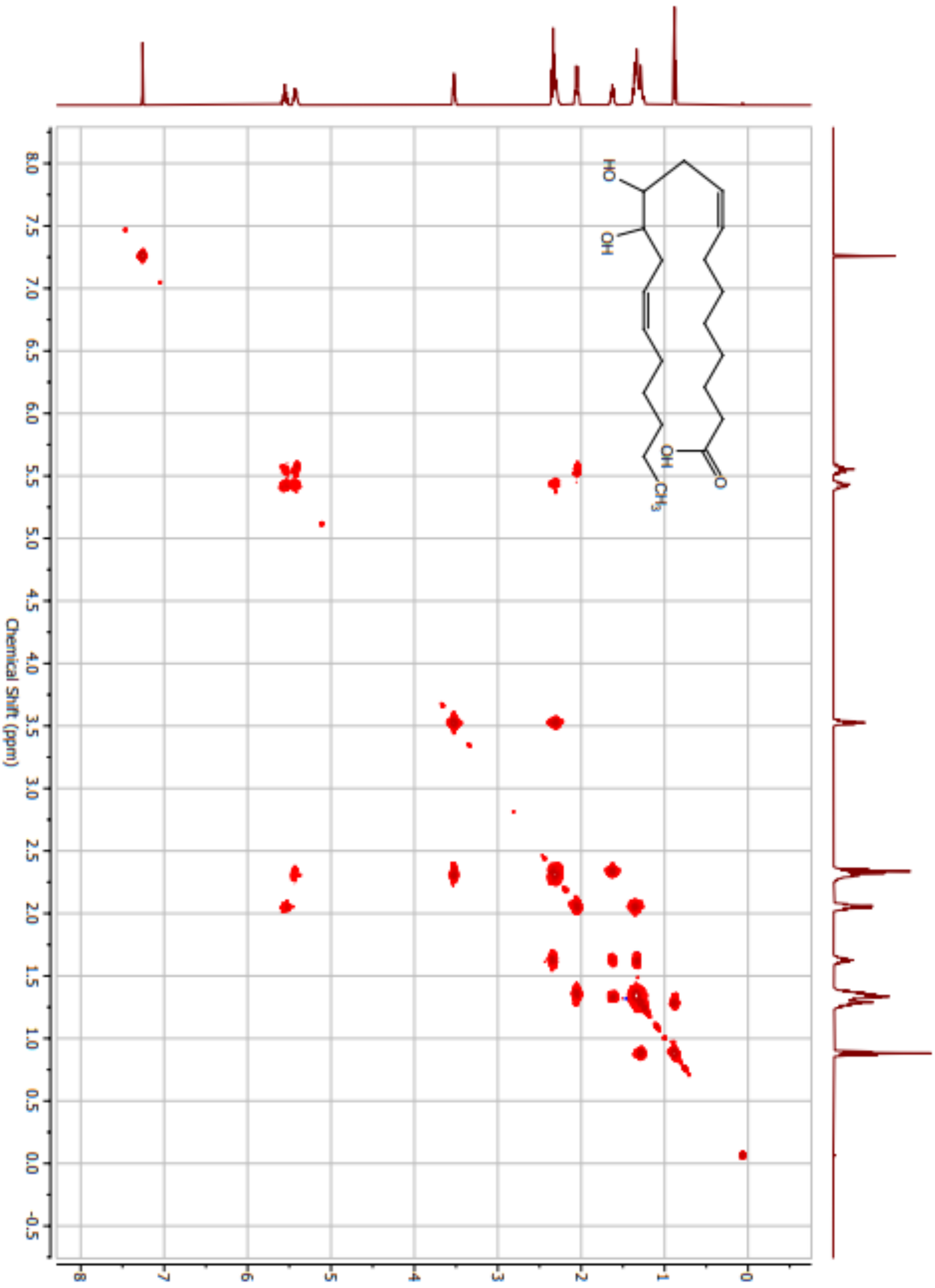
Chemical Shift (ppm)
HO
OH
OH
O
CH3

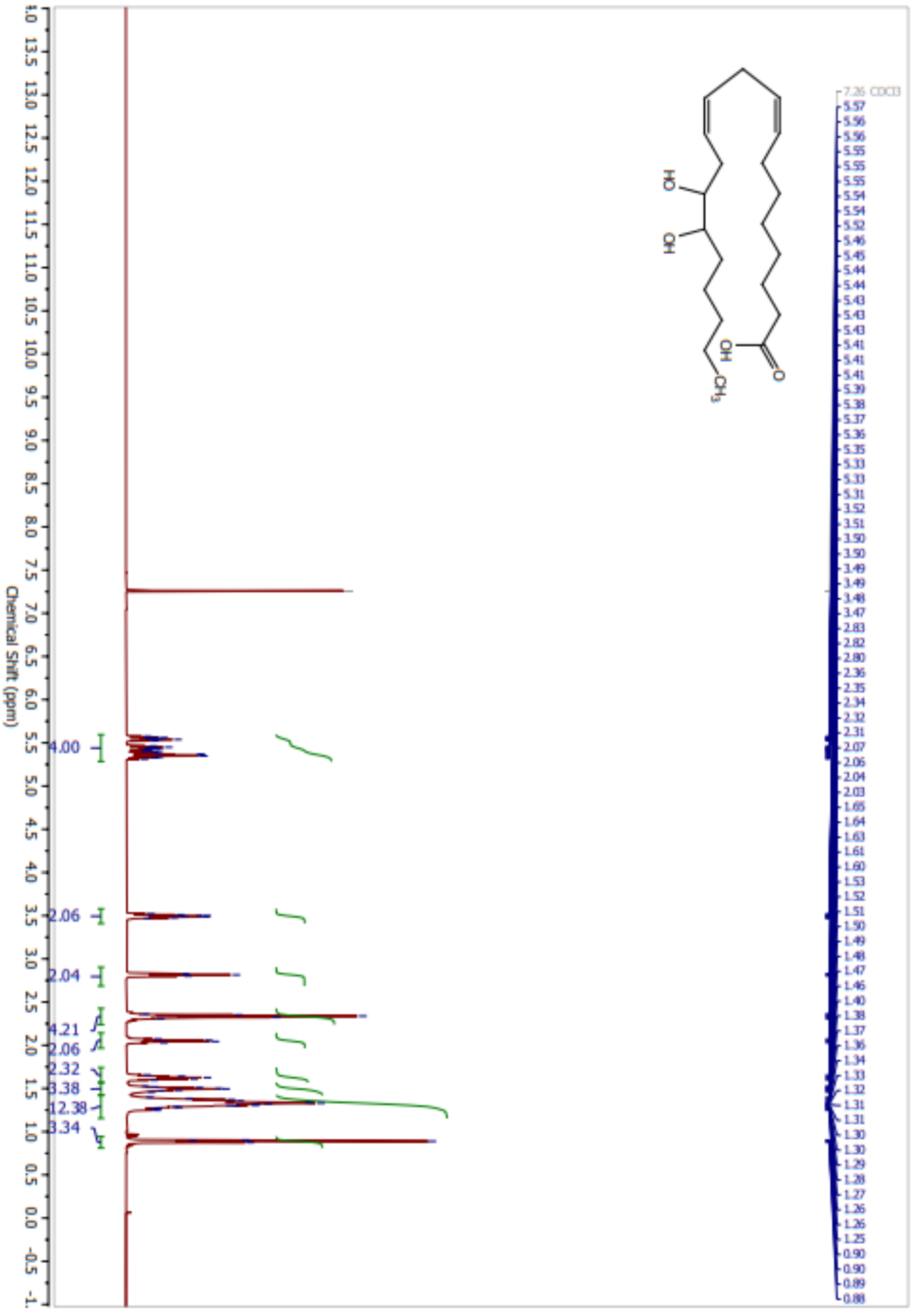

Chemical Shift (ppm)
4.00
2.06
2.04
4.21
2.06
2.32
3.38
12.38
3.34
OH
HO
OH
O
CH3
7.26 CDCl3
5.57
5.56
5.56
5.55
5.55
5.55
5.54
5.54
5.52
5.46
5.45
5.44
5.44
5.43
5.43
5.43
5.41
5.41
5.41
5.39
5.38
5.37
5.36
5.35
5.33
5.33
5.31
3.52
3.51
3.50
3.50
3.49
3.49
3.48
3.47
2.83
2.82
2.80
2.36
2.35
2.34
2.32
2.31
2.07
2.06
2.04
2.03
1.65
1.64
1.63
1.61
1.60
1.53
1.52
1.51
1.50
1.49
1.48
1.47
1.46
1.40
1.38
1.37
1.36
1.34
1.33
1.32
1.31
1.31
1.30
1.30
1.29
1.28
1.27
1.26
1.26
1.25
0.90
0.90
0.89
0.88

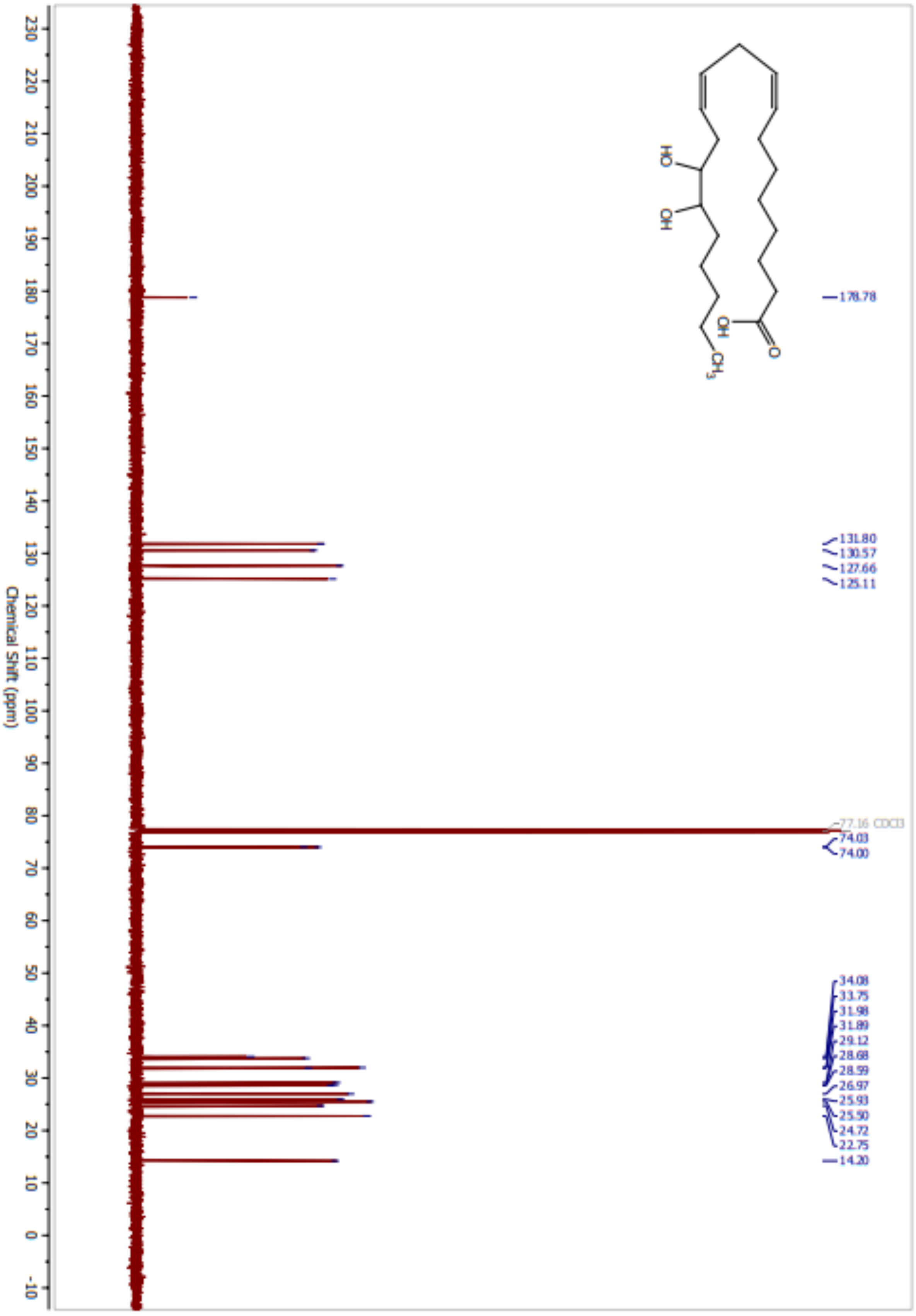

178.78
131.80
130.57
127.66
125.11
77.16 CDCl3
74.03
74.00
34.08
33.75
31.98
31.89
29.12
28.68
28.59
26.97
25.93
25.50
24.72
22.75
14.20
Chemical Shift (ppm)
OH
HO
OH
O
CH3

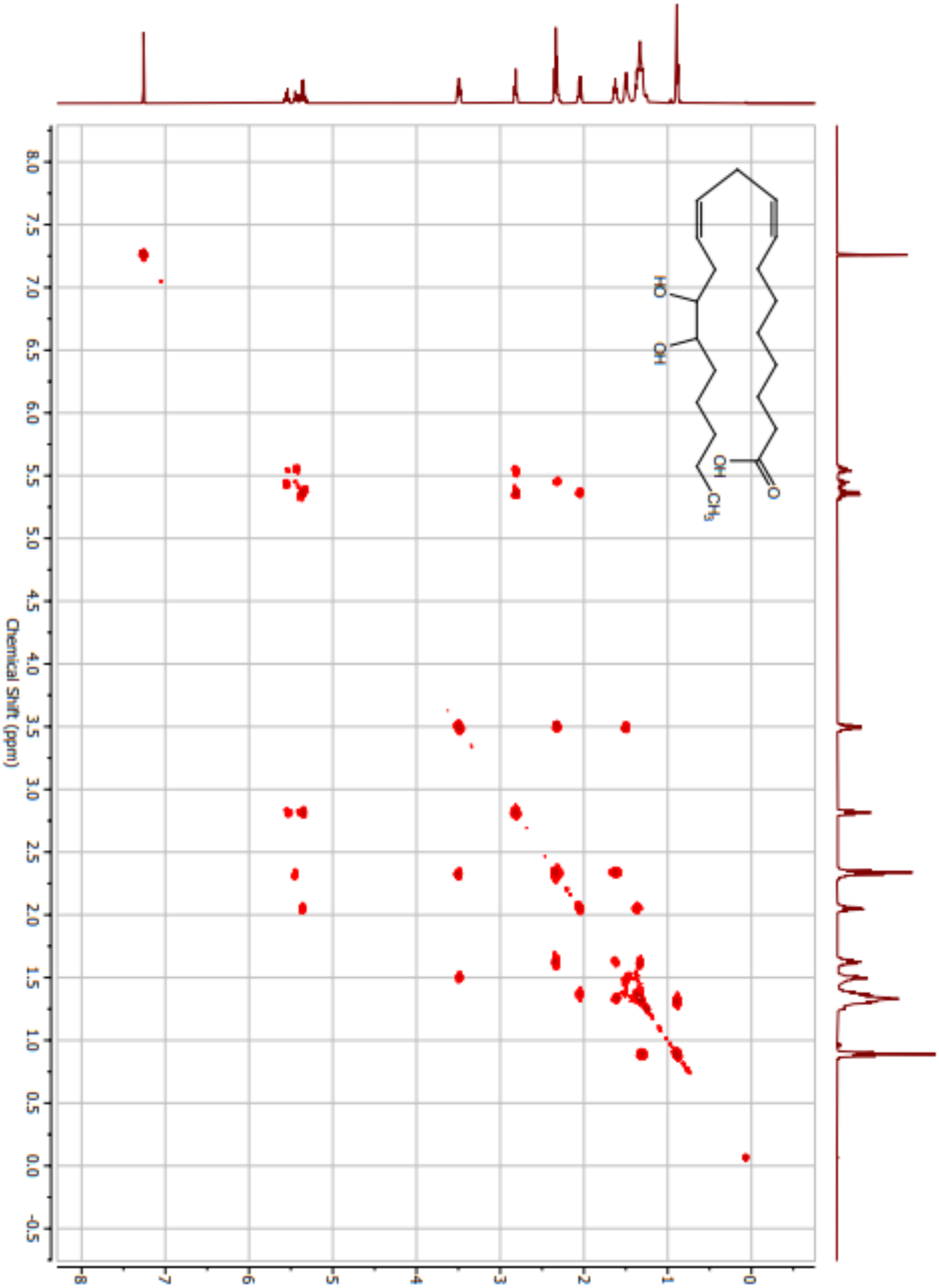
Chemical Shift (ppm)
HO
OH
OH
O
CH3

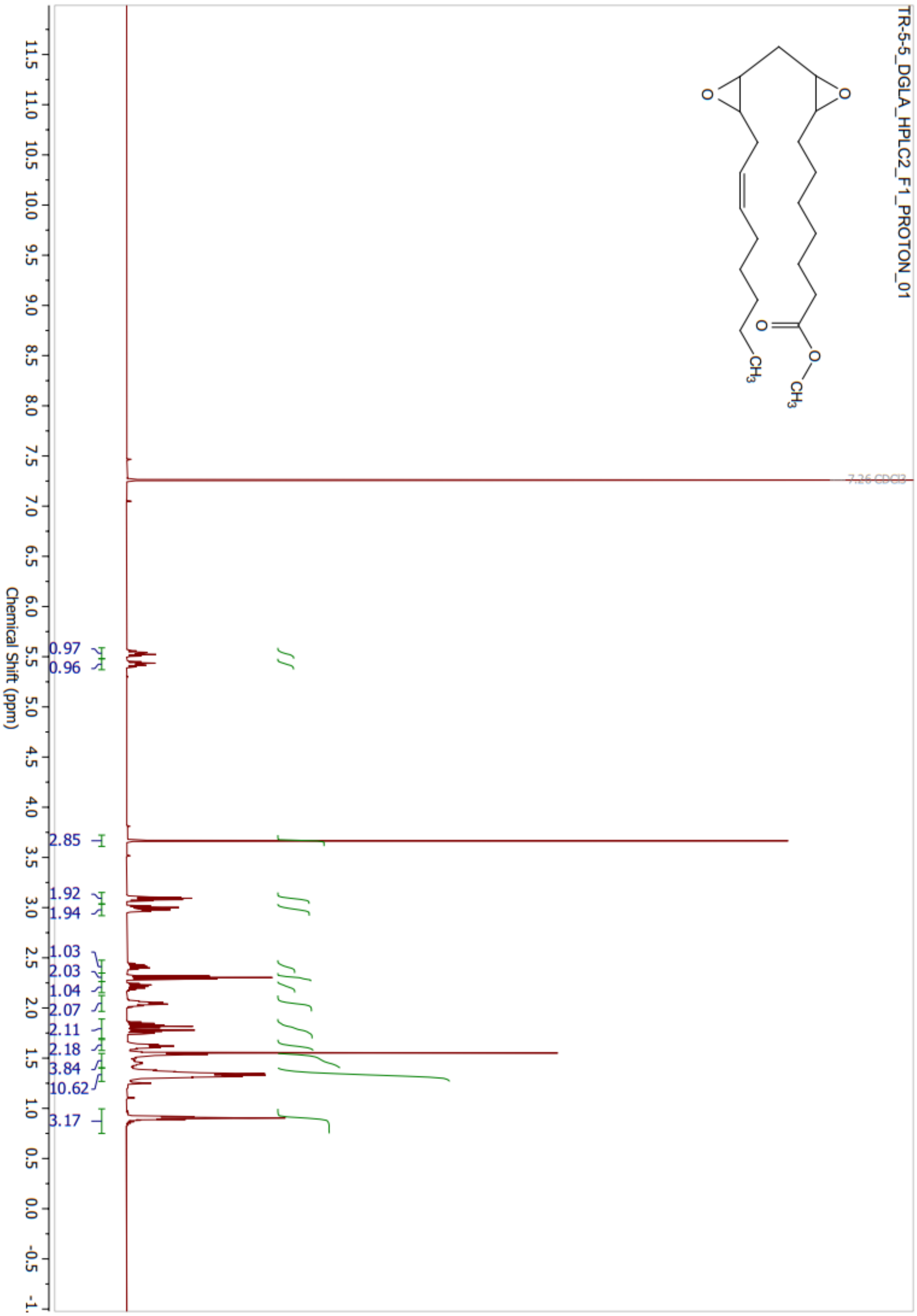
TR-5-5_DGLA_HPLC2_F1_PROTON_01
7.26 CDCl3
0.97
0.96
2.85
1.92
1.94
1.03
2.03
1.04
2.07
2.11
2.18
3.84
10.62
3.17
Chemical Shift (ppm)

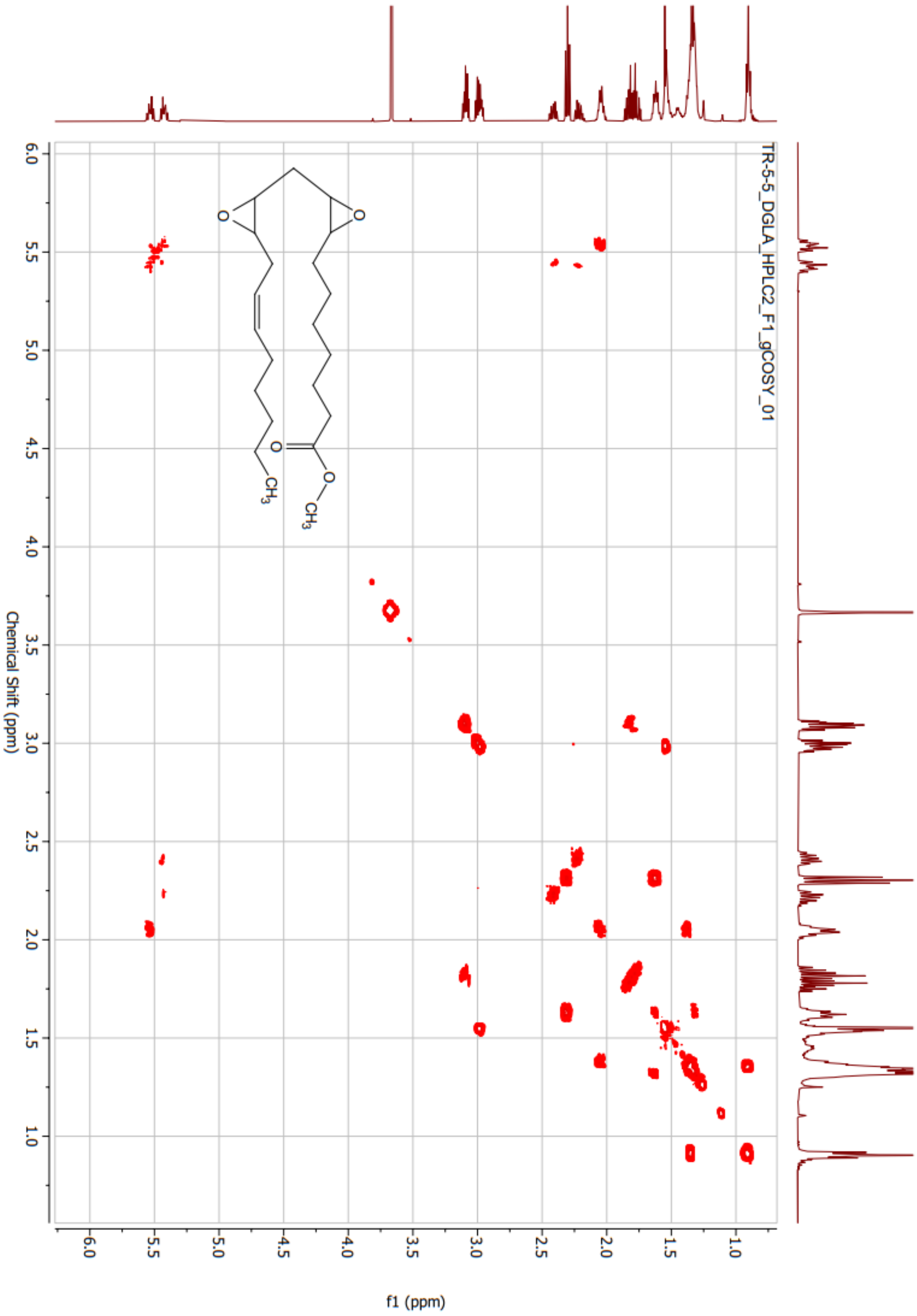
TR-5-5_DGLA_HPLC2_F1_gCOSY_01
Chemical Shift (ppm)
f1 (ppm)

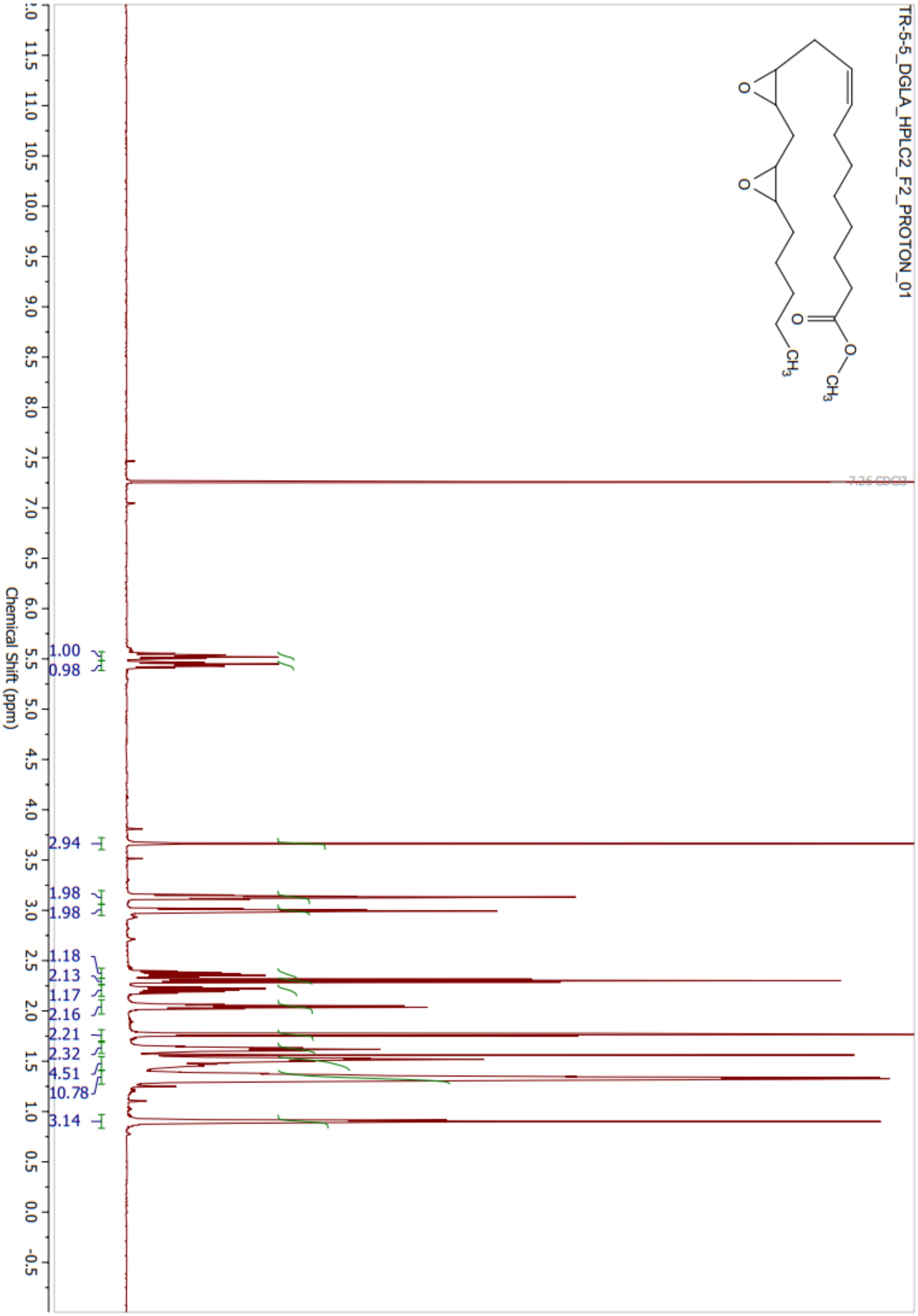
TR-5-5_DGLA_HPLC2_F2_PROTON_01
O
O
O
O
CH3
CH3
7.26 CDCl3
1.00
0.98
2.94
1.98
1.98
1.18
2.13
1.17
2.16
2.21
2.32
4.51
10.78
3.14
Chemical Shift (ppm)
12.0
11.5
11.0
10.5
10.0
9.5
9.0
8.5
8.0
7.5
7.0
6.5
6.0
5.5
5.0
4.5
4.0
3.5
3.0
2.5
2.0
1.5
1.0
0.5
0.0
-0.5

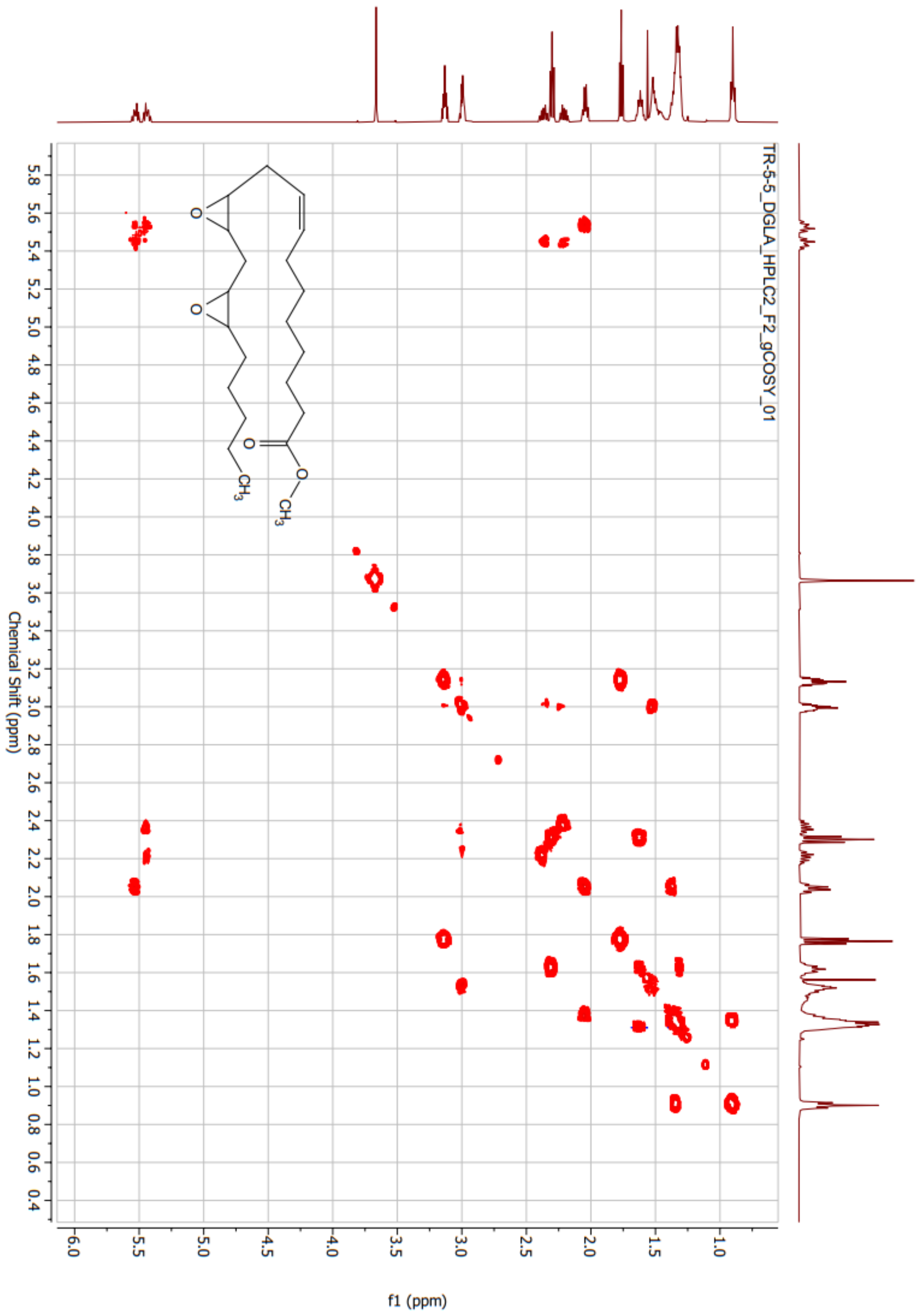
TR-5-5_DGLA_HPLC2_F2_gCOSY_01
Chemical Shift (ppm)
f1 (ppm)

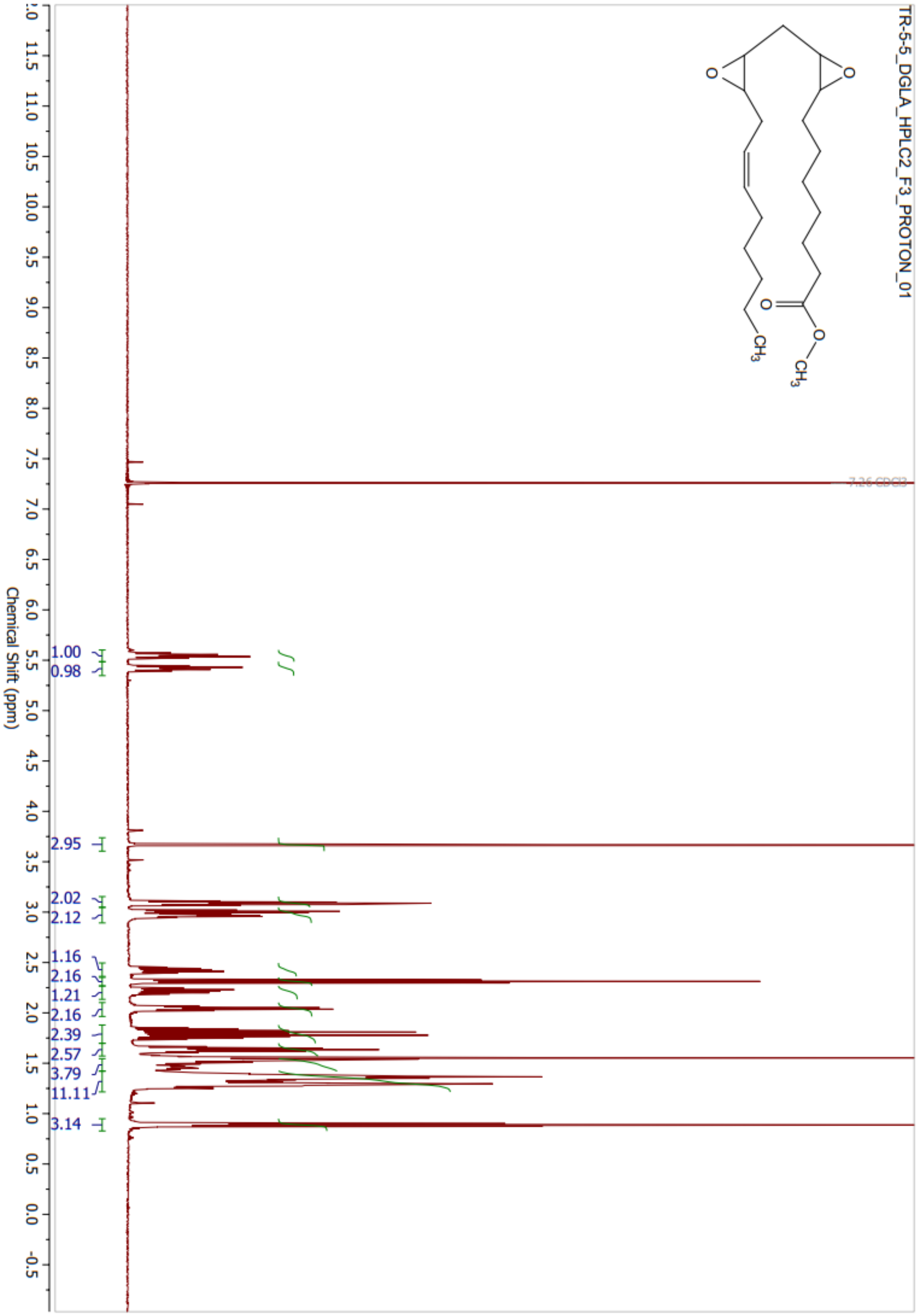
TR-5-5_DGLA_HPLC2_F3_PROTON_01
O
O
O
O
CH3
CH3
7.26 CDCl3
1.00
0.98
2.95
2.02
2.12
1.16
2.16
1.21
2.16
2.39
2.57
3.79
11.11
3.14
Chemical Shift (ppm)

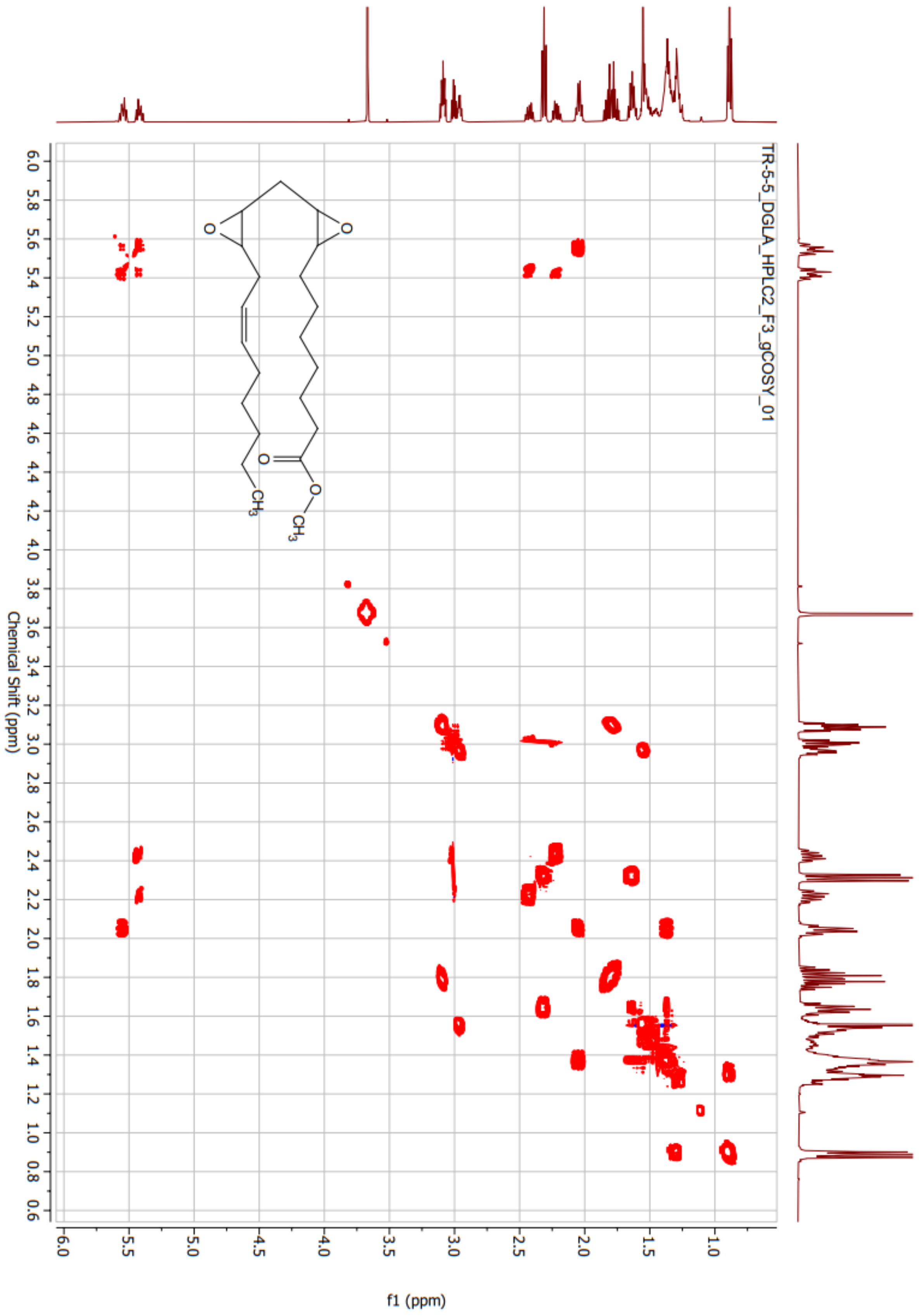
TR-5-5_DGLA_HPLC2_F3_gCOSY_01
Chemical Shift (ppm)
f1 (ppm)
O
O
O
O
CH3
CH3

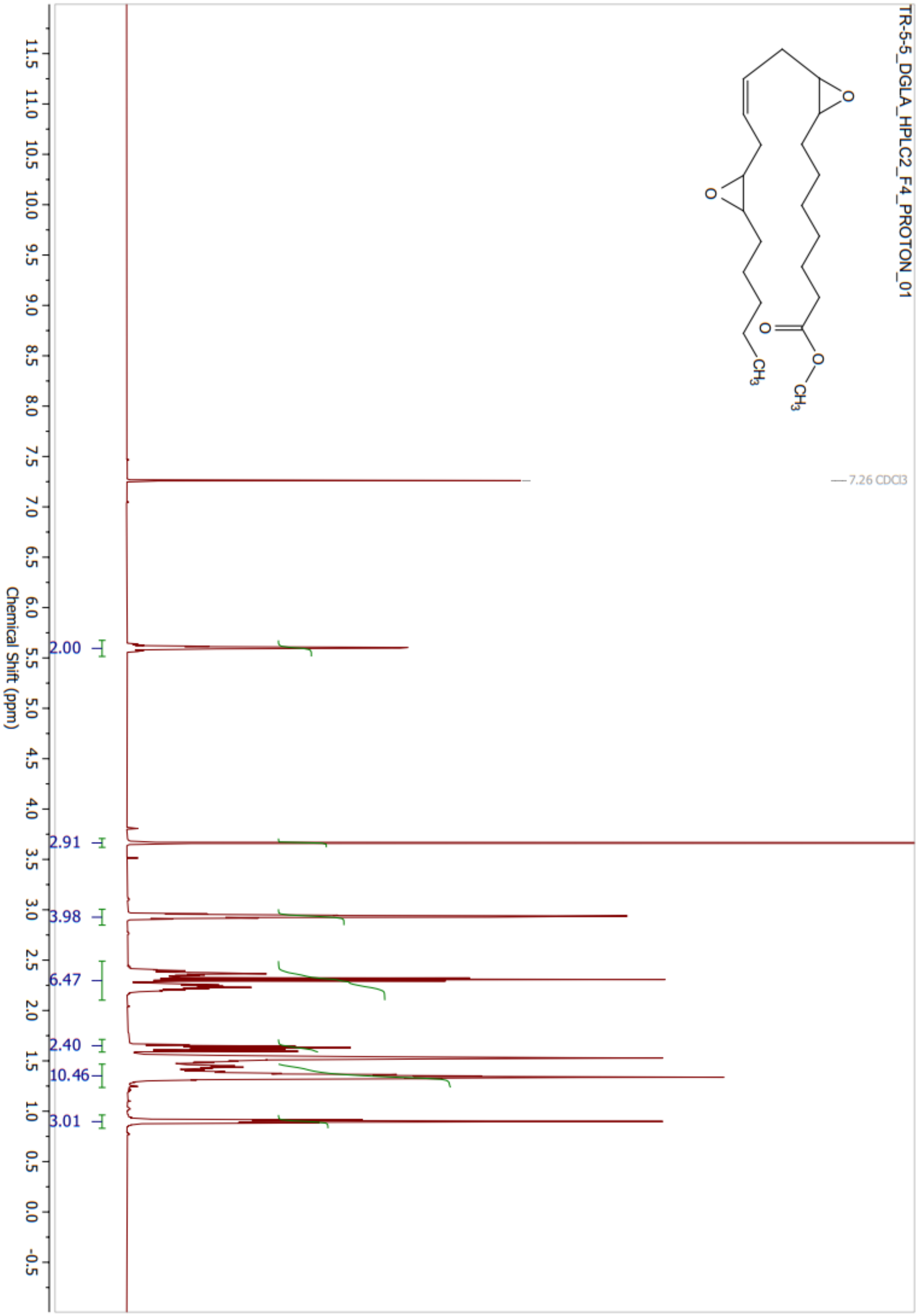

TR-5-5_DGLA_HPLC2_F4_PROTON_01
7.26 CDCl3
2.00
2.91
3.98
6.47
2.40
10.46
3.01
11.5 11.0 10.5 10.0 9.5 9.0 8.5 8.0 7.5 7.0 6.5 6.0 5.5 5.0 4.5 4.0 3.5 3.0 2.5 2.0 1.5 1.0 0.5 0.0 -0.5
Chemical Shift (ppm)

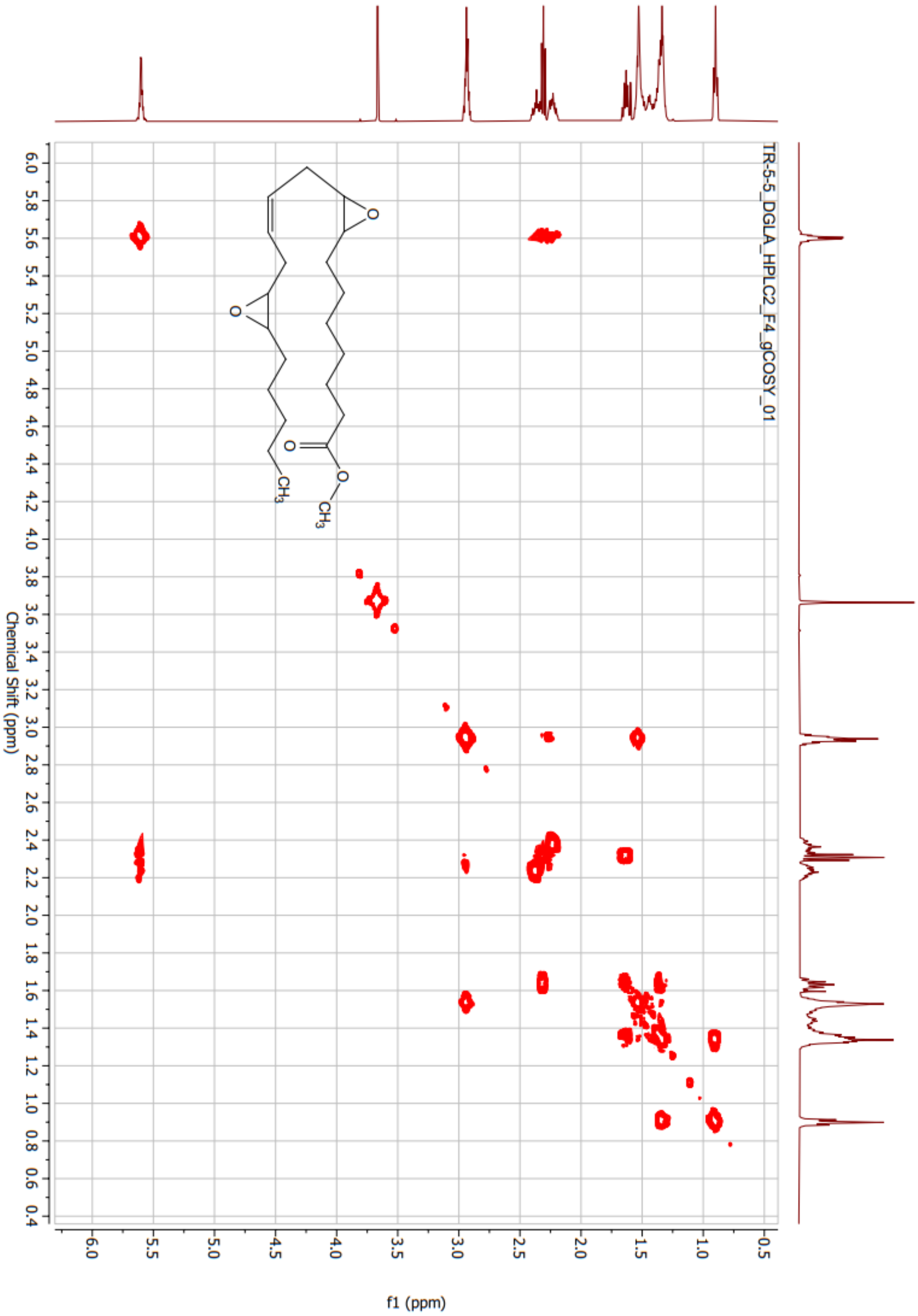
TR-5-5_DGLA_HPLC2_F4_gCOSY_01
Chemical Shift (ppm)
f1 (ppm)

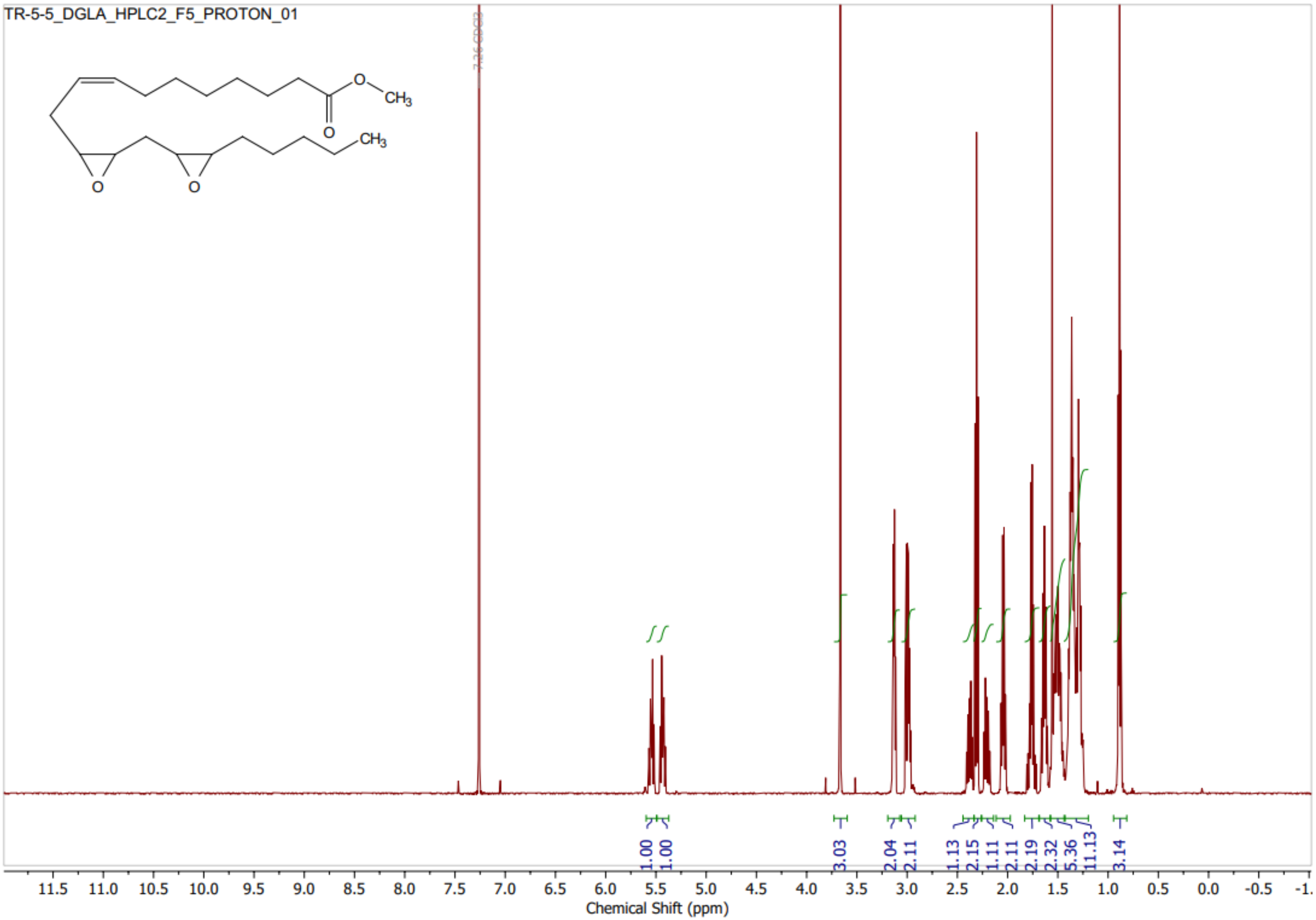
TR-5-5_DGLA_HPLC2_F5_PROTON_01
O
CH3
CH3
7.26 CDCl3
1.00
1.00
3.03
2.04
2.11
1.13
2.15
1.11
2.11
2.19
2.32
5.36
11.13
3.14
11.5 11.0 10.5 10.0 9.5 9.0 8.5 8.0 7.5 7.0 6.5 6.0 5.5 5.0 4.5 4.0 3.5 3.0 2.5 2.0 1.5 1.0 0.5 0.0 -0.5 -1.
Chemical Shift (ppm)

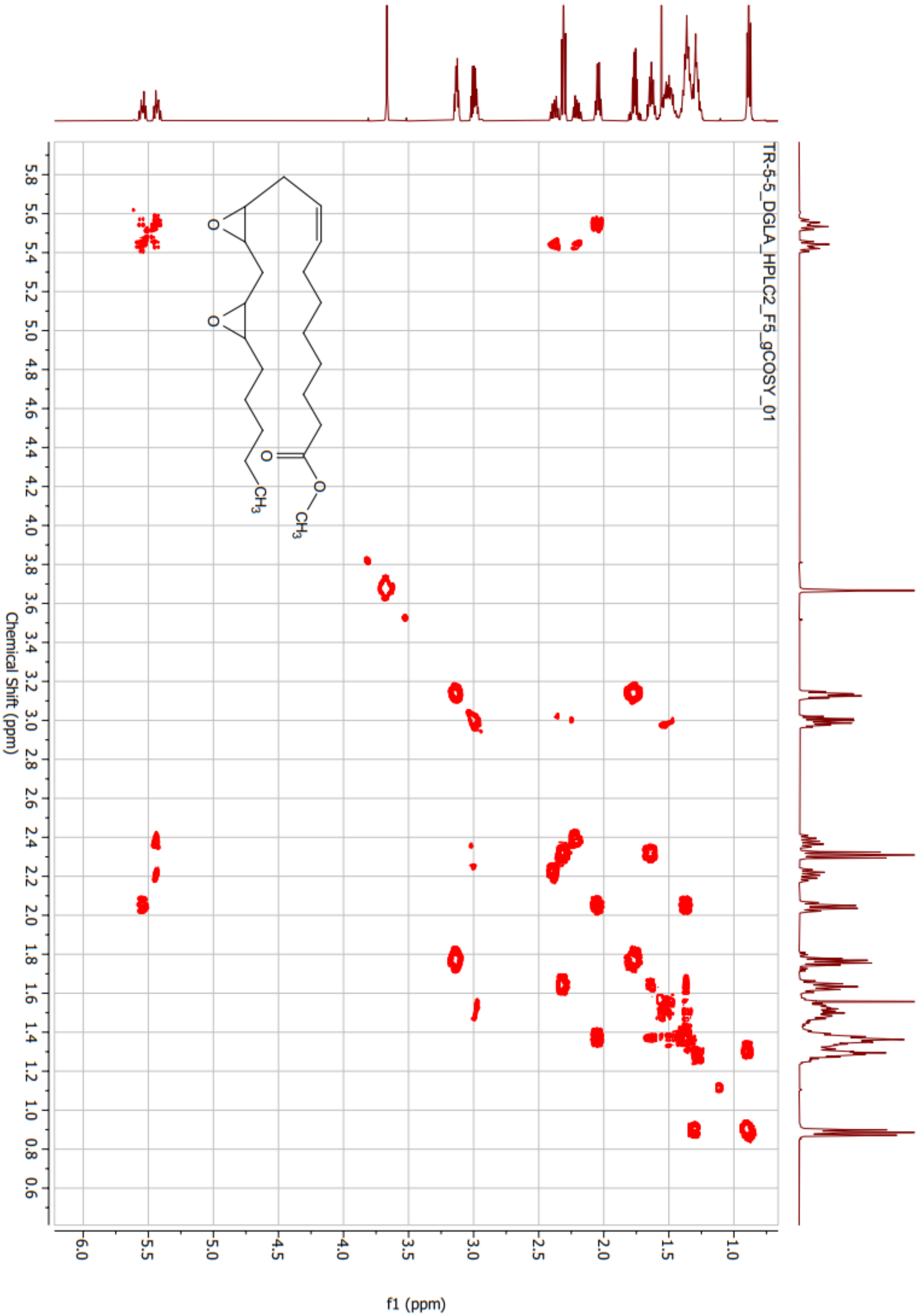
TR-5-5_DGLA_HPLC2_F5_gCOSY_01
Chemical Shift (ppm)
f1 (ppm)

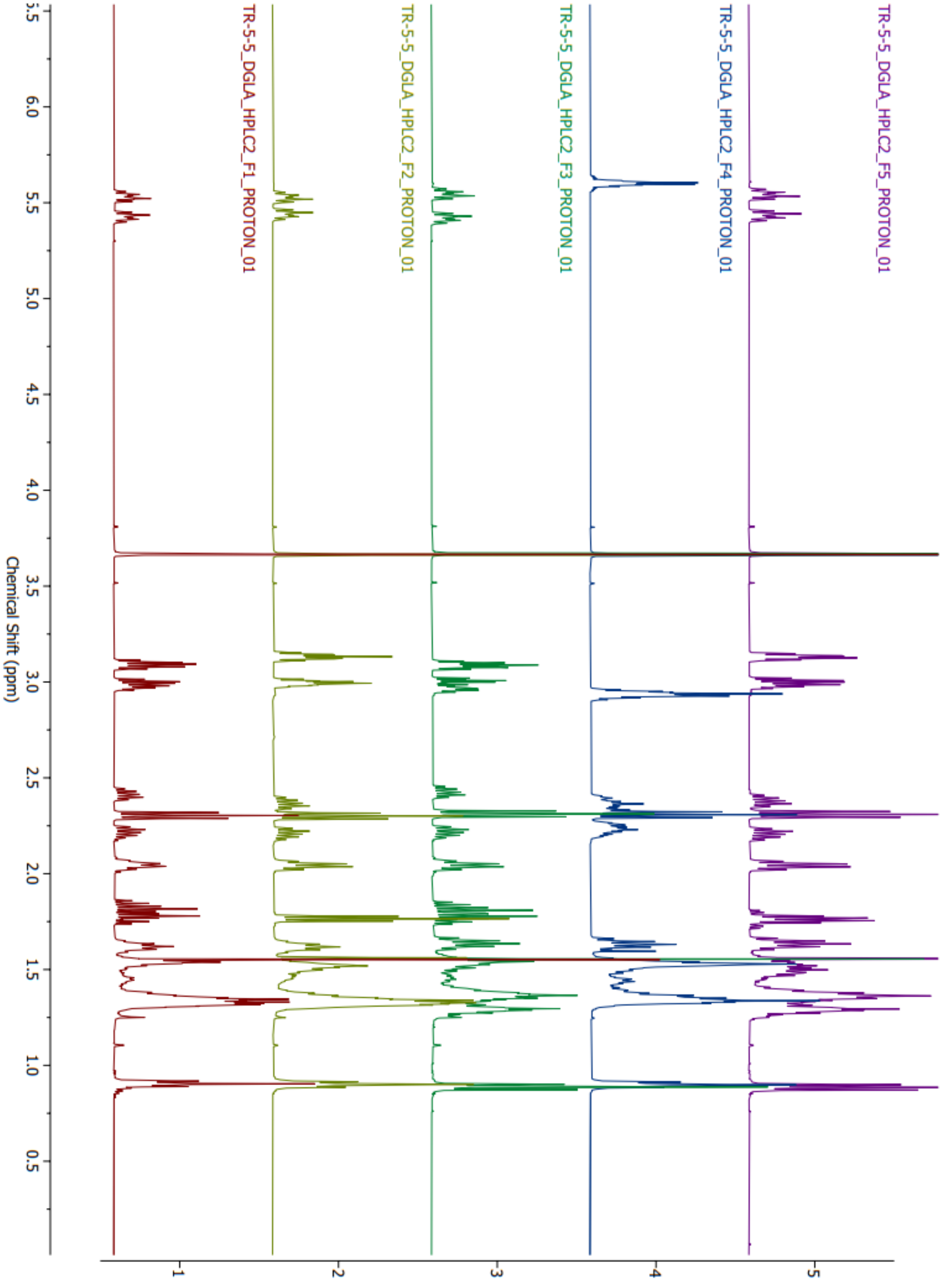
TR-5-5_DGLA_HPLC2_F5_PROTON_01
TR-5-5_DGLA_HPLC2_F4_PROTON_01
TR-5-5_DGLA_HPLC2_F3_PROTON_01
TR-5-5_DGLA_HPLC2_F2_PROTON_01
TR-5-5_DGLA_HPLC2_F1_PROTON_01
Chemical Shift (ppm)

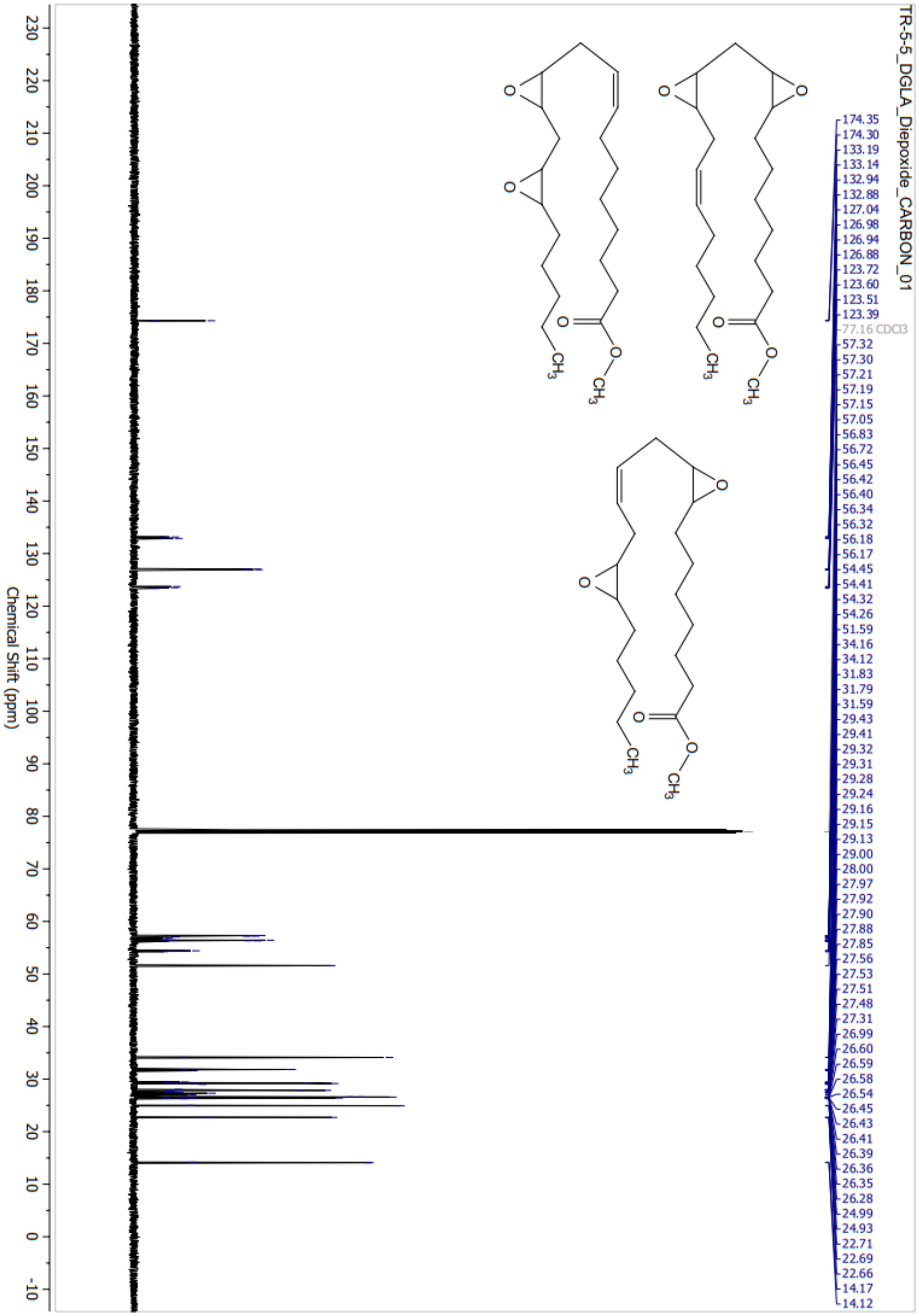
TR-5-5_DGLA_Diepoxide_CARBON_01
174.35
174.30
133.19
133.14
132.94
132.88
127.04
126.98
126.94
126.88
123.72
123.60
123.51
123.39
77.16 CDCl3
57.32
57.30
57.21
57.19
57.15
57.05
56.83
56.72
56.45
56.42
56.40
56.34
56.32
56.18
56.17
54.45
54.41
54.32
54.26
51.59
34.16
34.12
31.83
31.79
31.59
29.43
29.41
29.32
29.31
29.28
29.24
29.16
29.15
29.13
29.00
28.00
27.97
27.92
27.90
27.88
27.85
27.56
27.53
27.51
27.48
27.31
26.99
26.60
26.59
26.58
26.54
26.45
26.43
26.41
26.39
26.36
26.35
26.28
24.99
24.93
22.71
22.69
22.66
14.17
14.12
Chemical Shift (ppm)

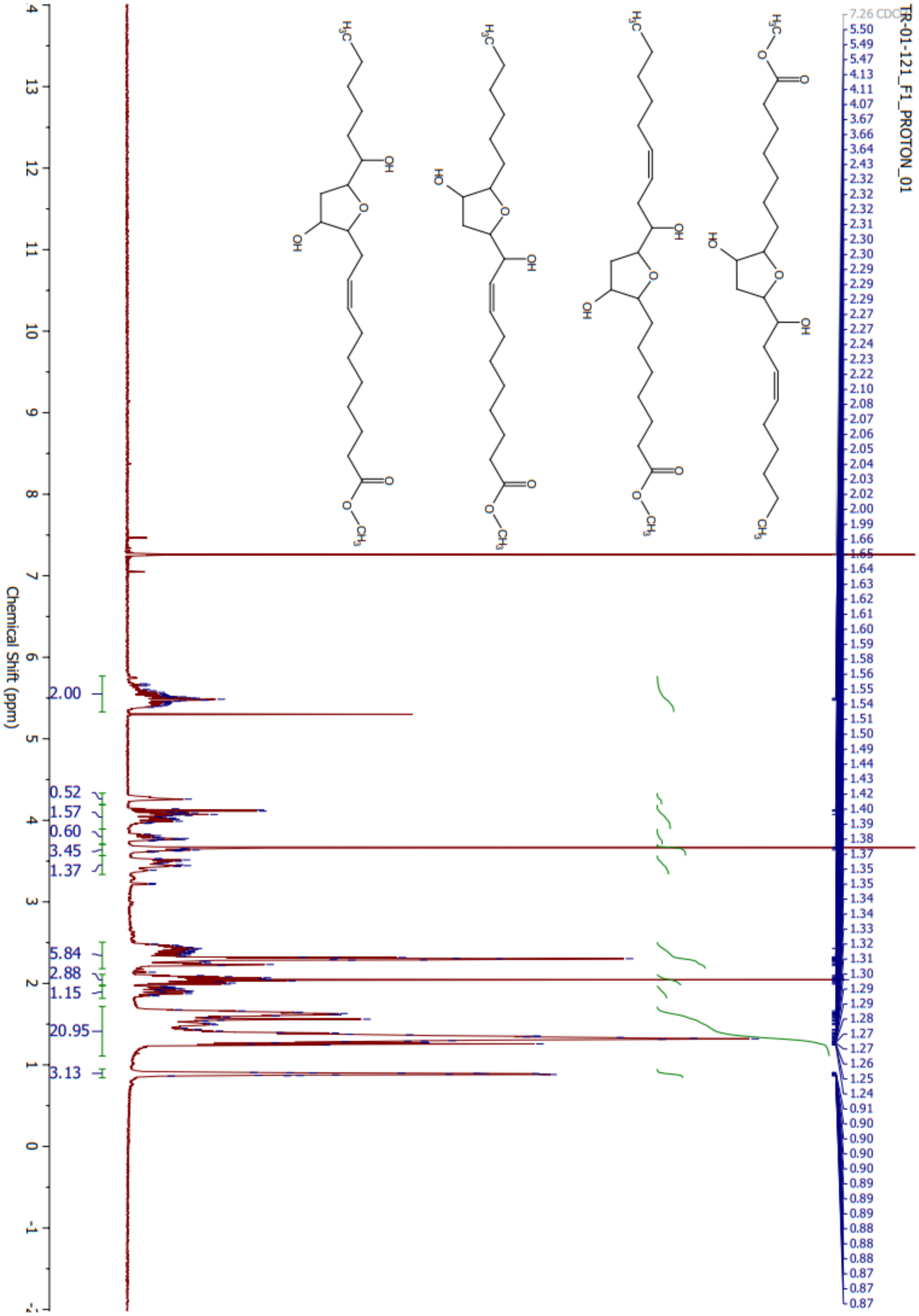
TR-01-121_F1_PROTON_01
Chemical Shift (ppm)
2.00
0.52
1.57
0.60
3.45
1.37
5.84
2.88
1.15
20.95
3.13

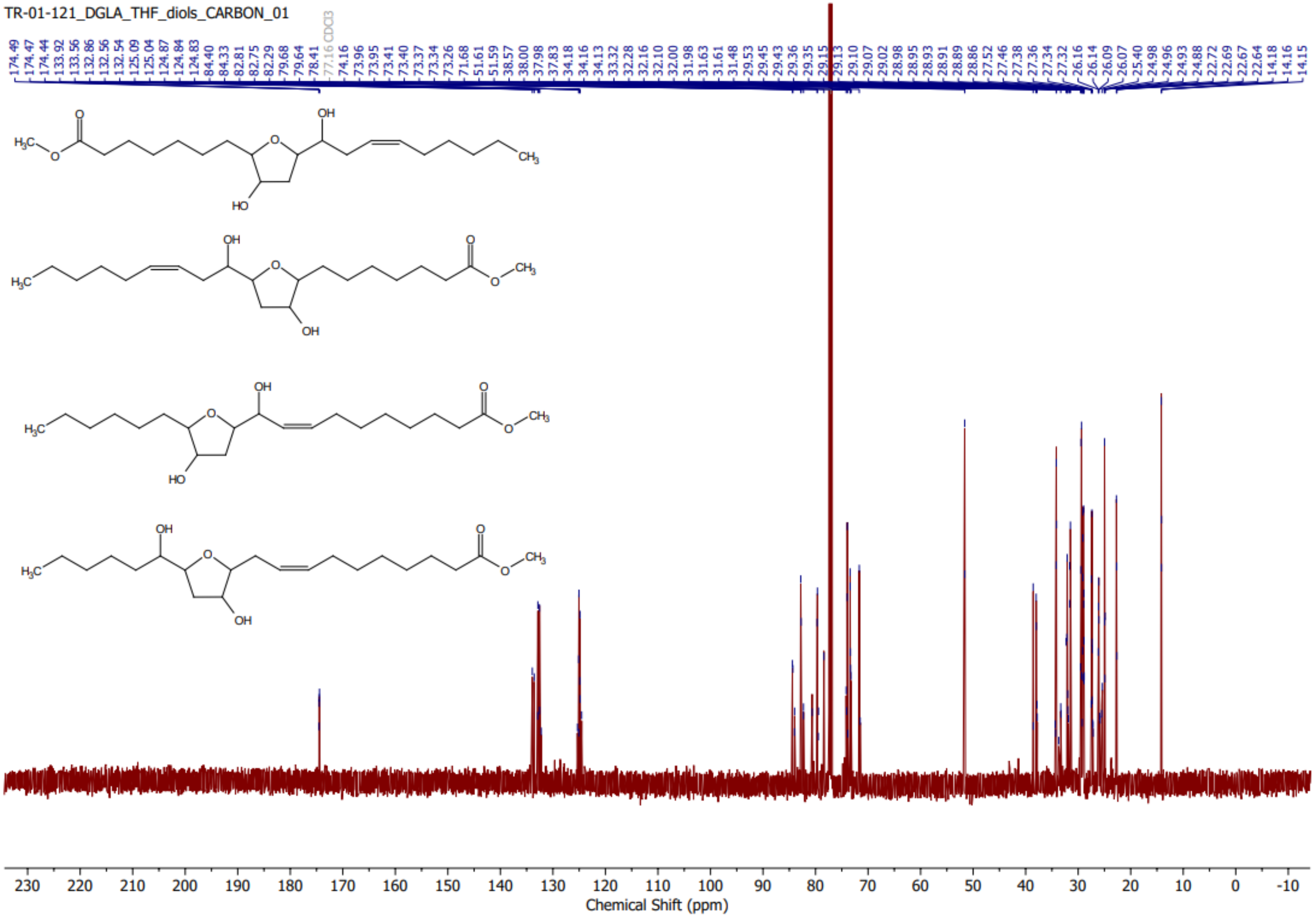
TR-01-121_DGLA_THF_diols_CARBON_01
174.49
174.47
174.44
133.92
133.56
132.86
132.56
132.54
125.09
125.04
124.87
124.84
124.83
84.40
84.33
82.81
82.75
82.29
79.68
79.64
78.41
77.16 CDCl3
74.16
73.96
73.95
73.41
73.40
73.37
73.34
73.26
71.68
51.61
51.59
38.57
38.00
37.98
37.83
34.18
34.16
34.13
33.32
32.28
32.16
32.10
32.00
31.98
31.63
31.61
31.48
29.53
29.45
29.43
29.36
29.35
29.15
29.13
29.10
29.07
29.02
28.98
28.95
28.93
28.91
28.89
28.86
27.52
27.46
27.38
27.36
27.34
27.32
26.16
26.14
26.09
26.07
25.40
24.98
24.96
24.93
24.88
22.72
22.69
22.67
22.64
14.18
14.16
14.15
Chemical Shift (ppm)

Date: Tue, Oct 20, 1959 5:00 PM
Data: DGLA diepox best run for quant

Sampling Int: 0.1 Seconds

Data:

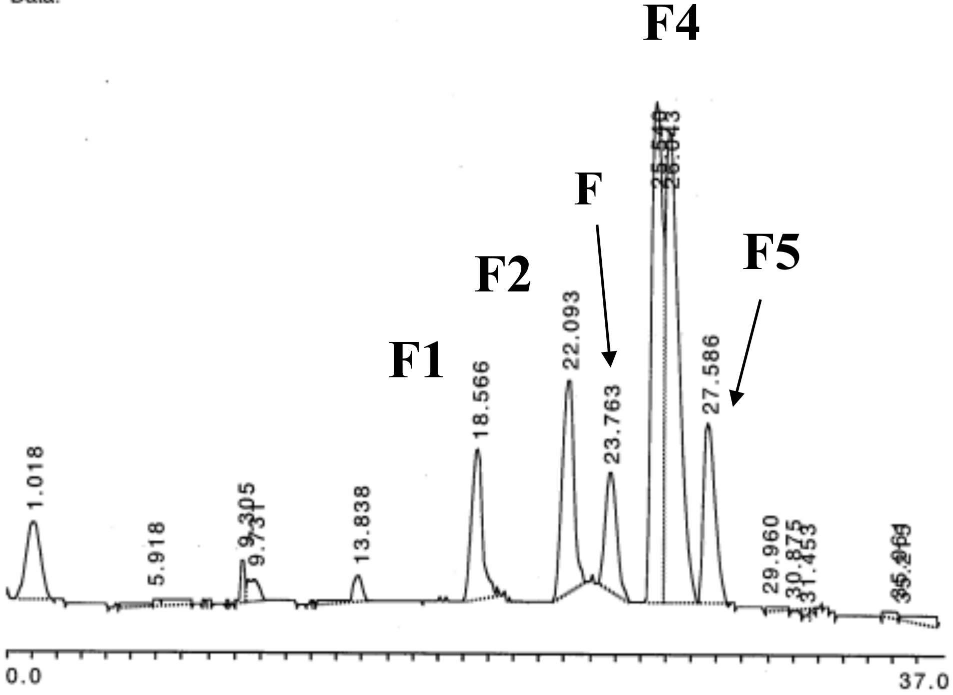

Analysis: Channel A

| Peak No. | Time | Type | Height(μV) | Area(μV-sec) | Area% |
|---|---|---|---|---|---|
| 1 | 1.018 | N | 8266 | 280257 | 4.972 |
| 2 | 5.918 | N | 740 | 82522 | 1.464 |
| | 7.770 | N | 486 | 7080 | 0.125 |
| | 8.693 | N1 | 466 | 2325 | 0.041 |
| | 8.868 | N2 | 69 | 1313 | 0.023 |
| 3 | 9.305 | N3 | 4674 | 39810 | 0.706 |
| 4 | 9.731 | N4 | 2537 | 53201 | 0.943 |
| | 10.088 | N5 | 475 | 425 | 0.007 |
| 5 | 13.838 | N | 2962 | 70951 | 1.258 |
| 6 | 18.566 | N | 15895 | 415424 | 7.370 |
| 7 | 22.093 | N | 22355 | 605169 | 10.736 |
| 8 | 23.763 | N | 12442 | 339999 | 6.032 |
| 9 | 25.540 | N1 | 52728 | 1345581 | 23.872 |
| 10 | 26.043 | N2 | 49795 | 1759054 | 31.207 |

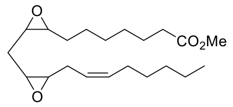

**1a**

***Identified as F1 & F3***

***Relative Ratios = 15%***

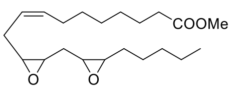

**1b**

***Identified as F2 & F5***

***Relative Ratios = 22%***

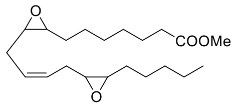

**1c**

***Identified as F4***

***Relative Ratios = 63%***

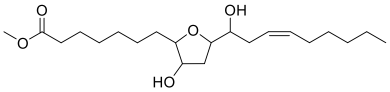

**2** *(+ structural & stereoisomers)*

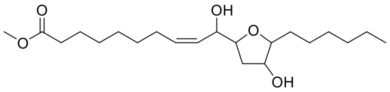

**3** *(+ structural & stereoisomers)*

*Relative ratio of* ***2:3*** *= 3:2*

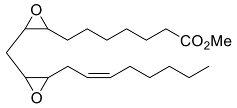

**1a**

***Identified as F1 & F3***

***Relative Ratios = 15%***

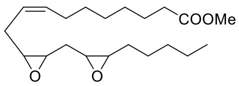

**1b**

***Identified as F2 & F5***

***Relative Ratios = 22%***

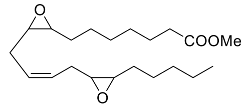

**1c**

***Identified as F4***

***Relative Ratios = 63%***

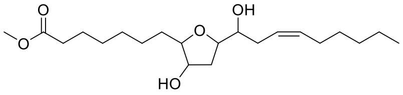

**2** *(+ structural & stereoisomers)*

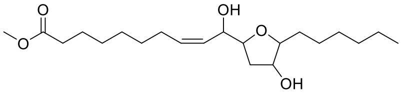

**3** *(+ structural & stereoisomers)*

*Relative ratio of* ***2:3*** *= 3:2*

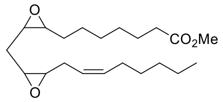

**1a**

***Identified as F1 & F3***

***Relative Ratios = 15%***

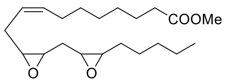

**1b**

***Identified as F2 & F5***

***Relative Ratios = 22%***

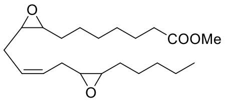

**1c**

***Identified as F4***

***Relative Ratios = 63%***

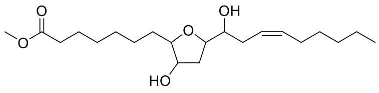

**2** *(+ structural & stereoisomers)*

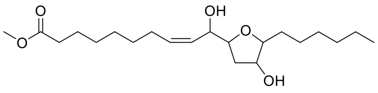

**3** *(+ structural & stereoisomers)*

*Relative ratio of* ***2:3*** *= 3:2*

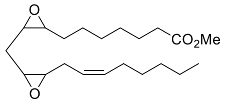

**1a**

***Identified as F1 & F3***

***Relative Ratios = 15%***

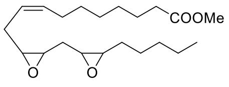

**1b**

***Identified as F2 & F5***

***Relative Ratios = 22%***

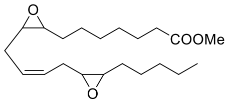

**1c**

***Identified as F4***

***Relative Ratios = 63%***

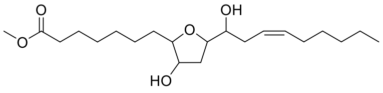

**2** *(+ structural & stereoisomers)*

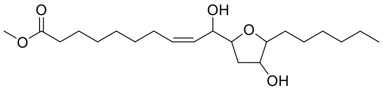

**3** *(+ structural & stereoisomers)*

*Relative ratio of* ***2:3*** *= 3:2*

Date: Tue, Oct 20, 1959 5:00 PM
Data: DGLA diepox best run for quant

Analysis: Channel A

| Peak No. | Time | Type | Height(μV) | Area(μV-sec) | Area% |
|---|---|---|---|---|---|
| 11 | 27.586 | N3 | 19023 | 506174 | 8.980 |
| 12 | 29.960 | N1 | 494 | 20073 | 0.356 |
| 13 | 30.875 | N2 | 189 | 10108 | 0.179 |
| 14 | 31.453 | N3 | 813 | 16459 | 0.292 |
| | 31.698 | N4 | 377 | 5591 | 0.099 |
| | 31.915 | N5 | 60 | 107 | 0.001 |
| 15 | 35.061 | N1 | 723 | 19789 | 0.351 |
| 16 | 35.215 | N2 | 297 | 55154 | 0.978 |
| Total Area | | | | 5636566 | 99.992 |

```
File       :C:\msdchem\1\data\Tommy\EI-AJ.M\TR-5-5 HPLC2 F3.D
Operator   : Jesus
Acquired   : 24 May 2022  13:44     using AcqMethod AJ_EI.M
Instrument :   5975MSD
Sample Name:
Misc Info  :
Vial Number: 0
```

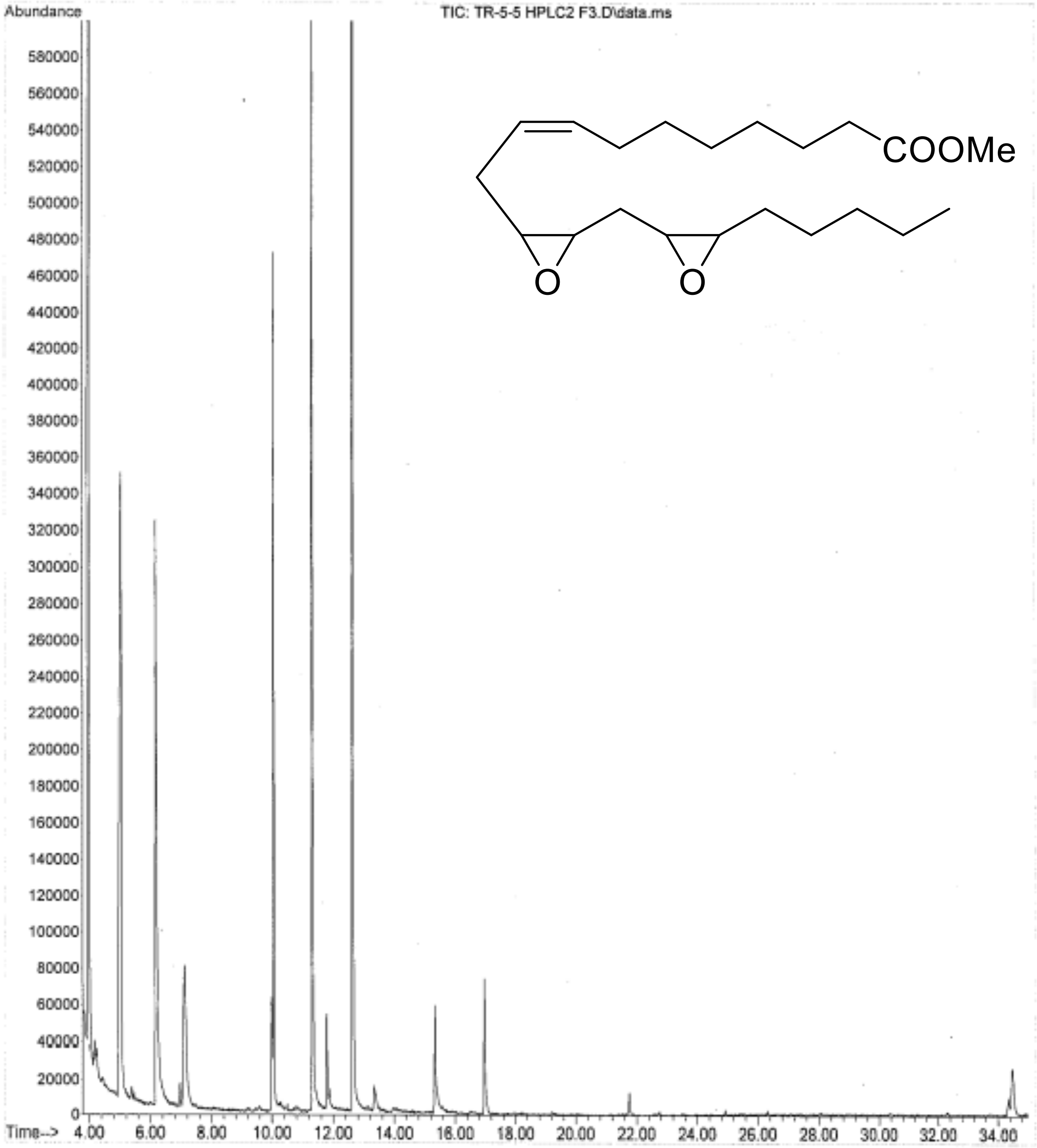

```
File        :C:\msdchem\1\data\Tommy\EI-AJ.M\TR-5-5 HPLC2 F5.D
Operator    : Jesus
Acquired    : 24 May 2022   14:34      using AcqMethod AJ_EI.M
Instrument :    5975MSD
Sample Name:
Misc Info   :
Vial Number: 0
```

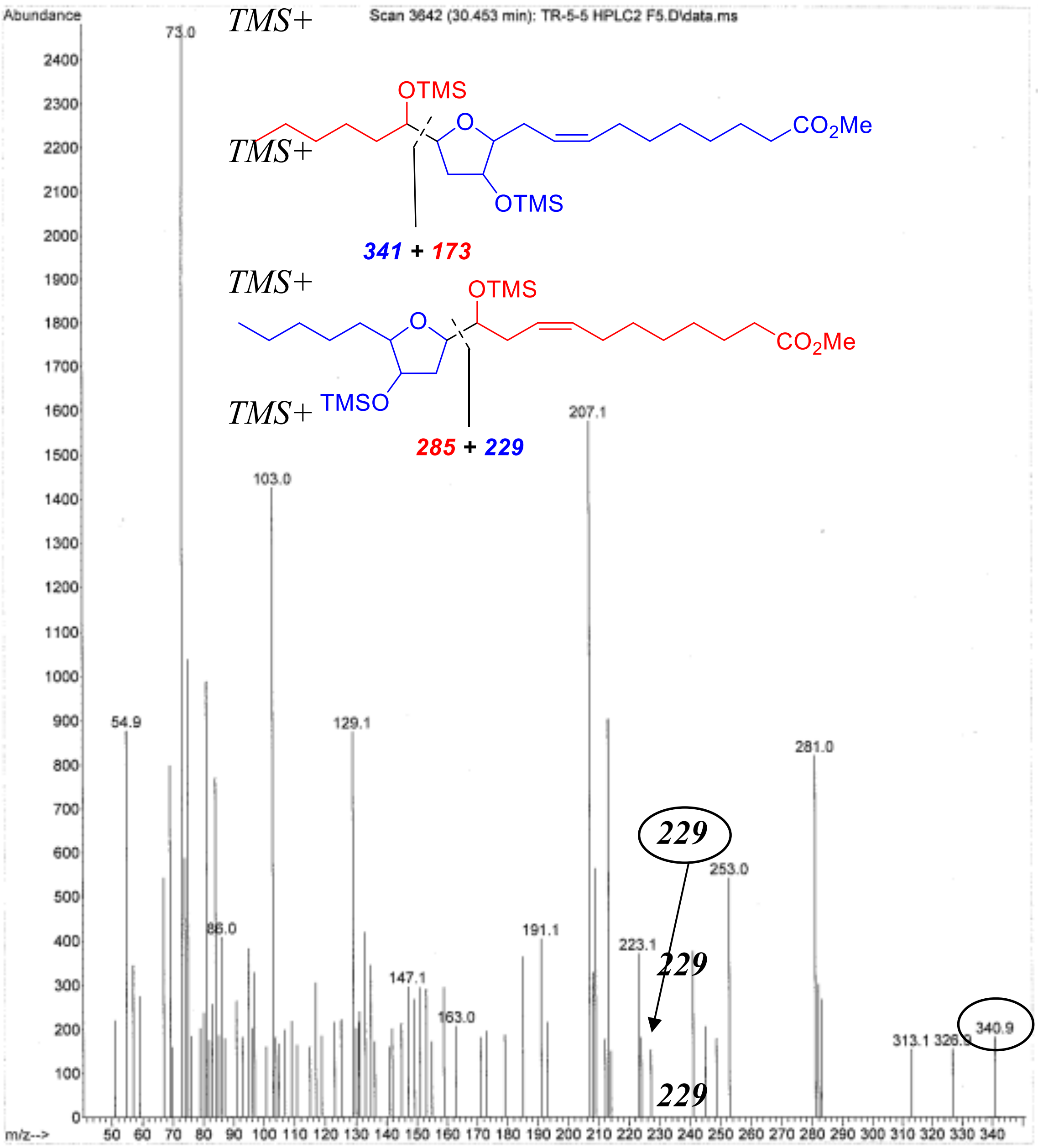

229

```
File        :C:\msdchem\1\data\Tommy\EI-AJ.M\TR-5-5 HPLC2 F5.D
Operator    : Jesus
Acquired    : 24 May 2022   14:34     using AcqMethod AJ_EI.M
Instrument  :    5975MSD
Sample Name:
Misc Info   :
Vial Number: 0
```

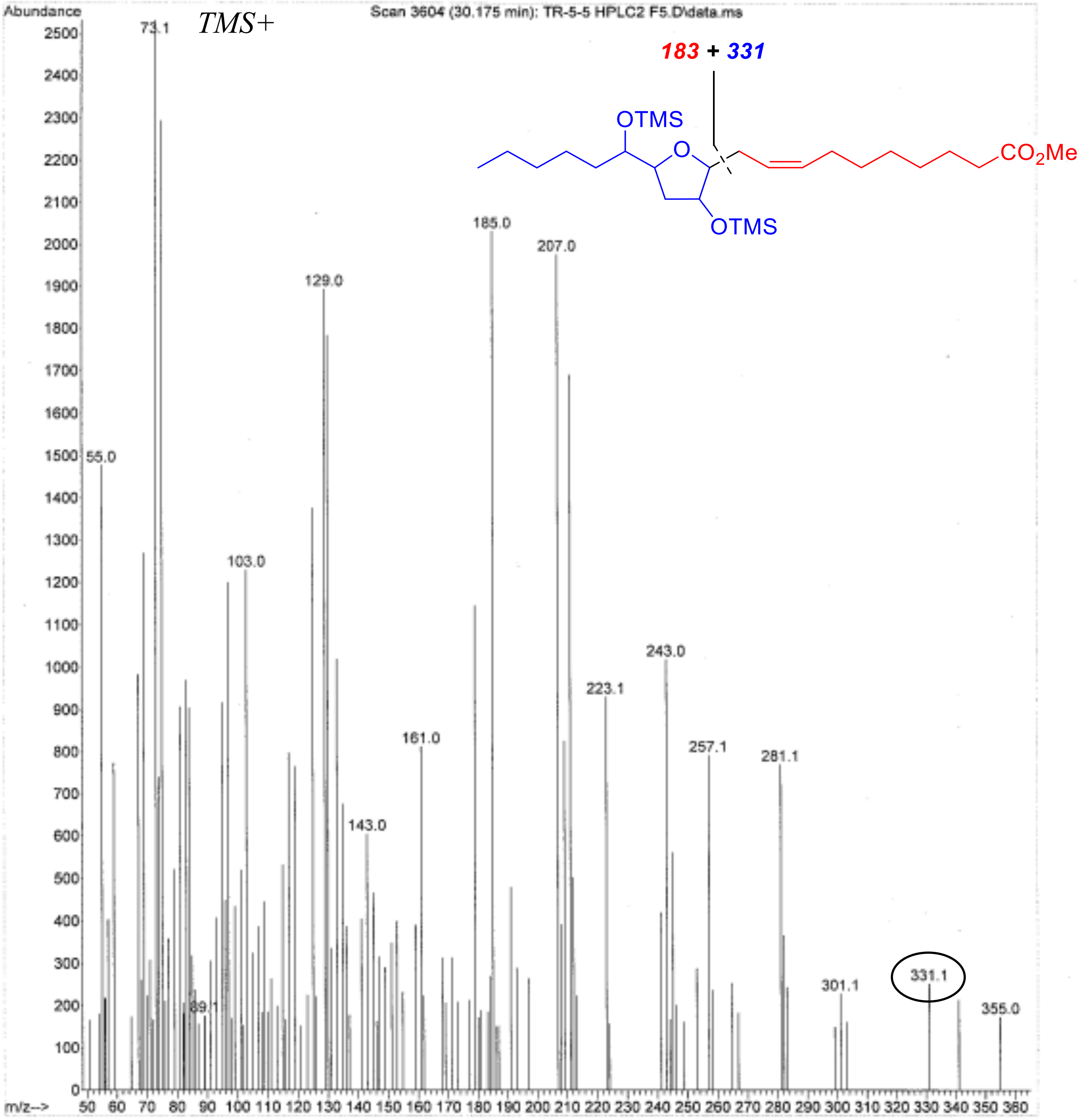

```
File       :C:\msdchem\1\data\Tommy\EI-AJ.M\TR-5-5 HPLC2 F5.D
Operator   : Jesus
Acquired   : 24 May 2022  14:34     using AcqMethod AJ_EI.M
Instrument :    5975MSD
Sample Name:
Misc Info  :
Vial Number: 0
```

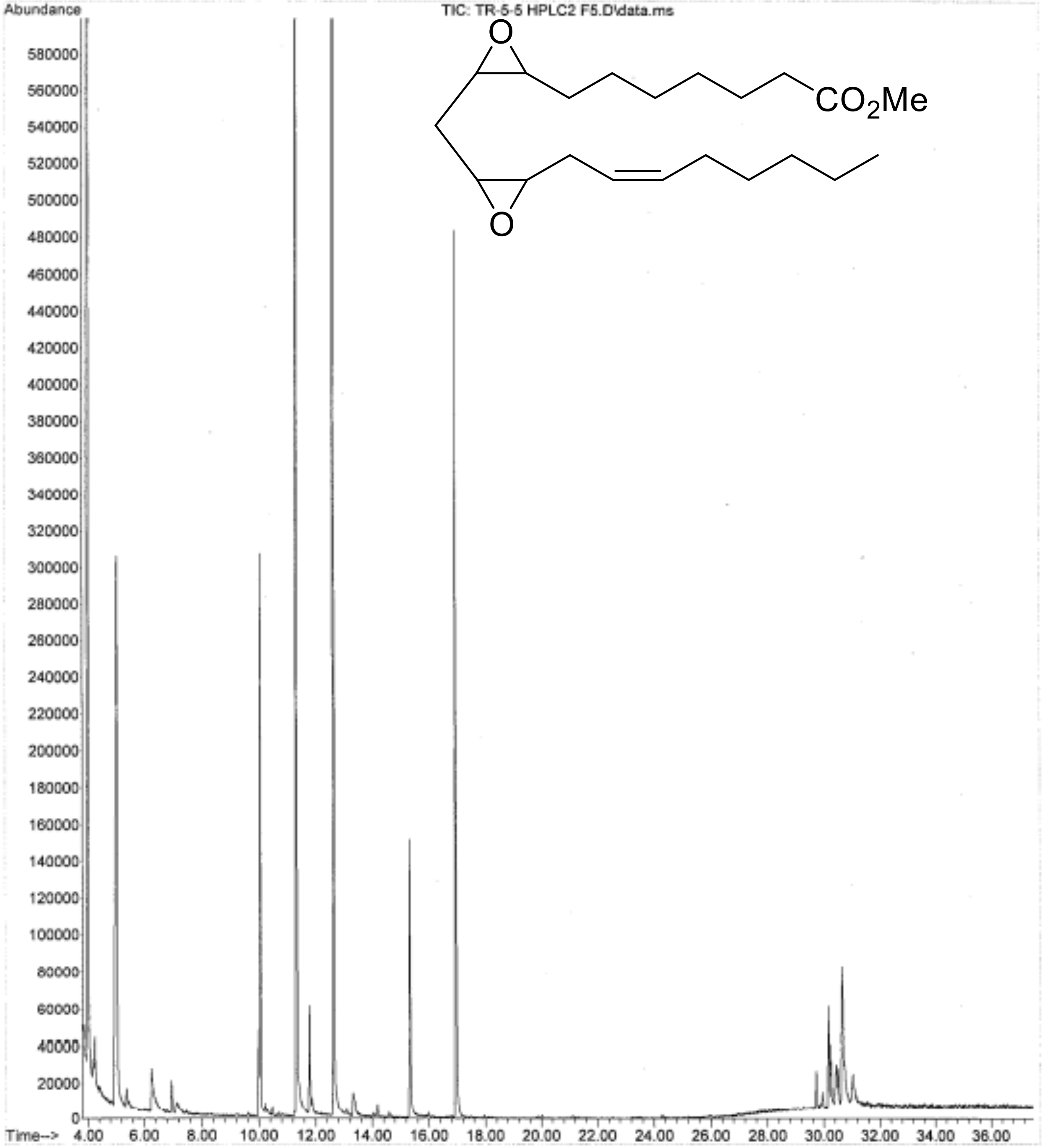

```
File        :C:\msdchem\1\data\Tommy\EI-AJ.M\TR-5-5 HPLC2 F3.D
Operator    : Jesus
Acquired    : 24 May 2022   13:44      using AcqMethod AJ_EI.M
Instrument  :    5975MSD
Sample Name:
Misc Info   :
Vial Number: 0
```

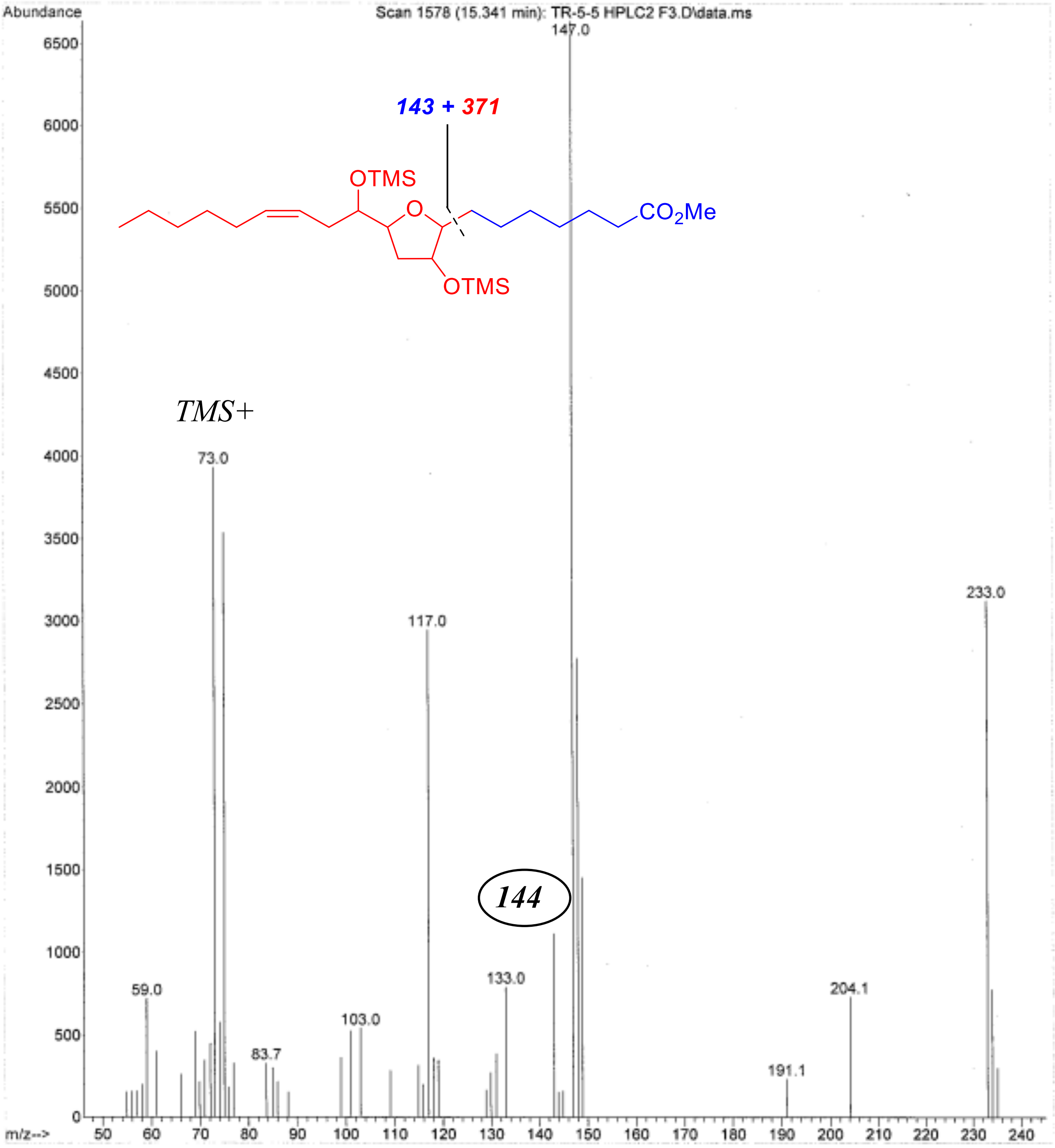

```
File        :C:\msdchem\1\data\Tommy\EI-AJ.M\TR-5-5 HPLC2 F3.D
Operator    : Jesus
Acquired    : 24 May 2022   13:44      using AcqMethod AJ_EI.M
Instrument :    5975MSD
Sample Name:
Misc Info  :
Vial Number: 0
```

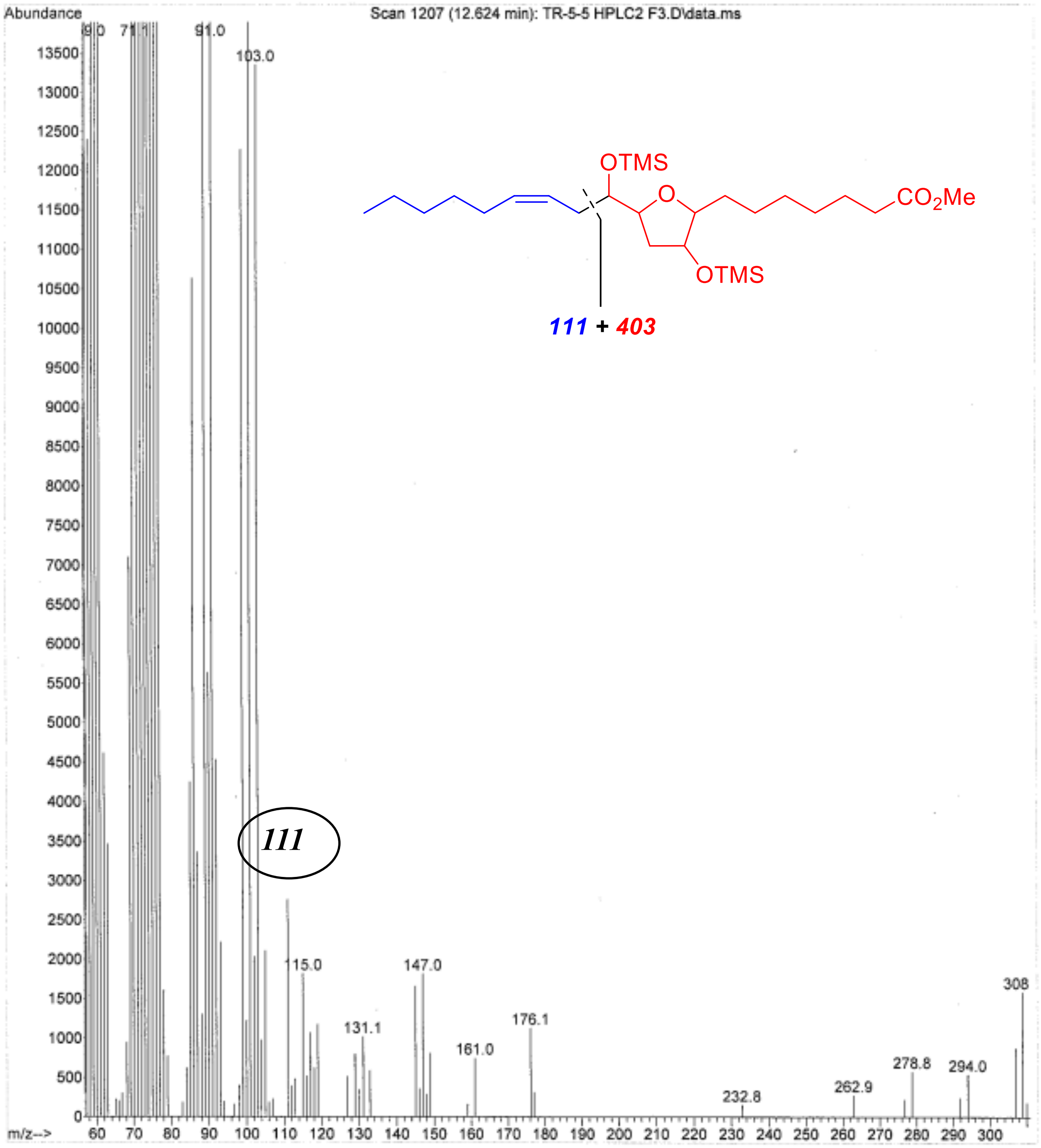

1. Fay, D. (2006). Genetic mapping and manipulation: chapter 1--Introduction and basics. WormBook, 1–12.

2. Vladis, N.A., Fischer, K.E., Digalaki, E., Marcu, D.-C., Bimpos, M.N., Greer, P., Ayres, A., Li, Q., and Busch, K.E. (2019). Gap junctions in the C. elegans nervous system regulate ageing and lifespan. bioRxiv, 657817. 10.1101/657817.

3. Stiernagle, T. (2006). Maintenance of C. elegans. WormBook, 1–11.

4. Tucci, M.L., Harrington, A.J., Caldwell, G.A., and Caldwell, K.A. (2011). Modeling dopamine neuron degeneration in Caenorhabditis elegans. Methods Mol. Biol. *793*, 129–148.

5. Kim, I.-H., Nishi, K., Tsai, H.-J., Bradford, T., Koda, Y., Watanabe, T., Morisseau, C., Blanchfield, J., Toth, I., and Hammock, B.D. (2007). Design of bioavailable derivatives of 12-(3-adamantan-1-yl-ureido)dodecanoic acid, a potent inhibitor of the soluble epoxide hydrolase. Bioorg. Med. Chem. *15*, 312–323.
